# Targeting GFPT2 to reinvigorate immunotherapy in EGFR-mutated NSCLC

**DOI:** 10.1101/2024.03.01.582888

**Authors:** Luyao Ao, Wenjing Jia, Qixing Gong, Jiawen Cui, Jun Wang, Ying Yu, Chenghao Fu, Haobin Li, Jia Wei, Ruiqi Wang, Feiyi Wang, Xin Shang, Yantong Li, Shencun Fang, Guangji Wang, Fang Zhou, Jiali Liu

## Abstract

In the evolving field of cancer immunotherapy, EGFR-mutated NSCLC presents a significant challenge, demonstrating marked innate resistance to established treatments. An effective method to counter this resistance remains elusive. Through comprehensive genetic and pharmacological analyses across various models, we have identified glutamine fructose-6-phosphate transaminase 2 (GFPT2) as a key facilitator of immune evasion in EGFR-mutated NSCLC. Mechanistically, under EGFR mutation condition, GFPT2 expression, which is typically low in normal tissues, is highly induced via EGFR/IRE1α/Xbp1s signaling axis, leading to a significant increase in intracellular UDP-GlcNAc and consequently, an altered N-glycosylation profile. GFPT2 escalates the expression and glycosylation of PD-L1, PVR and CD276, bolstering their interactions with CD8^+^T cells, and amplifies CD73 glycosylation, thereby intensifying adenosine-mediated CD8^+^T cells suppression. These actions collectively reduce tumor cell vulnerability to CD8^+^T cell-mediated death. Moreover, GFPT2 regulates EGFR glycosylation, which consequentially modulates the EGFR-dependent secretion of CXCL10 and VEGF, thus impeding CD8^+^T cell infiltration within tumors. We further identified a GFPT2 isoform-specific inhibitor that potentiates PD-1 blockade therapy beyond that of existing strategy, corroborated by results in xenografts and patient-derived organoids. Together, these findings illuminate the promising therapeutic potential of GFPT2 as a metabolic checkpoint, offering an innovative approach to invigorate immunotherapy in NSCLC with EGFR mutations.

## Introduction

Lung carcinoma leads globally in cancer mortality and ranks second in morbidity. Non-small cell lung cancer (NSCLC) is the predominant subtype and epidermal growth factor receptor (EGFR) is the most common driver mutation^1^. While immunotherapy benefits various cancers, none have shown clinical advantage in subgroups of patients with NSCLC harboring EGFR mutations^2^. Addressing the gap in immunotherapy for EGFR-mutated NSCLC, is a pressing issue, considering the inevitable resistance of current EGFR tyrosine kinase inhibitors (TKIs) therapy and the ensuing limited long-lasting clinical benefit^3^.

The compromised antitumor immune response observed in EGFR-mutated NSCLC is a consequence of a multifactorial suppressive tumor immune microenvironment (TIME). This environment obstructs the recruitment and activation of immune cells through diminished chemokine secretion, excessed production of negative immune modulators, and elevated expression of immunosuppressive molecules, necessitating interventions at multiple stages simultaneously to reinvigorate immunotherapy in EGFR-mutated NSCLC^2^. Given the pivotal role of tumor cell-driven EGFR mutations in shaping this immunosuppressive microenvironment, combination therapies involving TKIs and immunotherapy have been proposed. However, the therapeutic benefit is limited to a transient window and safety risk^4–6^. Post-TKI acquired resistance, the immunosuppressive microenvironment intensifies, posing an even greater challenge^4,6^. To date, revitalizing immunotherapy in EGFR-mutated NSCLC remains a significant challenge.

In the tumor microenvironment, there is a pronounced metabolic-immune interaction, with endogenous metabolism occasionally acting as an upstream regulator of immune responses. Metabolic immune checkpoints, functioning as key mediators of tumor metabolic variations, orchestrate immune control through either metabolite competition or metabolite-driven biological processes^7–10^. These metabolic checkpoints, are gaining prominence as potential therapeutic targets for modulating tumor immunity. In NSCLC, a variety of genetic mutations induce unique endogenous metabolic phenotypes^11,12^; however, the consequences of these mutation-induced metabolic alterations on the tumor immune phenotype are not fully understood. Elucidating the nexus between EGFR mutation-driven metabolic reprogramming and the resultant complex suppressive TIME might unveil novel targets for effective immunomodulatory cancer treatments.

Our research reveals that GFPT2 expression is inducibly expressed by in the context of EGFR-mutant, leading to significant accumulation of UDP-GlcNAc and subsequent hyper-glycosylation within tumor cells. GFPT2, by modulating protein glycosylation, plays a pivotal role in immune evasion. This discovery positions GFPT2 as a novel ‘metabolic immune checkpoint’, providing a fresh avenue to enhance the efficacy of immunotherapy in EGFR-mutated NSCLC.

## Results

### Metabolic profile in the context of EGFR-mutant

An unbiased metabolomic analysis has disclosed a significant metabolic divergence between NSCLC with EGFR mutations and those with wild-type EGFR. This divergence is characterized by a marked increase in uridine 5’-diphospho-N-acetylglucosamine (UDP-GlcNAc) in EGFR-mutated NSCLC ***(Fig. 1a)***. This strong association between UDP-GlcNAc accrual and EGFR mutations was further substantiated by consistent metabolic alterations in BaF3 cells following the induction of various oncogenic EGFR mutations ***(Extended Data Fig. 1a-1d, Fig. 1b)***.

**Fig.1.**
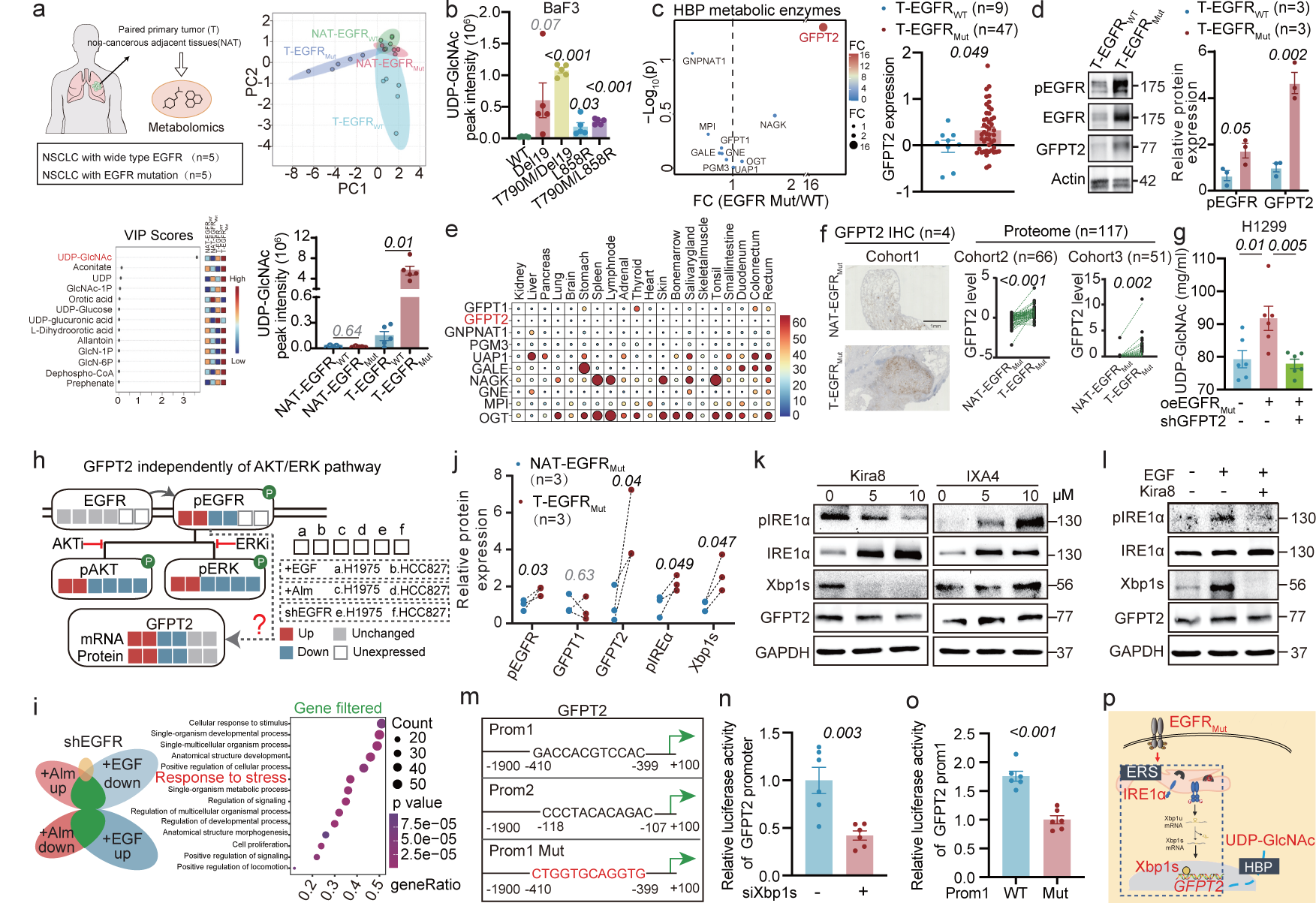
Induction of GFPT2 expression in the context of EGFR mutations. **a**, Metabolomic analysis contrasts tumor (T) and non-tumor adjacent tissue (NAT) in NSCLC patients with wild-type EGFR (n=5) and those with EGFR mutations (n=5). Principal component analysis (PCA) was utilized to differentiate metabolite profiles between these groups, and the variable importance in projection (VIP) scores identified the top 15 significant metabolites. Among these, UDP-GlcNAc levels were specifically evaluated. **b**, HPLC-QTOF/MS quantification of UDP-GlcNAc in BaF3 cells expressing various EGFR_Mut_ oncogenes (n=5) highlights the metabolic impact of EGFR mutations. **c**, Differential expression of HBP metabolic enzymes was analyzed in proteomic data from 9 NSCLC patients with wild-type EGFR and 47 NSCLC patients with mutated EGFR. The data were sourced from the Database of Genotypes and Phenotypes (ph-s001954.v1.p1). **d**, Immunoblot analyses showing the levels of phosphorylated EGFR (pEGFR), total EGFR, and GFPT2 in tumor samples from NSCLC patients with wild-type EGFR (n=3) and those harboring EGFR mutations (n=3), using Actin as a loading control. **e**, Heatmap illustrating the expression levels of hexosamine biosynthetic pathway (HBP) metabolic enzymes across major organs, utilizing data from the proteomic database (PRJNA183192). **f**, Comparative analysis of GFPT2 expression in T and matched NAT from NSCLC patients with EGFR mutations. Immunohistochemistry (IHC) samples were obtained from Jiangsu Province Hospital. Proteomic data were derived from two independent databases (IPX0001804000 and PRJNA183192). **g**, HPLC-QTRAP/MS analysis of UDP-GlcNAc levels in H1299 (n=6). **h**, Schematic of GFPT2 mRNA/protein expression changes and downstream EGFR pathway modifications post EGF/Almonertinib (Alm) treatment or EGFR knockdown. **i**, RNA-seq in H1975 cells post-treatment: gene filtering strategy (left) and Gene Ontology enrichment (right). **j**, Western blot quantification of pEGFR, pIREα, Xbp1s, and GFPT2 in T and NAT from NSCLC patients harboring EGFR mutations (n=3). **k-l**, Blot representations of pIREα, IRE, Xbp1s, and GFPT2 expressions in H1975 cells treated with IREα inhibitor (Kira8)/agonist (IXA4) or EGF, including GAPDH as the loading control. **m**, Predicted Xbp1s binding sequences on GFPT2 promoter by Jasper, featuring promoter 1 Mut as the reverse complement. **n**, Dual-Luciferase assay in H1299 cells for GFPT2 promoter activity with siRNA interventions (n=6). **o**, Dual-Luciferase assay in H1299 cells for GFPT2 promoter 1 mutation (n=6). **p**, Schematic of GFPT2 upregulation mechanism in EGFR_Mut_ NSCLC. The immunoblots in **j** and **k** are representative of two independent experiments. Data are mean ± s.e.m. except in **f, j**, Two-tailed paired Student’s t-test. **c**, **d**, **n**, **o**, Two-tailed unpaired Student’s t-test. **a** (UDP-GlcNAc level), **b**, **g**, One-way ANOVA.

A fraction (2 - 5 %) of glucose uptake is channeled into hexosamine biosynthetic pathway (HBP) to generate UDP-GlcNAc^13^. Within the cadre of metabolic enzymes integral to HBP, glutamine fructose-6-phosphate transaminase 2 (GFPT2) is singularly conspicuous. Comparative analysis of proteomic data from 9 EGFR wild-type and 47 EGFR-mutated NSCLC cases revealed significant overexpression of GFPT2 specifically in EGFR-mutated NSCLC, with no notable changes in other HBP metabolic enzymes ***(Fig.1c)***, which was further validated by immunoblotting assay with six independent clinical samples ***(Fig.1d)***. Consistent with these clinical observations, GFPT2 expression was induced in various cell lines following the introduction of EGFR mutations ***(Extended Data Fig. 1e)***. Furthermore, GFPT2 exhibits a distinct expression, being relatively subdued levels in normal tissues juxtaposed against marked upregulation in NSCLC neoplasms ***(Fig.1e-1f, Extended Data Fig.1f-1g)***, and its high expression correlates with poor NSCLC outcomes ***(Extended Data Fig.1h-1i)***, identifying it as mutation-induced protein in the EGFR-driven tumorigenesis. GFPT2 shares a similar enzymatic function with its isoform GFPT1 ***(Extended Data Fig.1j)***, but is notably less affected by negative feedback inhibition from HBP metabolites, suggesting it potentially facilitates sustained HBP activation and subsequent aberrant biosynthesis of UDP-GlcNAc^14,15^***(Extended Data Fig.1k)***. Upon introducing EGFR mutation, the elevated UDP-GlcNAc levels abrogated with GFPT2 silencing, indicating GFPT2’s predominant role in the aberrant UDP-GlcNAc accrual mediated by the EGFR mutation ***(Fig.1g)***. Furthermore, GFPT2-dependent UDP-GlcNAc accrual is also observed in EGFR TKI-resistant NSCLC subtypes ***(Extended Data Fig.1l-1p)***.

### GFPT2 induction via EGFR/IREα/Xbp1 axis

Next, we delved deeper into the molecular underpinnings governing the augmented GFPT2 levels in the context of EGFR mutation and intriguingly found that GFPT2 regulation does not follow the classical downstream pathways of hyperactivated EGFR ***(Fig.1h)***. GFPT2’s transcriptional and translational expressions can be modulated by EGFR inhibition with Almonertinib (Alm) and EGFR activation via EGF, yet remain unchanged by EGFR ablation ***(Extended Data Fig.2a-2b).*** This suggests a unique regulatory mechanism for GFPT2, divergent from the well-characterized AKT and ERK signaling alterations^16^, as evidenced by the inability of AKT or ERK inhibitors to modulate GFPT2 expression ***(Extended Data Fig.2c-2d)***, pointing to an unidentified pathway modulating GFPT2 in response to EGFR hyperactivation.

TraCe-seq analysis revealed endoplasmic reticulum stress (ERS) alterations in the anti-EGFR treatment cohort, which was not observed in cells undergoing EGFR-targeted degradation^17^, a finding echoed by our independent unbiased bulk-seq analysis ***(Fig.1i, Extended Data Fig.2e)***. Within the three distinct ERS pathways, we identified a nucleotide sequence in the GFPT2 promoter that resembles the canonical Xbp1s binding site, a spliced variant of Xbp1 mRNA produced by phosphorylation-activated IRE1α^18^, which is more prevalent in various tumors than in normal tissues^19^ ***(Extended Data Fig.2f)***. A paired analysis of tumor versus adjacent non-tumor tissues showed that EGFR activation and GFPT2 overexpression in tumors are accompanied by increased IRE1α phosphorylation and Xbp1s expression ***(Fig.1j, Extended Data Fig.2g)***, suggesting the IRE1α/Xbp1 axis as a potential mediator of GFPT2 upregulation in EGFR-mutated NSCLC. The influence of EGFR on the IRE1α/Xbp1 pathway has been confirmed in both cell culture and xenograft models using H1975 and HCC827 NSCLC cell lines, which possess distinct EGFR mutations ***(Extended Data Fig.2h-2j)***. The role of IRE1α/Xbp1 in regulating GFPT2 expression was further substantiated by utilizing the IRE1α-specific inhibitor Kira8 and the agonist IXA4 ***(Fig.1k)***. Importantly, Kira8 effectively counteracted EGF-induced GFPT2 upregulation, supporting the notion that EGFR’s regulatory effect on GFPT2 primarily occurs through the IRE1α/Xbp1 signaling pathway ***(Fig.1l)***. In a similar vein, IRE1α/Xbp1 activation and subsequent GFPT2 expression enhancement were observed and were even more pronounced in TKI-resistant NSCLC subtypes ***(Extended Data Fig.2k)***. To determine whether Xbp1s orchestrates direct transcriptional modulation of GFPT2, we engineered a recombinant luciferase reporter construct encompassing the whole promoter region (−1900 to 100 bp) of human GFPT2. RNA interference-mediated depletion of Xbp1s yielded a marked decrease in luciferase enzymatic activity ***(Fig.1m-1n)***. Chromatin Immunoprecipitation (ChIP) assays showed Xbp1s enrichment in the presumed GFPT2 promoter binding site at −410 to −399 bp, affirming direct interaction ***(Extended Data Fig.2l)***, which was substantiated by luciferase assays using a mutated promoter variant, confirming the specificity of the binding ***(Fig.1o)***.

Synthetically collating our findings, GFPT2, a protein with inducible expression, is upregulated via the EGFR/IRE1α/Xbp1 signaling pathway, which is critical for the accumulation of UDP-GlcNAc in EGFR-mutated NSCLC (***Fig.1p***).

### Genetic disruption of tumoral GFPT2 reinvigorates anti-tumor immunity

GFPT2 is reported to be closely associated with the growth of tumor cells in KRAS/LKB1 co-mutant NSCLC due to their higher reliance on HBP pathway for survival^12^, yet this association is not observed in EGFR-mutated NSCLC cells. Genetic disruption of GFPT2 elicited no conspicuous alternations in the H1975 proliferative phenotype, as evinced by consistent metrics of cell viability, proliferation, and colony establishment ***(Extended Data Fig.3a-3d)***. Furthermore, although Azaserine (AZA), a non-selective inhibitor of GFPT1 and GFPT2, significantly inhibited the HBP pathway, it did not affect the growth of H1975 cells, both *in vitro* and in nude mouse xenograft models ***(Extended Data Fig.3e-3h)*.**

Remarkably, introducing an immune context engendered distinct phenotypic outcomes of GFPT2 knockdown in tumor cells. Within the immunodeficient NOD.Cg-PrkdscidIl2rgtm1Vst/Vs (NPG) cohort, shNC and shGFPT2 H1975 mirrored each other in growth dynamics ***(Fig.2a, Extended Data Fig.3i-3j)***. In a stark divergence, the humanized mice, conceived via the engraftment of human CD34^+^ hematopoietic precursors into NPG mice to induce human innate and adaptive immunological development (HSPC-NPG)^20^, displayed a significant reduction in tumor burden in shGFPT2 cells ***(Fig.2a, Extended Data Fig.3i-3j)***. Mirroring observations from Osimertinib-sensitive cells, changes in Osimertinib-resistant cells post-GFPT2 disruption were uniquely observed ***(Fig.2b)***. To circumvent potential false positives due to the dearth of lymph node structures and germinal centers in HSPC-NPG mice^20^, we utilized a murine NSCLC cell line with EGFR T790M/L858R mutation (EGFR-mutated LLC) in immunocompetent C57BL/6N mice, providing a more comprehensive immune context. In these mice, GFPT2-knockdown tumor cells grew significantly slower than controls. This growth suppression was notably reversed by FTY720, an immunosuppressant blocking T cell lymph node egress, underscoring the reliance of GFPT2’s tumor-suppressive function on the presence of immune cells within the tumor microenvironment ***(Fig.2c, Extended Data Fig.3k)***.

**Fig.2.**
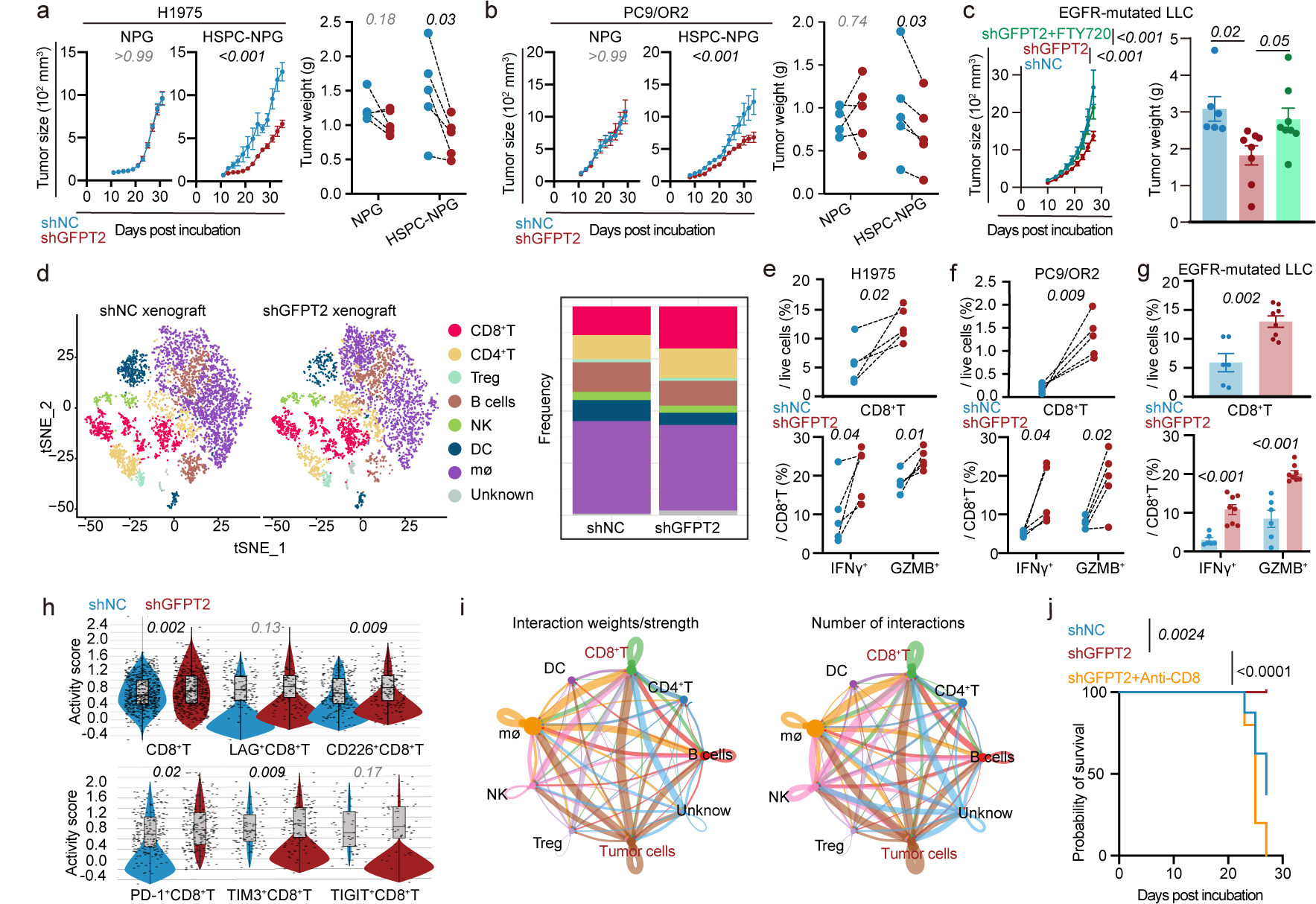
Assessing GFPT2 intervention effects on tumor progression and the tumor immune microenvironment. **a**, **b**, Tumor size (left) and tumor weight (right) in xenografts derived from H1975 or PC9/OR2 cells with GFPT2 silenced, transplanted into two flanks of NPG mice (immunodeficient) or HSPC-NPG mice (humanized immune-reconstituted), were evaluated (n=5 each group). **c**, Tumor size (left) and weight (right) in xenografts from EGFR-mutated LLC cells, treated with shNC (n=9), shGFPT2 (n=11), and shGFPT2+FTY720 (n=8), post transplantation into C57BL/6N mice. For single-cell sequencing, three mice from shNC and shGFPT2 groups were chosen, without recording tumor weights. **d**, Global representation of sorted CD45^+^ immune cells from EGFR-mutated LLC xenograft, including t-distributed stochastic neighbor embedding (t-SNE) plots denoting cell type and cell-type representation per genotype (stacked columns). DCs, dendritic cells; NK cells, natural killer; mø, Macrophages. **e-g**, Multicolor flow cytometry (FCM) validated CD8^+^T infiltration and activity changes in different mice tumors after GFPT2 knockdown (n=5 for H1975 and PC9/OR2 xenograft; n=6 for shNC, n=8 for shGFPT2 in EGFR-mutated LLC xenograft). **h**, scRNA-seq analysis of sorted CD8^+^T cells from EGFR-mutated LLC xenograft. Violin plots display early activation and effector/cytokine signaling in intratumoral CD8^+^T cells. Box plots indicate median and interquartile range. **i**, Cell-cell communication analysis in shNC and shGFPT2 EGFR-mutated LLC xenograft leveraging scRNA-seq data. **j**, Survival analysis of C57BL/6N mice with EGFR-mutated LLC xenograft post GFPT2 silenced, treated with IgG or anti-CD8. Data are mean ± s.e.m. except in **a**, **b**, **e**, **f**, Two-tailed paired Student’s t-test (**a**, **b**, tumor weight). **g h**, Two-tailed unpaired Student’s t-test. **c**, One-way ANOVA (tumor weight). **a–c**, Two-way ANOVA for tumor size. **j**, Mantel-Cox test.

Subsequent investigations into GFPT2 knockdown effects on the tumor immune microenvironment (TIME), utilizing single-cell RNA sequencing (scRNA-seq), identified an increased prevalence of CD8^+^T cells in tumors where GFPT2 was silenced ***(Fig.2d)***. This finding is particularly significant considering the usual depletion of these cells in patients with EGFR-mutated NSCLC ***(Extended Data Fig.4a)***. This increase was substantiated by flow cytometry (FCM) and immunofluorescence (IF) analyses across three different mouse models ***(Fig.2e-2g, Extended Data Fig.4b-4f)***. Moreover, scRNA-seq disclosed enhanced effector/cytokine signaling^21^ in CD8^+^T cells in GFPT2-silenced xenografts ***(Fig.2h)***, which was matched by an uptick in IFNγ and GZMB-positive cells as per FCM ***(Fig.2e-2g)***, indicating a boost in T cell effector function. Recent research has elucidated the complexity and variability of CD8^+^T cell dysfunction^22^, including heightened expression of immune checkpoints such as TIM3^7^, PD-1, LAG3^23^, TIGIT^24^, and CD226. In GFPT2-silenced xenograft models, there was a notable increase in effector/cytokine signaling across TIM3^+^CD8^+^T, PD-1^+^CD8^+^T, and CD226^+^CD8^+^T cells ***(Fig.2h)***. While dendritic cells (DCs) were markedly reduced in shGFPT2 xenografts (***Fig.2d***), and the capacity for tumor–cDC1 interaction in activating cytotoxic T cells was reported^7,25^, GFPT2 knockdown remained effective in HSPC-NPG mice ***(Fig.2a-2b)***, which lack a fully developed DC system^20^. This suggests that DCs are not critical upstream mediator for GFPT2’s function. Furthermore, cell-cell communication analysis leveraging scRNA-seq data confirmed a dominant and direct interaction between tumor cells and CD8^+^T cells ***(Fig.2i)***. Consistently, the survival benefit of GFPT2 silencing was abrogated with the administration of anti-CD8 antibodies in immunologically competent mice bearing EGFR-mutated LLC ***(Fig.2j)***, directing our focus towards understanding how GFPT2 influences the crosstalk between tumor cells and CD8^+^T cells.

### N-glycoproteomes post genetic disruption of tumoral GFPT2

UDP-GlcNAc serves as an essential precursor for protein glycosylation, with the Golgi apparatus’s synthesis of tri- and tetra-antennary N-glycans being particularly sensitive to its concentration^26^. These complex N-glycans, upon binding with galectins, form a molecular lattice that impedes glycoprotein endocytosis^26^, contributing to the development of a ‘glycan coat’, as evidenced by PHA staining targeting β-1-6-N-acetylglucosamine branches ***(Extended Data Fig.5a)***. Diminished GFPT2 expression correlates with reduced UDP-GlcNAc levels, leading to a decrease in cell surface protein glycosylation, and conversely, an increase in GFPT2 results in heightened glycosylation ***(Extended Data Fig.5b-5c)***. Moreover, paralleling UDP-GlcNAc trends ***(Fig.1g)***, enhanced hyper-glycosylation due to EGFR mutation was nullified by GFPT2 suppression, highlighting GFPT2’s central role in abnormal glycosylation patterns associated with EGFR mutations ***(Extended Data Fig.5d)***. Investigations using N-glycoproteomics post-GFPT2 knockdown in H1975 cells revealed significant shifts in surface glycoprotein composition: a majority (60.8 %, 259 out of 426) of the proteins displayed reduced glycosylation, 24.4 % (104 out of 426) remained unaltered, and a minority (14.8 %, 63 out of 426) showed increased glycosylation ***(Extended Data Fig.5e)***. Notably, the subset with decreased glycosylation was characterized by a heightened frequency of glycan modifications and a marked enrichment of sialylated glycans ***(Extended Data Fig.5f-5g)***. This phenomenon is primarily attributed to the proteins’ heightened sensitivity to UDP-GlcNAc fluctuations—especially those with a greater number of N-glycan sites^26^—and to the observed reduction in Neu5Ac, a metabolic derivative of UDP-GlcNAc, subsequent to GFPT2 knockdown^12^. Given the critical role that hyper-glycosylation and sialylation play in the immune evasion mechanisms of cancer cells^27^, we proposed that GFPT2 genetic disruption may instigate a phenotypic shift from ‘masked and immune-resistant’ to ‘unmasked and immune-sensitive’. Furthermore, the application of a K-means clustering algorithm segregated proteins displaying reduced glycosylation into three discrete cohorts ***(Extended Data Fig. 5h)***. Significantly, cluster 2 encompassed 122 proteins that engaged extensively with EGFR, with a remarkable 47.5 % (58 out of 122) implicated directly in the regulation of immune functions ***(Extended Data Fig.5h)***. This evidence suggests that glycosylation variations prompted by genetic interference in GFPT2 might act as a crucial intermediary mechanism in restoring anti-tumor immune operations.

### Tumoral GFPT2 orchestrates cytotoxic responses to CD8^+^T

Subsequently, we evaluated the impact of GFPT2 on the susceptibility of tumor cells to cytotoxicity mediated by CD8^+^T cells. In experiments utilizing EGFR-mutated NSCLC tumor cells as the target and co-culturing them with activated CD8^+^T cells, we discerned an elevated cytotoxic response from CD8^+^T cell in GFPT2-knockdown H1975 cells compared to control groups ***(Fig.3a)***. This enhancement was accompanied by a surge in IFNγ and IL-2 secretion in culture supernatant of co-culture system ***(Fig.3b)***. Conversely, overexpressing GFPT2 in H1299 cells resulted in a reverse trend ***(Extended Data Fig.6a)***. GFPT2 knockdown also reinstated T-cell responsiveness in Osimertinib-resistant PC9/OR2 cells ***(Extended Data Fig.6b-6c)***. Reintroduction of GFPT2 validated the specificity of the observed effects by negating off-target concerns ***(Extended Data Fig.6d)***. The immunosuppressive ligand-receptor axis is recognized as the chief mechanism through which tumor cells exploit to impede CD8^+^T-cell cytotoxicity via direct cellular interaction^28^. N-glycoproteomes revealed diminished glycosylation of three immunosuppressive ligands—PD-L1, CD276 and PVR—following GFPT2 knockdown in H1975 cells, without affecting the antigen presentation protein HLA-I, and other immunosuppressive ligands ***(Fig.3c, Extended Data Fig.6e-6g)***. Immunoblotting analyses substantiated GFPT2’s influence on the expression of total and membrane-bound proteins in H1975 and H1299 cells ***(Extended Data Fig.6h-6i)***. PNGase F enzymatic digestion and IF co-staining ascertained that GFPT2 modulates the membrane trafficking and glycosylation levels ***(Fig.3d, Extended Data Fig.6j)***. These regulatory effects were further corroborated *in vivo* by administrating GFPT inhibitor, AZA ***(Fig.3e, Extended Data Fig.6k-6m)***. Taking the widely reported PD-L1 as a representative, we affirmed the pivotal role of GFPT2-mediated glycosylation in modulating the interaction between tumor cells and CD8^+^T cells. Cells subjected to GFPT2 knockdown or inhibition exhibited a reduced association of PD-1 protein on the cell surface, indicative of a dampened immunosuppressive crosstalk between tumor cells and T cells, and vice versa ***(Fig.3f-3g, Extended Data Fig.6n-o)***. Further, mutating the glycosylation sites of PD-L1 significantly impeded its glycosylation, which could not be restored by GFPT2 overexpression, thereby significantly augmenting CD8^+^T cell cytotoxicity upon co-culturing ***(Fig.3h, Extended Data Fig.6p-q)***.

**Fig.3.**
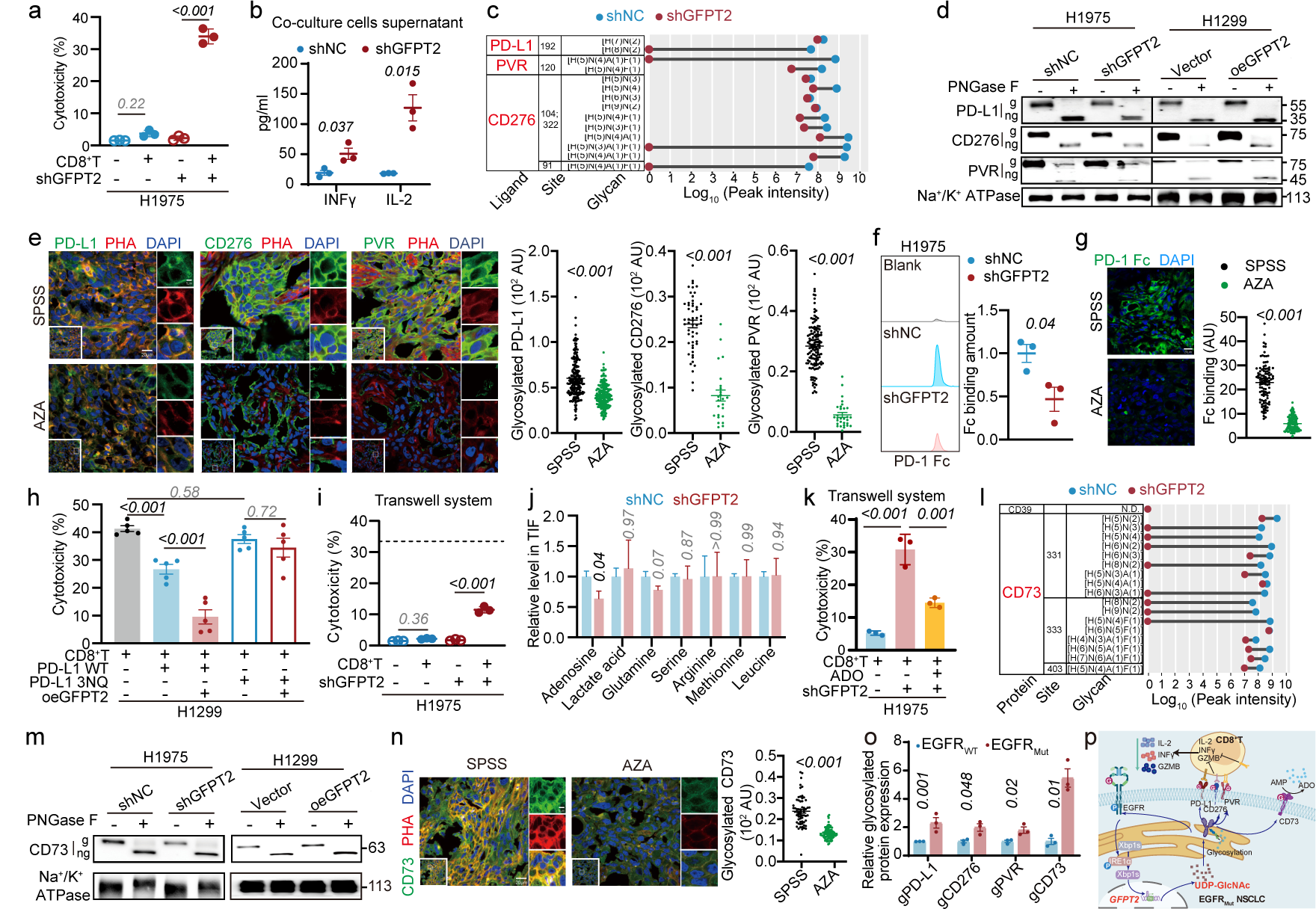
Mechanisms of GFPT2’s influence on tumor cell susceptibility to CD8^+^T cytotoxicity. **a**, FCM analyses of H1975 cell survival in co-culture with activated human CD8^+^T cell sorted from PBMC (n=3). **b**, INFγ and IL-2 levels in cell supernatant of co-culture system were determined by ELISA Kit (n=3). **c**, Analysis of N-linked glycans and peak intensities indicating glycosylation modifications on identified immune checkpoint ligands in shNC and shGFPT2 H1975 cells. **d**, Immunoblotting for protein glycosylation of PD-L1, CD276, and PVR on the cell membrane post-PNGase F digestion, with Na^+^/K^+^ ATPase as a loading control. g, glycosylated form; ng, non-glycosylated form. **e**, Immunofluorescent images of AZA-treated H1975 xenografts, stained for immune checkpoint ligands and glycosylation. SPSS, stroke-physiological saline solution. **f**, FCM quantification of PD-1 Fc binding to PD-L1 on shNC and shGFPT2 H1975 cell membranes, calculated by multiplying the percentage of positive cells by mean fluorescence intensity (n=3). **g**, Immunofluorescent staining for PD-1 Fc binding in tumor mass after AZA treatment. Fc binding amount was calculated according to the fluorescent intensity of single cell. **h**, FCM analyses of H1299 cell survival from co-culture experiments with activated human CD8^+^T cells sorted from PBMC. H1299 cells were transfected with either wild-type PD-L1 or glycosylation site mutant PD-L1, while overexpressing GFPT2. **i**, FCM analyses of H1975 cell survival in Transwell co-culture system with activated human CD8^+^T cell sorted from PBMC (n=3). **j**, HPLC-QTOF/MS analysis of metabolites in the tumor interstitial fluid (TIF) related to the suppression of CD8^+^T cell function (n=5). **k**, Flow cytometric evaluation of H1975 cell viability in a co-culture with activated human CD8^+^T cell using a Transwell setup (n=3) supplemented with adenosine (ADO). **l**, Analysis of N-linked glycans and peak intensities indicating glycosylation modifications on CD73 in shNC and shGFPT2 H1975 cells. **m**, Immunoblotting for post-translational glycosylation of CD73 on cell membranes following PNGase F treatment, using Na^+^/K^+^ ATPase as a loading control. g, glycosylated form; ng, non-glycosylated form. **n**, Immunofluorescent images of AZA-treated H1975 xenografts, with specific labeling for CD73 and its associated glycosylation patterns. **o**, Quantifications of glycosylated PD-L1, CD276, PVR and CD73 expressions in tumor tissues from NSCLC patients with EGFR wild-type and EGFR mutations, based on immunoblotting assay ***(Extended Data Fig.7j)***. **p**, Illustration depicting the mechanisms by which GFPT2 influences tumor cell susceptibility to CD8^+^T mediated cytotoxicity. Data are mean ± s.e.m. **b**, **e**, **f**, **g**, **j**, **n**, **o**, Two-tailed unpaired Student’s t-test. **a**, **h**, **i**, **k**, One-way ANOVA.

Remarkably, an analogous, albeit reduced, effect was discerned using a Transwell system, underscoring GFPT2’s potential to modulate CD8^+^T-cell cytotoxicity through both contact-dependent and independent mechanisms ***(Fig.3i, Extended Data Fig.7a)***. Nutrient and metabolite shift within the tumor microenvironment (TME) is known to affect tumor–immune communication via non-cell contact mechanism^29^. Metabolomic profiling of tumor interstitial fluid (TIF) exposed a notable decrease in adenosine upon GFPT2 knockdown in tumor cells ***(Fig.3j, Extended Data Fig.7b-7c)***, with mass spectrometry confirming a decline from 1.52 μg/ml to 0.78 μg/ml ***(Extended Data Fig.7d)***. Other TIF metabolites, previously associated with CD8^+^T cell functionality^30,31^, remained unchanged ***(Fig.3j)***. In the TIME, adenosine notably suppresses immune activity by engaging A2AR on immune cells, and *in vitro* assays have identified a threshold of 1.5 μg/ml as influential on T cell cytotoxicity in the Transwell setups ***(Extended Data Fig.7e)***. Subsequent analysis of adenosine in supernatants from GFPT2-knockdown tumor cell cultures confirmed a substantial decrease ***(Extended Data Fig.7f)***. When adenosine levels were restored, the increased vulnerability of GFPT2-knockdown H1975 cells to CD8^+^T-cell cytotoxicity in the Transwell setups was effectively mitigated ***(Fig.3k)***, alluding to adenosine’s primary role in the non-cell contact mechanism. Adenosine in TIF arises predominantly from ATP metabolized by membrane-associated enzymes, specifically through ATP degradation to AMP by CD39, followed by conversion to adenosine by CD73^32^. Glycoproteomic analysis did not reveal CD39; however, a significant decrease in glycosylated CD73 on the tumor cell membrane was evident post-GFPT2 knockdown ***(Fig.3l)***. Manipulation of GFPT2 levels—either through knockdown in H1975 cells, overexpression in H1299 cells, or inhibition in xenograft tumors using AZA—consistently demonstrated GFPT2’s regulatory impact on CD73’s total expression, membrane localization, and glycosylation ***(Fig.3m-3n, Extended Data Fig.7g-7i)***.

Consistently in clinical, we observed elevated expression and glycosylation of PD-L1, CD276, PVR, and CD73 in NSCLC patients with EGFR mutation, accompanied with higher GFPT2 expression *(**Fig.3o, Extended Data Fig.7j)***. In summary, GFPT2 downregulation in tumor cells notably diminishes the expression and glycosylation of critical immunosuppressive ligands, thus mitigating their interactions with T cells. Concurrently, it downregulates CD73 expression, curbing the adenosine-mediated inhibition of T cell. Together, these mechanisms heighten the vulnerability of tumor cells to CD8^+^T-mediated cytotoxicity (***Fig.3p***).

### Tumoral GFPT2 modulates CD8^+^T infiltration in TIME

Tumoral GFPT2 knockdown amplifies the intratumoral CD8^+^T cell population ***(Fig.2)***, yet, the proliferation capacity of intratumoral CD8^+^T cells remained unaltered, except minimal decrease in PD-1^+^CD8^+^T ***(Extended Data Fig. 8a)***, suggesting an increased infiltration of CD8^+^T cells into the tumor. A subsequent experiment utilizing immunodeficient mice, engrafted bilaterally with GFPT2-knockdown H1975 cells and control cells, validated enhanced T cell infiltration into the GFPT2-knockdown site within 24 hours post intravenous administration of activated T cells, excluding variations due to T cell exhaustion ***(Fig.4a)***. To understand the mechanisms driving immune cell infiltration, which involves chemotaxis and trans-endothelial migration, we assessed the influence of tumoral GFPT2 on intratumoral CD8^+^T cell dynamics ***(Fig.4b)***. CD8^+^T cells are typically attracted by chemokines such as CCL5 and CXCL9/10/11^33^. Notably, CXCL10 was identified as the sole chemokine with increased levels in TIF following tumoral GFPT2 knockdown ***(Fig.4c, Extended Data Fig.8b)***. This observation was corroborated *in vitro*, where GFPT2-silenced H1975 cells exhibited an upsurge in CXCL10 release ***(Extended Data Fig.8c)***. Transwell migration assays revealed enhanced CD8^+^T cell migration towards GFPT2-knockdown H1975 cells, a process inhibited by CXCL10 neutralization ***(Fig.4d)***, suggesting GFPT2 disruption in tumors preferentially promotes CD8^+^T cell chemotaxis via CXCL10. In tumors chronically exposed to the GFPT inhibitor AZA, vascular normalization was indicated by an increase in elongated vascular length and αSMA^+^ vessels ***(Fig.4e)***. Downregulation of GFPT2 and AZA treatment both resulted in lowered VEGF levels in tumor homogenate and TIF, aligning with recognized role of VEGF in vascular normalization ***(Fig.4f, Extended Data Fig.8d-8e)***. Additionally, silencing GFPT2 in H1975 cells led to reduced VEGF secretion, correlating with a decrease in endothelial cell migration in Transwell co-culture system observed *in vitro* ***(Extended Data Fig.8f-8g)***. Conversely, overexpression of GFPT2 in H1299 cells enhanced endothelial cell migration. This increase was reversible by treatment with the VEGF-specific monoclonal antibody Bevacizumab ***(Fig.4g, Extended Data Fig.8h)***, confirming GFPT2’s regulation of endothelial dynamics via VEGF.

**Fig.4.**
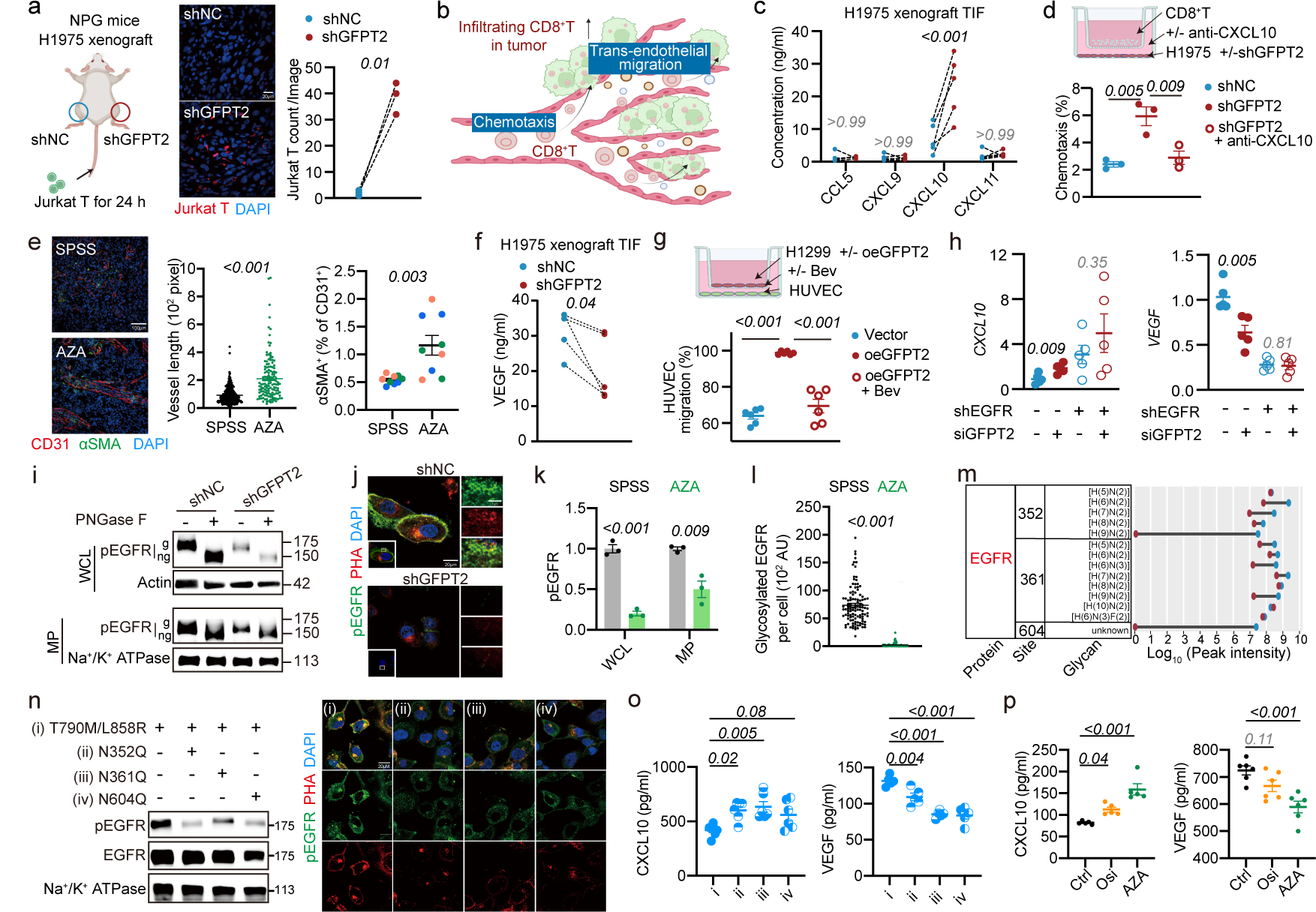
Tumor cell GFPT2 modulates CD8^+^T cell infiltration into the tumor microenvironment via positive feedback on EGFR glycosylation. **a**, IF was used to detect the content of Jurkat T cells within the tumors on both flanks with H1975 xenografts treated with shNC and shGFPT2 (n=3). **b**, Schematic of CD8^+^T cell infiltration into the tumor. **c**, ELISA measurement of chemokines in the TIF of shNC and shGFPT2 H1975 xenografts in mice (n=5). **d**, Transwell co-culture system assessing the dependence of GFPT2-mediated CD8^+^T cell chemotaxis on CXCL10 (n=3). **e**, IF staining to evaluate coverage by α-SMA and tumor vessel length following 2 weeks of treatment with GFPT1/2 inhibitor AZA (n=3, three fields per group). **f**, ELISA measurement of VEGF concentration in the TIF of shNC and shGFPT2 H1975 xenografts in mice (n=5). **g**, Transwell co-culture system to assess the effect of tumor cell GFPT2 intervention on endothelial cells, dependent on VEGF (n=6). Bev, Bevacizumab. **h**, Gene expression of *CXCL10* and *VEGF* in H1975 cells after specific interventions (n=5). **i**, Immunoblotting for glycosylation of pEGFR in whole cell lysate and cell membrane fraction following PNGase F digestion, with Actin and Na^+^/K^+^ ATPase serving as loading controls, respectively. **j**, Immunofluorescent images of shGFPT2-treated H1975 cells, stained for pEGFR and glycosylation (representative images from three independent experiments). **k**, Western blot quantifications of pEGFR in whole cell lysate and cell membrane fraction following treatment with AZA, with Actin and Na^+^/K^+^ ATPase as loading controls, respectively (n=3). **l**, Quantification of glycosylated pEGFR per cell in AZA-treated H1975 xenografts. **m**, Analysis of N-linked glycans and peak intensities indicating glycosylation modifications on EGFR in shNC and shGFPT2 H1975 cells. **n**, Examination of intracellular pEGFR levels by immunoblotting and assessment of membrane localization and glycosylation levels by IF in H1299 cells transfected with wild-type EGFR or site-mutated EGFR and overexpressing GFPT2. **o-p**, ELISA measurement of CXCL10 and VEGF secretion in cell supernatants of **(o)** H1299 cells (n=5) and **(p)** PC9/OR2 cells (n=6). Data are mean ± s.e.m. except in **a**, **c**, **f**, Two-tailed paired Student’s t-test. **e**, **k**, **l**, Two-tailed unpaired Student’s t-test. **d**, **g**, **h**, **o**, **p**, One-way ANOVA.

GFPT2 distinctly regulates CXCL10 and VEGF, which are not glycosylated, by modifying their gene expression, in contrast to its effect on glycosylated immunosuppressive ligands ***(Fig.4h, Extended Data Fig.6e*)**. In EGFR-mutated cells, the EGFR/AKT/IRF1 axis transcriptionally downregulates CXCL10^2,34^, while the EGFR/ERK/NRP1 pathway upregulates VEGF transcriptionally^35,36^. Absence of EGFR in H1975 cells negates GFPT2’s effect on CXCL10 and VEGF mRNA and secretion levels ***(Fig.4h, Extended Data Fig.8i)***, yet does not affect PD-L1, CD276, PVR and CD73 expression ***(Extended Data Fig.8j)***. These findings highlight GFPT2’s specific influence on CXCL10 and VEGF, mediated by EGFR. GFPT2’s role in EGFR regulation was confirmed by knockdown and overexpression studies in H1975 and H1299 cells, respectively. Immunoblotting highlighted GFPT2’s impact on global and membrane-bound pEGFR ***(Extended Data Fig.8k)***, while PNGase F treatment and IF co-staining clarified its effect on pEGFR’s membrane trafficking and glycosylation ***(Fig.4i-4j, Extended Data Fig.8l-8m)***. These regulatory mechanisms were further validated *in vitro* and *in vivo* using the GFPT inhibitor AZA ***(Fig.4k-4l, Extended Data Fig.8n-8p)***. Glycopeptidomic analysis identified primary glycosylation of EGFR at asparagine residues 352, 361, and 604 ***(Fig.4m)***. Mutagenesis of these asparagines (N) to glutamines (Q) in mutant EGFR T790M/L858R, when transfected into H1299 cells, resulted in diminished membrane-bound active pEGFR and glycosylation following GFPT2 overexpression ***(Fig.4n)***. These modifications led to increased extracellular CXCL10 secretion and a marked decrease in VEGF secretion ***(Fig.4o)***.

As summarized, in EGFR-mutated NSCLC cells, GFPT2, upregulated by the EGFR/IRE1α/Xbp1s pathway, feedback-regulates EGFR glycosylation, in turn affecting CXCL10 and VEGF secretion and CD8^+^T cell infiltration in an EGFR-dependent manner. Moreover, targeting GFPT2 presents advantages for its efficacy in drug-resistant cells ***(Fig.4p and Fig.2b)***, thus might bypassing the transient effect on immune infiltration seen with long-term EGFR TKI resistance^34^.

### Clinical potential of GFPT2 inhibition

IHC analysis of human tumors corroborates animal data, showing increased EGFR hyperactivation, heightened PHA staining, and significantly reduced CD8^+^T cell infiltration in GFPT2-positive tumors versus GFPT2-negative ones ***(Fig.5a)***. This aligns with the diminished therapeutic response to Pembrolizumab and Nivolumab in patients with high GFPT2 expression ***(Fig.5b)***, indicating the therapeutic value of GFPT2 as a target. However, the current lack of GFPT2-selective agents, with non-specific inhibitors like AZA and DON affecting GFPT1 as well, raises concerns about off-target toxicity due to GFPT1’s essential physiological roles and pervasive presence^8^. Our next aim is to isolate undesirable off-tumor, on-target effects by developing GFPT2 isoform-specific inhibitor.

**Fig.5.**
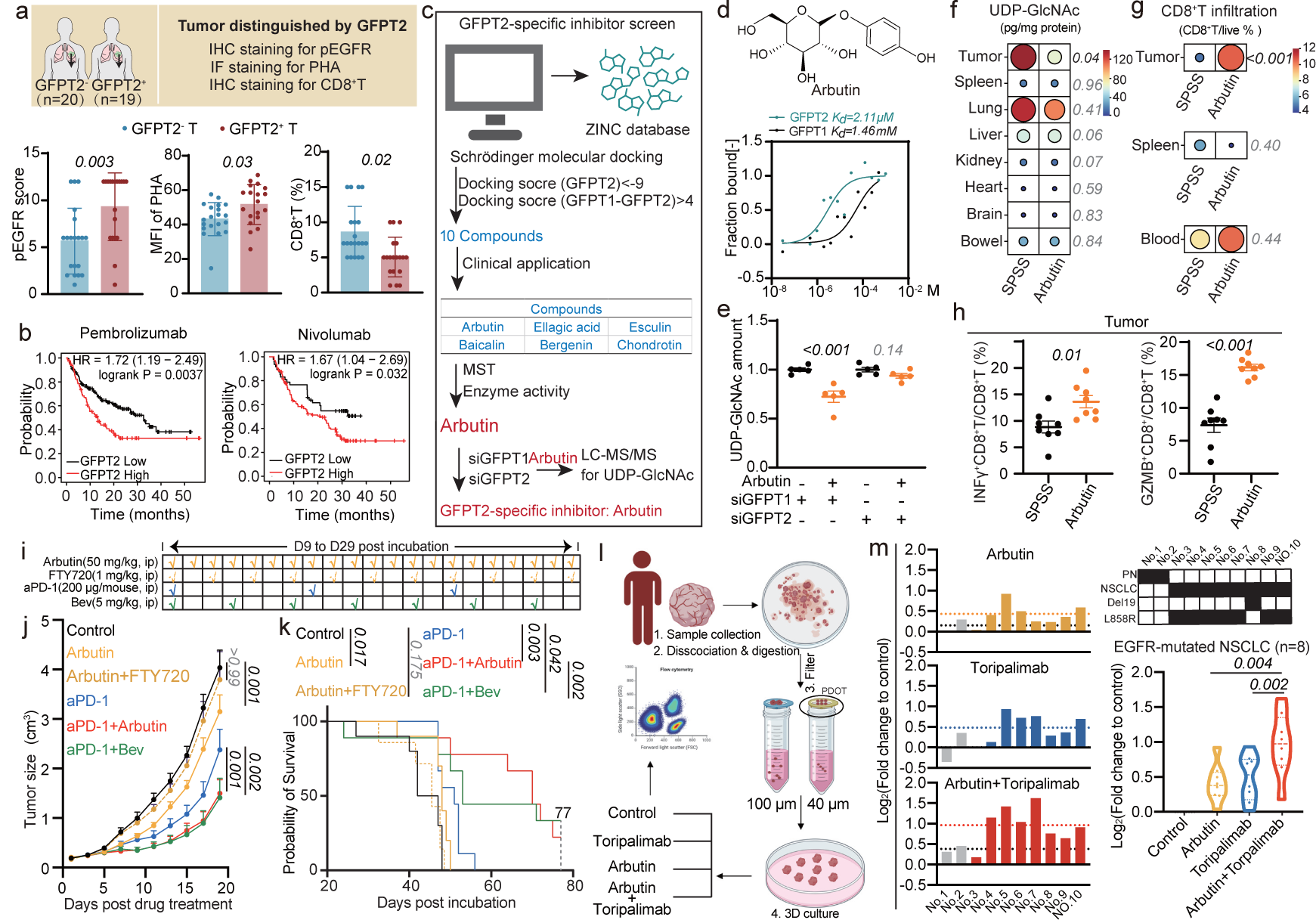
Identifying and validating a GFPT2-specific inhibitor to enhance immunotherapy in EGFR-mutated NSCLC. **a**, Quantifications of pEGFR IHC staining, PHA IF staining and CD8^+^T IHC staining in GFPT2 negative (GFPT2^-^) and GFPT2 positive (GFPT2^+^) NSCLC patients. **b**, The relationship between GFPT2 expression and clinical outcomes in response to Pembrolizumab and Nivolumab treatment. **c**, GFPT2-specific inhibitor screening process. **d**, The structure of Arbutin obtained by screening and the curve of Arbutin binding to GFPT2 and GFPT1 analyzed by microscale thermophoresis assay. **e**, Effects of silencing GFPT1 or GFPT2 on the inhibitory activity of Arbutin on intracellular UDP-GlcNAc production (n=5). **f**, HPLC-QTRAP/MS quantification of UDP-GlcNAc levels in tumors and major organs following three weeks of Arbutin administration in EGFR-mutated LLC xenograft mice (n=8), with data visualized as mean values in a heatmap. **g**, FCM analysis of changes in CD8^+^T cell infiltration within the tumor (n=8), as well as alterations in CD8^+^T cells in peripheral blood and spleen following Arbutin administration (n=3). Data was presented as means by heatmap. **h**, FCM analysis of changes in CD8^+^T cell activation within the tumor (n=8). **i**, Schematic diagram of drug administration. **j-k**, Tumor volume and survival analysis of C57BL/6N mice harboring EGFR-mutated LLC tumors with indicated treatments (n=10 for control; n=10 for Arbutin; n=7 for Arbutin+FTY720; n=10 for aPD-1; n=11 for aPD-1+Arbutin; n=9 for aPD-1+Bev). **l**, Schematic representation of the PDOT evaluation. **m**, FCM evaluation of tumor cell death in PDOTs following treatment with Arbutin, Toripalimab, or their combination. Data are mean ± s.e.m. **a** (MFI of PHA, CD8^+^T proportion), **f**, **g**, **h**, Two-tailed unpaired Student’s t-test. **e**, **m**, One-way ANOVA. **j**, Two-way ANOVA for tumour size. **a** (pEGFR score), Mann-Whitney test. **k**, Mantel-Cox test.

Contrary to glutamine, which binds similarly to both GFPT1 and GFPT2 isoforms, fructose-6-phosphate exhibits unique binding profile for GFPT2, distinct from GFPT1 ***(Extended Data Fig.9a-9b)***. This differential binding informed our search for GFPT2 inhibitors from a natural product library, guided by the structural similarity of many natural compounds to glycosides, resembling fructose-6-phosphate. Employing a high-content docking system, we identified 10 compounds from 190,295 candidates (https://zinc15.docking.org/substances/subsets/natural-products+for-sale/) with high GFPT2 affinity and isoform selectivity ***(Fig.5c)***. Considering the limitations of contrast agents and antibiotics, we narrowed down to 6 promising hits. We refined our search with HPLC-QTRAP/MS to pinpoint compounds that reduce cellular GFPT activity and used microscale thermophoresis (MST) to confirm isoform-specific binding. This comprehensive approach identified Arbutin (Ar) as the most active compound with the highest isoform specificity ***(Fig.5d-5e, Extended Data Fig.9c-9d)***. GFPT2 knockdown negated Arbutin’s impact on UDP-GlcNAc level, whereas GFPT1 silenced showed no effect, affirming Arbutin’s isoform selectivity ***(Fig.5e, Extended Data Fig.9e)***. Furthermore, Arbutin altered the response of H1975 and PC9/OR2 cells to CD8^+^T cell cytotoxicity, diminishing inhibitory ligands PD-L1, CD276, PVR, and CD73 expression, and boosting T-cell chemotaxis, paralleling GFPT2 knockdown effects ***(Extended Data Fig.10a-10e)***. Administration of Arbutin *in vivo*, exploiting GFPT2’s tumor-specific expression, selectively disrupts tumor HBP metabolism and encourages CD8^+^T cell infiltration and activation, leading to suppressed tumor growth without affecting HBP metabolism in vital organs or disturbing the peripheral immune equilibrium, confirming its safety ***(Fig.5f-5h, Extended Data Fig.10f-10k)***. Our studies further explored Arbutin’s potential to boost EGFR-mutated NSCLC’s responsiveness to immune checkpoint blocker (ICB) therapy through GFPT2-specific inhibition. In immunocompetent mice with EGFR-mutated LLC xenografts, those treated with Arbutin and anti-PD-1 antibodies (aPD-1) combination exhibited improved tumor control and survival compared to those treated with a single agent or placebo ***(Fig.5i-5k)***. The treatment was well tolerated, with no observed toxicity or weight loss ***(Extended Data Fig.10h)***. Against the backdrop of the aPD-1 and Bevacizumab regimen, Chinese approved treatment for EGFR-mutated non-squamous NSCLC and a first globally^37^, Arbutin’s combination was found to inhibit tumor growth comparably and enhanced survival benefits ***(Fig.5j-5k)***.

To examine GFPT2 inhibition as a strategy to surmount ICB resistance in human EGFR-mutated NSCLC cancer, we conducted *ex vivo* patient-derived organotypic tumor spheroid (PDOT) from human tumors explants ***(Fig.5l)***^38,39^. PDOTs from 8 NSCLC patients harboring EGFR mutations along with 2 cases of pulmonary nodule (PN) were cultured with GFPT2 specific inhibitor (Arbutin, 50 μM) alone or in combination with a commercial aPD-1 (Toripalimab, 200 μg/ml). The analysis showed a significant greater reduction in tumor growth with combined GFPT2 inhibitor plus PD-1 blockade for EGFR-mutated adenocarcinoma subgroups, compared to either treatment alone. These results support the potential of GFPT2 isoform-specific inhibition as a viable strategy to enhance immunotherapy efficacy in EGFR-mutated NSCLC.

## Discussion

Here, we demonstrate that GFPT2 orchestrates a reprogramming of the metabolic-dependent protein glycosylation profile, thereby promoting an immunosuppressive environment (***Extended Data Fig.10l***). Inhibition of GFPT2 sensitizes EGFR-mutated NSCLC to immunotherapeutic interventions, paving the way for future translational research.

Distinct from their healthy counterparts, tumor cells displayed a markedly altered ‘glycan coat’, characterized by significant alterations in glycan modifications on their cell surface glycoproteins^40^. Such aberrant glycosylation reshapes how the immune system recognizes the tumor and promotes immunosuppressive signaling through glycan-binding receptors, underscoring the therapeutic potential of glycan targeting^27,41^. Our unbiased analysis has uncovered that EGFR mutations precipitate an anomalous increase in intracellular UDP-GlcNAc, leading to hyperglycosylation of tumor cells. Although HBP, culminating in UDP-GlcNAc, is known to influence innate immunity^42^, its role in cancer immunity remains to be fully defined. We have identified the rate-limiting enzyme isoform GFPT2 within the HBP as a novel therapeutic target for enhancing antitumor immunity in EGFR-mutated NSCLC. GFPT2, unlike other constitutively expressed enzymes in the HBP, is notably upregulated in response to EGFR mutations and is minimally expressed in normal tissues. Targeting GFPT2 with a specific inhibitor represents a clinically relevant strategy to selectively disrupt tumor metabolism, offering precision without compromising the HBP’s functions and the immune environment in healthy organs.

Mechanistically, the upregulation of GFPT2 bypasses the canonical downstream EGFR signaling pathways, ERK/AKT, in favor of an undocumented EGFR/IRE1α/Xbp1s pathway. Notably, this novel route is likewise operational in Osimertinib-resistant phenotypes, indicating the potential of GFPT2-targeted therapies to benefit patients who are resistant to EGFR TKI treatments. Analogously, KRAS mutations engage IRE1α/Xbp1s^43^, and a concomitant increase in GFPT2 expression is observed in KRAS/LKB1-mutated NSCLC^12^. The potential of GFPT2-targeted interventions in enhancing antitumor immunity in KRAS-mutated contexts remains to be further elucidated. Beyond the scope of glycan-blocking or degradative tactics, metabolic-dependent glycosylation regulation by targeting GFPT2 encompasses a wider array of immune regulatory membrane proteins, thereby not only priming tumor cells for CD8^+^ T cell-mediated cytotoxicity but also fostering CD8^+^T cell tumor infiltration, effectuating multifaceted modulation of the cancer-immune cycle^44^. Our experiments also confirmed that combining GFPT2 inhibition with aPD-1 extends survival beyond what is observed in the combination of Bevacizumab and aPD-1. While monotherapy with GFPT2 inhibitor significantly improve immune-mediated tumor suppression, its effect is modest. This limitation may be attributed to the compound’s micromolar affinity. The development of novel potent inhibitors to realize the full therapeutic potential of GFPT2 inhibition as a monotherapy warrants further investigation.

## Methods

### Clinical Specimens

We collected 18 fresh NSCLC tissue samples and 2 pulmonary nodule tissue samples (from patients P01-P20) for metabolomic analyses, immunoblotting, and PDOT evaluations from The First Affiliated Hospital of Nanjing Medical University and The Affiliated Brain Hospital of Nanjing Medical University. The study received approval from the Research Ethics Committees of both institutions (2023-KL008-02, 2019-SR-450), with all participating patients providing written informed consent. Patient information for P1-P10 is listed in ***Supplementary Table.S1***, and for P11-P20 in ***Supplementary Table.S13***. Paired tumor tissues and non-cancerous adjacent tissues samples from 5 patients with wild-type EGFR (P01-P05) and 5 with EGFR-mutated NSCLC (P06-P10) were used for unbiased metabolomic analysis (data presented in ***Fig.1a***). Tumor and matched adjacent non-tumor samples from 3 wild-type EGFR patients (P01, P04, P05) and 3 EGFR-mutated patients (P06, P07, P08) underwent immunoblot analysis (data presented in ***Fig.1d, Fig.1i, Extended Data Fig.2g, Fig.3o, Extended Data Fig.7j***). Two pulmonary nodule samples (from patients P11, P12) and samples from 8 tumor samples from EGFR-mutated NSCLC patients (P13-P20) were utilized to prepare PDOT models (data shown in ***Fig.5m***).

Furthermore, we collected 39 paraffin-embedded NSCLC samples (from patients P21-P59) from The First Affiliated Hospital of Nanjing Medical University for clinical validation of the correlation between GFPT2 expression, EGFR activation, hyperglycosylation of tumors, and CD8^+^T cell infiltration. Patient information for P21-P59 is detailed in **Supplementary *Table.S11***. Through immunohistochemical analysis of GFPT2, these 39 patients were classified into a GFPT2-positive expression subgroup (n=19, P21-P39) and a GFPT2-negative expression subgroup (n=20, P40-P59), followed by pEGFR, CD8^+^T cell IHC, and PHA lectin immunofluorescence (IF) staining (data presented in ***Fig.5a***; original histological imaging provided in ***Supplementary Fig.S2-S4***). Notably, tumor samples and matched NAT from 4 patients (P21, P27, P33, and P36) underwent GFPT2 immunohistochemical analysis (data presented in Fig.1f, with original histological imaging shown in ***Supplementary Fig.S1***)

In total, three publicly available clinical datasets were analyzed for the tissue distribution expression of HBP metabolic enzymes. (1) Proteomic data from 8 EGFR wild-type (EGFR_WT_) and 47 EGFR-mutated (EGFR_Mut_) NSCLC patients, sourced from the Database of Genotypes and Phenotypes (ph-s001954.v1.p1), were utilized to investigate the induced expression of HBP metabolic enzymes under EGFR mutation conditions (data presented in ***Fig.1c***). Patient information and related data are listed in ***Supplementary Table.S3***. (2) Two independent human proteomic databases, the tissue-based map of the human proteome (PRJNA183192) and the Human Proteome Map database (available at http://www.humanproteomemap.org/query.php), were utilized to analyze the expression of HBP metabolic enzymes across various major organs (data presented in ***Fig.1e*** and ***Extended Data Fig.1f***). Patient information and related data are listed in ***Supplementary Table.S4***. (3) Proteomic profiles from matched tumor and adjacent non-tumor tissues of 66 NSCLC patients with EGFR mutations (sourced from the Database of Genotypes and Phenotypes, ph-s001954.v1.p1), along with those from an additional independent cohort of 51 EGFR-mutated NSCLC patients (IPX0001804000), were analyzed to determine whether GFPT2 is overexpressed in tumors due to EGFR mutation induction (data presented in ***Fig.1f***). Patient information and related data are listed in ***Supplementary Table.S4***.

### Cell lines and cell culture

All cell lines utilized were sourced from the Cell bank of Chinese Academy of Sciences (Shanghai, China). Human NSCLC cell lines encompassing H1975 (EGFR T790M/L858R), HCC827 (EGFR Del19), PC9 (EGFR Del19), and H1299 (EGFR WT), alongside lymphocytes Jurkat T were cultured in RPMI 1640, enriched with 10 % fetal bovine serum (FBS), 100 U/mL penicillin and 100 μg/mL streptomycin. The mouse progenitor B cell line, BaF3, was cultured in RPMI 1640 supplemented with 10 % FBS, 1 % antibiotic mixture, and 15 ng/mL IL −3. The mouse cancer cell line LLC and human umbilical vein endothelial cell (HUVEC) were cultured in DMEM containing 10 % FBS and a 1 % antibiotic mixture. All aforementioned cell lines were incubated at 37 °C with a 5 % CO_2_-humidified atmosphere.

To establish EGFR TKI-resistant sublines, parental PC9 cells were subjected to two distinct Osimertinib adaptation protocols. The first regimen involved a stepwise adaptation of PC9 cells to increasing Osimertinib concentrations, ranging from 10 to 500 nM over a span of six months, leading to the PC9/OR1 cell line. In the second regimen, cells were initially exposed to 10 nM Osimertinib for two weeks, followed by prolonged culture in 50 nM Osimertinib for an additional six months, generating the PC9/OR2 cell line. For resistance validation, PC9, PC9/OR1 and PC9/OR2 cells were seeded at 10,000 cells/well in 96-well plates, and challenged with Osimertinib concentrations of 10, 100, 1000 and 5000 nM for 48 hours, followed by cytotoxicity assessment via a Cell Counting Kit-8.

To establish EGFR-mutated murine cell line, LLC cells (designated as EGFR-mutated LLC) were engineered to stably overexpress the EGFR T790M/L858R mutation via retroviral transduction using a pLenti-GIII-CMV-CBH-GFO-2A-Puro vector. Specifically, LLC cells were allocated at a density of 2 × 10^5^ cells per well in a 6-well plate and, following a 24 h incubation, were transduced with high-titer lentivirus at a specified multiplicity of infection (MOI = 20) in complete medium. Subsequently, cells exposed to the pertinent lentivirus were subjected to a selection pressure using 4 µg/mL puromycin (Beyotime, Nanjing, China) over a 2 - 4 day period, prior to being enlisted for subsequent bioassays. The resultant EGFR-mutated LLC cells were then characterized based on various markers: increased EGFR phosphorylation, upregulated GFPT2 expression, enhanced glycosylation of immune suppressive ligands (PD-L1, CD276, PVR, CD73), and elevated secretion of CXCL10 and VEGF. Collectively, these traits confirmed the cell line’s ability to mimic the immunosuppressive phenotype associated with EGFR-mutant NSCLC.

### Isolation of CD8^+^T from human PBMC

Pre-coat a 12-well plate overnight at 4 °C using CD3 antibody (5 μg/ml, 317326, Biolegend), and before inoculation, rinse the wells three times with phosphate buffer saline (PBS) solution to remove any residual CD3. Collect fresh human peripheral blood into a 50 mL EP tube, add an equal volume of PBS, and gently mix by pipetting. Add 1.5 times the volume of the blood of Ficoll solution (00421, Serumwerk Bernburg AG, Germany) to another 50 mL EP tube, and gently layer the mixed blood along the side of the tube onto the Ficoll solution, avoiding mixing the two. Then centrifuge at room temperature at 800 g for 20 minutes, requiring slow acceleration and deceleration (acceleration 3, deceleration 0). After centrifugation, the solution is separated from top to bottom into plasma, peripheral blood mononuclear cells (PBMCs), lymphocyte separation liquid, and granulocytes. Discard the plasma layer, collect the PBMCs into a new 15 ml EP tube, and wash twice with 5 volumes of PBS, each time centrifuging at room temperature at 1500 rpm for 10 minutes. Discard the supernatant, resuspend the cells in 4 ml of red blood cell lysis solution (C3702, Beyotime), and let stand at room temperature for 5 minutes to lyse the remaining red blood cells. Then add 8 ml PBS to stop the lysis and centrifuge at room temperature at 1500 rpm for 10 minutes. Resuspend the cells in PBS and follow the procedure in the EasySepTM Human CD^+^8 T Cell Iso Kit (17953, STEMCELL) manual to extract CD8^+^T cells: add cocktail to the cell suspension at 50 μl/ml, invert several times, and incubate for 5 minutes at room temperature. Add magnetic beads to the cell suspension at 50 μl/ml, invert several times, place on a magnetic rack, and after 5 minutes, transfer the supernatant to a new 15 mL EP tube. Wash twice with 5 volumes of PBS, each time centrifuging at room temperature at 1500 rpm for 10 minutes. Then resuspend and count in 1640 complete medium containing CD28 (5 μg/ml, 302934, Biolegend) and IL-2 (10 ng/ml, 200-02, Pepertech), and finally inoculate at a density of 1 × 10^6^ cells/ml onto the pre-coated CD3 12-well plate. After 48 hours, transfer and continue to culture in 1640 complete medium containing IL-2, stimulating weekly with CD3 and CD28 antibodies as described above.

### PDOT isolation and drug treatment

Fresh tumors from NSCLC patients were thoroughly washed in PBS and minced into a pulp under sterile and cool conditions. Digestive fluid was added at a ratio of 15 mL per 1 g of tumor (complete 1640 medium, 1 mg/ml type IV collagenase, 15 mmol/l HEPES) and digested on a shaker at 37 °C at 90 rpm for 20 minutes, after which an equal volume of complete 1640 medium was added to terminate the digestion. Spheroids ranging from 40-100 μm in diameter were collected by sieving twice. To lyse red blood cells, 5 ml of red blood cell lysis solution was added and the mixture was left to stand at room temperature for 5 minutes, followed by the addition of twice the volume of PBS to cease lysis, and centrifuged at 1500 rpm for 5 minutes at room temperature. The spheroids were resuspended in ECM gel (2.5 mg/ml type I collagen from mouse tails, 2 mM acetic acid, 100 mM HEPES, 1 g/l sodium bicarbonate) and seeded into a pre-cooled 48-well plate at a density of 100 clusters per 150 μl of ECM. After 24 hours, complete 1640 medium was added to continue culture. The similarity in the proportion of immune cells within the spheroids to that within the tumor was validated by flow cytometry (FCM). The tumor classification and identification of EGFR mutations were confirmed by pathology experts.

PDOTs received treatment with 50 µM Arbutin, 200 µg/ml of the commercialized anti-PD-1 antibody (aPD-1), Tripolyimumab, or their combination. Four days post-treatment, PDOTs were enzymatically dissociated into single cells for FCM. Non-specific antibody binding was precluded with Human TruStain FcX™ (422301, Biolegend, USA). A suite of fluorochrome-conjugated monoclonal antibodies was then employed to identify surface antigens. Five minutes prior to FCM, propidium iodide (PI) staining was performed. The subset of CD45^-^PI^+^ cells was delineated to quantify the proportion of tumor cell death within the PDOTs. Gating strategies are presented in ***Supplementary Fig6***.

### Plasmid transfections, viral infections and RNA interference

For suspension-cultured cell BaF3, 7 × 10^7^ cells were suspended with 100 μl PBS solution in electric shock cups and transfected with 1 μg/μl plasmid (10 μl per cup, 0.1 μg/μl final) under 500 μF, 500 ω, and 160 V conditions provided by Gene importer (SCIENTZ-2C, Ningbo China). Cells were incubated on the ice for 10 min post electric shock and then seeded into a 6-well plate with complete medium. The transfected cells were used in following bioassays 2 - 4 days post-transfection. BaF3 cells upon the introduction of oncogenic EGFR-mutated forms were constructed. pcDNA3.1-EGFR WT, pcDNA3.1-EGFR T790M, pcDNA3.1-EGFR L858R, pcDNA3.1-EGFR Del19, pcDNA3.1-EGFR T790M/L858R, and pcDNA3.1-EGFR T790M/Del19 used here were purchased from Nanjing Jinbeijin Biotechnology Co., Ltd. (Nanjing, China).

For adherent cells, parental cells were seeded at 5 × 10^5^ cells per well into a 6-well plate and transfected after 24 hours with 1 μg/μl plasmid (1 μl per well, 0.5 pg/μl final) using lipofectamine 3000 (Invitrogen, L3000015) (2.5 μl per well) in serum-free medium. The medium was replaced with fresh complete medium at 6 hours post-transfection. The transfected cells were then harvested and used in following bioassays 2 - 4 days post-transfection. Overexpressing-GFPT2, GFPT2 promoter containing Dual-Luciferase H1299 cells and other overexpressing cells were constructed as the method described above. pEGFP-N1-GFPT1, pEGFP-N1-GFPT2, pGL4.10-Luciferase-GFPT2-Promoter, pGL4.10-Luciferase-GFPT2-Promoter Mut, pcDNA3.1-Flag-N1-EGFR T790M/L858R/N352Q, pcDNA3.1-Flag-N1-EGFR T790M/L858R/N361Q and pcDNA3.1-Flag-N1-EGFR T790M/L858R/N614Q used here were purchased from Nanjing Jinbeijin Biotechnology Co., Ltd. (Nanjing, China).

Parental cells were seeded at 5 × 10^5^ cells per well into a 6-well plate and transfected after 24 hours with 20 μM siRNA (2 μl per well, 40 pmol final) using lipofectamine RNAiMAX (Invitrogen, 13778150) (5 μl per well) in serum-free medium. The medium was replaced with fresh complete medium at 6 hours post-transfection. The transfected cells were then harvested and used in following bioassays 2 - 4 days post-transfection. The sequences of siRNA targeting GFPT2, GFPT1 and Xbp1s was UCCACUUUUAAGUCCAUGC (siGFPT2), UUCGAAGUCAUAGCCUUUG (siGFPT1), UCUGAAGAGUCAAUACCGCTT (siXbp1s), respectively.

pLenti-GIII-CMV-CBH-GFO-2A-Puro EGFR T790M/L858R and pLVX-shGFPT2-zsGreen-PGK-Puro were purchased from Nanjing Jinbeijin Biotechnology Co., Ltd (Nanjing, China). The sequence of shGFPT2 was UCCACUUUUAAGUCCAUGC. pLenti-CMV-hU6-shEGFR-CBH-gcGFP-Puro was purchased from Shanghai GK Gene Medical Technology Co., Ltd (Shanghai, China) and the sequence of shEGFR was GCGAUACUCUACCUCCUUCU. Parental cells were seeded at 2 × 10^5^ cells per well into a 6-well plate and infected after 24 hours with high-titer lentivirus in complete medium at the volume determined by specific multiplicity of infection (MOI). Cells infected with the indicated lentivirus were treated with puromycin (ST551, Beyotime) for 2 - 4 days and then used in following bioassays. The MOI of H1975, HCC827 and LLC was 50, 20 and 20, respectively and the concentration of puromycin was 2, 1 and 4 μg/ml, respectively. Stable overexpressing-EGFR T790M/L858R LLC cells (EGFR-mutated LLC), stable knockdown-GFPT2 H1975 (shGFPT2 H1975) and PC9OR2 (shGFPT2 PC9OR2) were established as the method described above.

### Ex vivo co-culture

5 × 10^4^ tumor cells were plated in each well of a 48-well plate. These cells were then exposed to pre-activated T cells at a specific effector-to-target ratio (2:1). After a 48-hour co-culture period, cells were stained with 5 μg/ml PI (ST1569-10mg, Beyotime) and immediately analyzed using the CytoFLEX flow cytometer (Beckman Coulter, USA). Tumor cells were distinguished based on GFP positivity or CD45 negativity. Gating strategies are presented in ***Supplementary Fig6***. In order to verify whether GFPT2 in tumor cells may influence T cells in a non-cell-contact manner, the T cell cytotoxicity experiments described above were also conducted in Transwell inserts.

In the CD8^+^T cell chemotaxis assay, tumor cells were seeded at a density of 2 × 10^5^ per well in the bottom chamber of 5-μm-pore Transwell inserts (3421, Costar). Pre-stained CD8^+^T cells were placed in the upper chamber at 1 × 10^5^ cells/well and incubated for 48 h, either with or without 1.2 μg/ml CXCL10 neutralizing antibody (33016, R&D, USA). Following incubation, T cell migration to the lower chamber was quantified using a CytoFLEX flow cytometer (Beckman Coulter, USA).

To evaluate the impact of tumor GFPT2 expression on vascular cells, HUVEC were seeded in 24-well plates at a density of 2 × 10^5^ cells/well and subjected to a wound healing assay. Once HUVEC reached 90-100 % confluence, a standardized wound was created in the cell monolayer using a pipette tip, with the well bottoms marked to guide the initial imaging of the wounded areas. Subsequently, tumor cells were introduced into the upper chamber of Transwell inserts and co-cultured with the indicated HUVECs for an additional 24 h. To neutralize the augmented VEGF secretion resulting from GFPT2 overexpression in tumor cell, 6 μg/ml Bevacizumab (Roche) was administered. At the 24-hour mark, wound gap was imaged using the Lionheart FX^TM^ Intelligent Live Cell Imaging Analysis System (Bio-Tek Instruments, USA). The wound gaps were measured by Image J software, and migration rates were calculated using the formula: *Migration rate = (wound gap (0 h) - wound gap (24 h))/wound gap (0 h)*, where wound *gap=wound area/wound length*.

### Animal treatment and tumor challenges

Animal care and experimentation received approval from the Institutional Animal Care and Use Committee (IACUC) of China Pharmaceutical University and were conducted under the National Research Council’s Guide for the care and use of laboratory animals and animal welfare. Female Balb/c Nude mice and C57BL/6N mice, both aged 4 - 5 weeks, were obtained from the Beijing Vital River Laboratory Animal Technology Co., Ltd. (Beijing, China). Additionally, female immunodeficient NOD.Cg-*PrkdscidIl2rgtm1Vst/Vs* (NPG) mice, alongside their humanized counterpart model HSPC-NPG (created by transplanting human CD34^+^ hematopoietic stem and progenitor cells into NPG mice), were obtained from Beijing Vitalstar Biotechnology Co.,Ltd. (Beijing, China). Mice were housed in specific-pathogen-free facilities at the China Pharmaceutical University (animal authorization reference number: SYXK2021-0011), where they were maintained under a controlled environment of 22 – 24 °C, 50–60 % humidity and a 12 h light/dark cycle (08:00 to 20:00 light on) with access to standard laboratory food and water ad libitum. Pre-inoculation screenings ensured mice were free from pathogens and mycoplasma.

To evaluate Azaserine’s (AZA) effect on tumor progression (***Extended Data Fig.3***), Balb/c nude mice were subcutaneously implanted with 1.0 × 10^7^ H1975 cells on the left flank and stratified into SPSS-treated or AZA-treated groups based on initial tumor volume. AZA was administered at 4 mg/kg intraperitoneally every other day to inhibit GFPT1/2 activity. Tumor growth was monitored bi-daily by caliper measurements using the formula *length × width*^2^ *× 0.5*. The study endpoint was reached when any group’s mean tumor volume exceeded the ethical threshold of 1,200 mm^3^ or upon signs of compromised animal welfare. Mice were euthanized by CO_2_ inhalation, and tumor weights were recorded at the endpoint.

In the studies depicted in ***Fig.2***, both NPG and HSPC-NPG mice were subcutaneously inoculated 1.0 × 10^7^ H1975 or PC9/OR2 cells featuring stable GFPT2 knockdown (shGFPT2) on their right flank, and an equivalent number of control cells (shNC) on the left flank. In a separate set-up, C57BL/6N immunocompetent mice were introduced subcutaneously on the right hind flank with 5 × 10^5^ EGFR-mutated LLC (EGFR T790M/L858R) cells, with or without GFPT2 interference. For GFPT2 interference subgroup, FTY720 was administrated (*1 mg/kg, i.p. qod*) to inhibit lymph node egress starting on day 6 post cell injection. Anti-CD8 group (*250 μg/mouse, i.p, qwd*) was employed to neutralize CD8^+^T cell activity starting on day 8 post cell injection. Upon the emergence of palpable tumors, tumor volume was periodically assessed every two days using the formula *length × width*^2^ *× 0.5*. Experimental endpoints were designated upon any group’s average tumor size attaining the ethical limit of 1200 mm^3^ or if the animals showed any signs indicative of adverse health. CO_2_ inhalation was used to euthanize mice. Tumor weights were determined at the endpoint. Sample sizes were not predetermined through statistical methodologies; however, each experimental group comprised a minimum of five mice. Randomization of mice was executed prior to treatment according to the initial tumor volumes (not blinded to investigators).

In the investigation of GFPT2 pharmacological inhibition (***Fig.5***), the *in vivo* efficacy of Arbutin was initially assessed, an inhibitor specific to the GFPT2 isoform, previously identified and validated *in vitro.* Female C57BL/6N immunocompetent mice were introduced subcutaneously on the right hind flank with 5 × 10^5^ EGFR-mutated LLC. The mice were randomized into two groups of eight mice on the basis of tumor size. Randomization and treatment initiated on day 1; the mean tumor volume at the start of dosing was 50 mm^3^. SPSS (*0.9 % w/v saline, i.p.*) or Arbution (*50 mg/kg, i.p.*) was administered daily. Tumor volume was periodically assessed every two days using the formula *length × width*^2^ *× 0.5*. After 21 days treatment, mice were sacrificed and tumor mass ascertained. Through evaluation of Arbutin’s capacity to curb tumor progression, curtail CD8^+^T cell infiltration and activity within tumors, and modulate the HBP pathway in a tissue-specific manner, its potential as a therapeutic inhibitor was appraised. Subsequently, the combinatorial efficacy of Arbutin with PD-1 blockade was assayed. Female C57BL/6N immunocompetent mice harboring EGFR-mutated LLC tumors were assorted into distinct treatment regimens: (a) Control (*0.9 % w/v saline, i.p., qd*); (b) Arbutin (*50 mg/kg, i.p. qd*); (c) aPD-1 (*200 μg/mouse, i.p., qw*); (d) Arbutin (*50 mg/kg, i.p. qd*) + aPD-1 (*200 μg/mouse, i.p., qw*); (e) Bevacizumab (*5 mg/kg, i.p., tiw*) + aPD-1 (*200 μg/mouse, i.p., qw*). Tumor volumes were assessed every other day, until the survival end point was reached. Survival probability for each group was determined based on actual mortality observed within 77 days following xenograft implantation.

### Non-targeted metabolomics analysis

In the clinical sample assay, tissues weighing 0.1 g from patients were homogenized in 1 ml of 80 % (v/v) methanol, incorporating 1 μg/ml chlorophenylalanine as the internal standard (IS). For the cell line assay, BaF3 cells expressing multiple EGFR mutation variants were thrice rinsed with cold PBS and then collected. These cells were permeabilized in 1 ml of 80 % (v/v) methanol containing 1 μg/ml chlorophenylalanine as IS. For tumor TIF assay, 10 μl of TIF was permeabilized in a similar methanol-IS mixture. Following permeabilization, the tissue homogenates, cell lysates and TIF underwent centrifugation at 18,000 rpm for 5 minutes at 4 °C. The subsequent supernatants were carefully evaporated to dryness. These dried samples were then reconstituted in 100 μl of HPLC-grade water and centrifuged again under identical conditions. The resulting supernatants were then readied for subsequent analyses.

Metabolite analysis was performed using a Shimadzu LC −30A LC system (Kyoto, Japan) coupled with a TripleTOF 5600 system (AB SCIEX, Foster City, CA, USA). Chromatographic separation occurred on an Amide XBridge HPLC column (3.5 µm, 4.6 mm i.d. × 100 mm length; Waters, cat. no. 186004868) at 40 °C. The mobile phase comprised solvent A (95 % dd H_2_O and 5 % acetonitrile, supplemented with 5 mM ammonium acetate; pH 9 adjusted to 9 with 10 % (v/v) ammonia) and solvent B (pure acetonitrile). The elution flow rate was set at 0.4 ml/ min. Gradient elution unfolded as follows: 0 - 3 min, 85 % B; 3 - 6 min, 85 - 30 % B; 6 - 15 min, 30 - 2 % B; 15 - 18 min, 2 % B; 18 - 19 min, 2 - 85 % B; 19 - 26 min, 85 % B. The DuoSpray ion source, operating in electrospray ionization (ESI) mode and utilizing a negative ion mode for scanning, facilitated ionization of the metabolites. Mass spectrometry (MS) parameters were defined as follows: a time-of-flight MS scan ranging from m/z 50 - 1000 Da; ion source gas 1 and 2 set at 33 psi; a curtain gas at 25 psi; source temperature, 500 °C; ion spray voltage floating, −4500 V; declustering potential (DP), −93 V; and collision energy (CE), −10 V. The product ion scan ranged from m/z 50 - 1000 Da, maintaining a temperature of 500 °C and employing a Collision Energy Spread (CES) of 20, Ion Release Delay (IRD) of 67, and Ion Release Width (IRW) of 25. Calibration of accurate mass was facilitated by the Calibration Delivery System (AB SCIEX), with automatic recalibration occurring every five samples.

### Quantification of HBP metabolites and Adenosine using MS

Cells were subjected to washing with cold PBS, followed by three freeze-thaw cycles in 300 μl HPLC-grade water. Tissue samples underwent homogenization in 300 μl HPLC-grade water using a homogenizer. Both Plasma and TIF were diluted to predetermined ratios. Cell lysates and tissue homogenate (200 μl) or diluted plasma and TIF (30 μl) were precipitated using a solution comprising a four-fold volume of methanol and IS (5 μg/ml ^13^C-Glutamine for metabolites within HBP and 1 μg/ml chlorophenylalanine for Adenosine). Centrifugation was performed at 18,000 rpm for 10 min at 4 °C, post which, the supernatants were evaporated to dryness. The residues were reconstituted with 100 μl HPLC-grade water and underwent another centrifugation at 18,000 rpm for 5 min at 4 °C. The resultant supernatants were then prepared for analysis.

Metabolites within HBP were analyzed using a Shimadzu LC −30A liquid chromatography (LC) system (Kyoto, Japan) interfaced with a QTRAP 5500 system (AB SCIEX, Foster City, CA, USA). Chromatographic separation was achieved using an XSelect® HSS T3 (3.5 µm, 4.6 mm i.d. × 150 mm length, Waters, USA) at a maintained temperature of 40 °C. The mobile phase comprised solvent A (5 % acetonitrile and 5 mM ammonium acetate) and solvent B (acetonitrile), run at a flow rate of 0.5 ml/min. The gradient elution was programmed as follows:1 - 5 min, 5 % B; 5.5 min, 90 % B; 7.5 min, 90 % B; 8 min, 5 % B; 10 min, 5 % B. The mass spectrometer was operated in negative electrospray ionization (ESI) mode with a 4.5 kV capillary voltage. The multiple rection monitoring (MRM) parameters were set as follows: declustering potential set at −50 V for fructose-6P and glucosamine-6P, −90 V for UDP-GlcNAc, −60 V for ^13^C-glutamine (IS), −55 V for glutamine and - 50 V for GlcNAc-1P; collision energy set at −18 eV for fructose-6P, −25 eV for glucosamine-6P, −39 eV for UDP-GlcNAc, −14 eV for ^13^C-glutamine (IS), −14 eV for glutamine and −20 eV for GlcNAc-1P. MRM transition set as m/z 259.2 → 97 for fructose-6P, m/z 258.1 → 96.7 for glucosamine-6P, m/z 606.5 → 358.3 for UDP-GlcNAc, m/z 150.2 → 131.9 for ^13^C-glutamine (IS), m/z 145.1 → 127.1 for glutamine and m/z 300.3 → 97.2 for GlcNAc-1P.

Adenosine quantification was performed using a Shimadzu LC −30A LC system (Kyoto, Japan) interfaced with a QTRAP 6500 system (AB SCIEX, Foster City, CA, USA). Chromatographic separation was achieved using an Amide XBridge HPLC column (3.5 µm, 4.6 mm i.d. × 100 mm, Waters, cat. no. 186004868) at 40 °C. The mobile phase encompassed solvent A (5 % acetonitrile and 5 mM ammonium acetate) and solvent B (pure acetonitrile), delivered at a flow rate of 0.4 ml/min. The gradient elution program was: 1 - 2 min, 85 % B; 5 min, 20 % B; 8 min, 20 % B; 10 min, 85 % B; 16 min, 85 % B. For the mass spectrometric analysis, negative electrospray ionization (ESI) was utilized with a capillary voltage of 4.5 kV. The MRM parameters were set as follows: declustering potential set at −40 V for adenosine and chlorophenylalanine (IS); collision energy set at −20 eV for adenosine and −16 eV for chlorophenylalanine (IS). MRM transition set as m/z 268.4 → 136 for adenosine, m/z 200 → 154 for chlorophenylalanine (IS).

### Bulk RNA-seq

Total RNA was extracted from H1975 cells treated with Alm, EGF, and EGFR-knockdown using TRIzol® (Magen). Libraries for paired-end sequencing were prepared with the ABclonal mRNA-seq Lib Prep Kit, starting from 1 μg of total RNA, with mRNA isolated using oligo(dT) magnetic beads. Fragmentation and cDNA synthesis were performed according to the ABclonal protocol, followed by adapter ligation and PCR amplification. Libraries were purified using the AMPure XP system and quality-checked on an Agilent Bioanalyzer 4150. Sequencing on the Illumina NovaSeq 6000 platform produced 150 bp paired-end reads. Clean reads were aligned to the Mus musculus genome via HISAT2 (http://daehwankimlab.github.io/hisat2/), with gene read counts determined by Feature Counts and normalization to FPKM values. Differential expression analysis used DESeq2 (http://bioconductor.org/packages/release/bioc/html/DESeq2.html), identifying significant genes with | log2FC | > 1 and Padj < 0.05. Functional insights from differentially expressed genes were gained through GO enrichment analysis with the clusterProfiler package.

### Single-cell RNA-seq alignment and analysis

To investigate the TIME diversity in GFPT2-knockdown tumors, we conducted single-cell RNA sequencing (scRNA-seq) on a pooled sample of 24,777 cells derived from three EGFR-mutated LLC C57BL/6N xenografts, each treated with either shNC or shGFPT2 (n=3 per group). Tumor tissues were washed in PBS, minced, enzymatically dissociated with 0.1 % trypsin for 8 minutes, followed by 0.8 mg/ml collagenase type I for 60 minutes at 37 °C with 5 % CO_2_, including intermittent vigorous shaking. The cell suspension was filtered through a 70-μm strainer, red blood cells were lysed, and the remaining cells were suspended in PBS for single-cell 3’-cDNA library construction using the 10X Genomics Chromium Single Cell 3’ protocol. The encapsulation, cDNA synthesis, and subsequent RNA-sequencing were conducted by Gene Denovo, Guangzhou, China.

Single-cell RNA-seq libraries were sequenced on Illumina NovaSeq using 150 nt paired-end reads. Data processing was performed with Cell Ranger 4.0.0, adhering to default and recommended parameters. Reads were aligned to the Mus musculus reference genome (Ensembl_release109) using STAR. After initial processing with the Seurat package into Seurat Objects, mitochondrial gene representation was assessed for quality control. Cells were excluded based on the following criteria: mitochondrial gene content over 10%, presence of more than 7,000 genes (nFeature > 7000), or a total count above 100,000 (nCount > 100,000). Post-filtering, the dataset was normalized and subjected to Principal Component Analysis for dimensionality reduction. Integration of different samples was performed with Harmony (Broad Institute), and clustering was conducted using the t-SNE algorithm. Cluster-defining marker genes were identified, with the top 10 markers delineating each cluster. CellChat was used for cell-cell communication analysis.

### Analysis of tumor-infiltrating immune cells using flow cytometry

Tumor tissues were finely minced on ice using sterile forceps and scalpel. The resultant fragments underwent digestion with a mix of DNase I (10104159001, Sigma, USA) and Collagenase D (11088858001, Sigma, USA) at 37 °C for 45 min with agitation set at 220 rpm. Following digestion, the tissues suspensions were filtered through a 70 μm cell strainer and centrifuged at 1,000 rpm for 5 min. Cell pellets were resuspended with red blood cell lysis buffer (C3702, Beyotime, Shanghai, China) for 4 min on ice, followed by a subsequent 5 min centrifugation at 1000 rpm. Obtained single cells were then activated using a cell activation cocktail (423301, Biolegend) and cultured for 6 h at 37 °C, preparing for intracellular IFNγ and GZMB staining. Cell viability was assessed using the Zombie Aqua Fixable Viability Kit (423101, Biolegend, USA) and nonspecific binding was blocked with either Human TruStain FcX™ (422301, Biolegend, USA) or anti-mouse CD16/CD32 blocking antibody (101319, Biolegend, USA). Both surface and intracellular antigens were detected using a panel of fluorochrome-conjugated monoclonal antibodies. Gating strategies employed to identify humanized or murine tumor-infiltrating immune cells are presented in ***Supplementary Fig.7 and 8***.

For humanized tumor-infiltrating immune cells, cells were stained with the various combinations of fluorochrome-labelled antibodies in FACS buffer (PBS + 2 % FBS). **Panel 1:** GFP-labeling Tumor, APC-conjugated CD3 (1:100, 317317, Biolegend), APC/Cy7-conjugated CD8 (1:100, 344713, Biolegend), BV421-conjugated IFNγ (1:100, 502531, Biolegend). **Panel 2:** GFP-labeling Tumor, APC-conjugated CD3 (1:100, 317317, Biolegend), APC/Cy7-conjugated CD8 (1:100, 344713, Biolegend), BV421-conjugated GZMB (1:100, 396413, Biolegend). **Panel 3:** GFP-labeling Tumor, APC-conjugated CD3 (1:100, 317317, Biolegend), PerCP/Cy5.5-conjugated CD4 (1:100, 317428, Biolegend), APC/Cy7-conjugated CD25 (1:100, 302613, Biolegend), PE-conjugated FOXP3 (1:100, 320007, Biolegend). Cells were analyzed using a CytoFLEX II flow cytometer (Beckman Coulter) with FlowJo software.

For murine tumor-infiltrating immune cells, cells were stained with the various combinations of fluorochrome-labelled antibodies in FACS buffer (PBS + 2 % FBS). **Panel 1:** APC/750-conjugated CD45 (1:100, 147713, Biolegend), BV421-conjugated CD3 (1:100, 100335, Biolegend), PerCP/Cy5.5-conjugated CD8 (1:100, 100733, Biolegend), BV605-conjugated IFNγ (1:100, 505840, Biolegend), APC-conjugated GZMB (1:100, 372204, Biolegend); **Panel 2:** BV421-conjugated CD45 (1:100, 103133, Biolegend), PE/Cy7-conjugated CD3 (1:100, 100219, Biolegend), BV605-conjugated CD4 (1:100, 100547, Biolegend), Alexa 647-conjugated CD25 (1:100, 302613, Biolegend), PE-conjugated FOXP3 (1:100, 320007, Biolegend); **Panel 3:** BV421-conjugated CD45 (1:100, 103133, Biolegend), PE/Cy7 conjugated F4/80 (1:100, 123113, Biolegend), BV605-conjugated CD11b (1:100, 101237, Biolegend), PE-conjugated CD206 (1:100, 141705, Biolegend), APC-conjugated CD86 (1:100, 105011, Biolegend); **Panel 4:** APC/750-conjugated CD45 (1:100, 147713, Biolegend), BV421-conjugated CD3 (1:100, 100335, Biolegend), Pacific Blue-conjugated CD19 (1:100, 115526, Biolegend), PE-conjugated NK1.1 (1:100, 108707, Biolegend). Cells were analyzed using a CytoFLEX II flow cytometer (Beckman Coulter) with FlowJo software.

### N-glycoproteomes

Proteins in the samples were quantified using the BCA protein assay. Subsequently, 1 mg of protein from each condition was aliquoted into a fresh Eppendorf tube, and its volume was brought to 200 μl using 8 M urea. The samples were then reduced by the addition of 20 μl of 0.5 M TCEP and incubated at 37 °C for 1 h. Alkylation was achieved by adding 40 μl of 1 M iodoacetamide and incubating in the dark at room temperature for 40 min. Proteins were precipitated by the addition of five volumes of pre-chilled acetone at −20 °C overnight. The resulting precipitates were rinsed twice with 1 ml of 90 % pre-chilled acetone and subsequently solubilized in 1 ml of 100 mM TEAB. Digestion was carried out using sequence-grade modified trypsin (Promega, Madison, WI) at an enzyme-to-protein ratio of 1:50 (w/w) and incubated at 37 °C overnight. The resulting peptides were purified using C18 ZipTips, quantified via the Pierce™ Quantitative Colorimetric Peptide Assay (23275, Thermo Scientific), and lyophilized using SpeedVac. The lyophilized peptides were resolubilized in 80 % acetone with 1 % TFA and loaded onto an ihouse ZIC-HILIC micro-column containing 30 mg of ZIC-HILIC particles (Merck Millipore, Merck KGaA, Darmstadt, Germany) with a C8 disk. The eluate was recycled over the column five times in total. After washing the column with 80 % acetone and 1 % TFA, enriched glycopeptides were sequentially eluted using 0.1 % TFA, 25 mM NH_4_HCO_3_ and 50 % acetone, then dried via vacuum centrifugation. These peptides were subsequently resuspended in 10 µl of solvent A (solvent A: 0.1 % formic acid in H_2_O) for analysis on an Orbitrap Fusion Lumos Tribrid mass spectrometer (Thermo Scientific, MA, USA) coupled to an EASY-nano-LC 1200 system (Thermo Scientific, MA, USA) without the trap column. 3 µl of the peptide sample was loaded onto a C18 spray tip column (15 cm × 75 μm i.d., Acclaim PepMap) and eluted over a 120 minutes gradient from 3 % to 32 % solvent B (solvent B: 0.1 % formic acid in ACN) at 300 nl/min. The electrospray voltage of 2 kV was applied relative to the mass spectrometer’s inlet. The mass spectrometer was run under data dependent acquisition mode, and automatically switched between MS and MS/MS mode. The parameters were MS: scan range (m / z) = 400 - 2000; resolution = 120,000; AGC target = 800,000; maximum injection time = 100 ms; included charge state = 2 – 6; dynamic exclusion after n times, n = 1; dynamic exclusion duration = 30 s; each selected precursor was subject to one HCD-MS/MS; HCD-MS/MS: isolation window = 4; detector type = Orbitrap; resolution = 15,000; AGC target = 200,000; maximum injection time = 22 ms; collision energy = 30 %; stepped collision mode on, energy difference of ± 10 % (10 % as absolute value in the Orbitrap Fusion). Tandem mass spectra were analyzed using by pGlyco 3.0 from pFind Studio. Searches were conducted against the UniProt-homo_sapiens protein database (ver.2022, 20589 entries) and the human-specific glycan database (pGlyco, species: human) assuming the digestion enzyme Trypsin. The search parameters for pGlyco 3.0 included a fragment ion mass tolerance of 20 ppm and a parent ion tolerance of 10 ppm. Carbamidomethylation (C) was set as a fixed modification, while oxidation (M) was considered a variable modification. Glycopeptide identifications were filtered to achieve a 1 % total FDR. For quantitative analysis of site-specific glycosylation, the pGlycoQuant tool within pFind Studio was employed using label-free quantification.

### Western blotting

Whole-cell lysates were prepared from cultured cells using NP40 Lysis Buffer and from tumor samples using RIPA Lysis Buffer. Specifically, cultured cells were rinsed thrice with cold PBS, followed by lysis on ice using NP40 buffer containing 100 μM Phenylmethanesulfonylfluoride and 0.1 % (v/v) phosphatase inhibitor (Beyotime Biotechnology, China) for 30 min. Tumor samples were homogenized in RIPA buffer supplemented with inhibitors and incubated on ice for 40 min. Post lysis, protein supernatants were acquired via centrifugation at 14,000 g for 5 min at 4 °C. Surface and cytoplasmic proteins were isolated using the Surface and Cytoplasmic Protein Reagent Kit (P0033, Beyotime, Shanghai, China), following the provided manufacturer’s protocol. In summary, cells and tumor samples were suspended and lysed in 1 ml of Isolation Solution A, supplemented with 1 % PMSF, and then homogenized on ice. The resulting lysates were centrifuged at 700 g for 10 min to remove nuclear. The collected supernatants were further centrifuged at 14,000 g for 30 min. The pellets were resuspended in Extraction Buffer B and lysed for an additional 20 min. Following a final centrifugation at 14,000 g for 5 min, the supernatant was designated as the membranous protein lysate. All protein samples had their concentrations assessed using the bicinchoninic acid (BCA) Protein Assay kit (Beyotime Biotechnology, China). These samples were denatured at 100 °C in 4 × XT Sample Buffer (Bio-Rad, USA). N-glycosylation modification of proteins was assessed using the PNGase F Glycan Cleavage Kit (A39245, Gibco, USA). In brief, protein lysates were standardized to a concentration of 5 μg/μl. A volume of 20 μl of this lysate was mixed with 4.5 μl of 10 × Glycoprotein Denaturing Buffer, 18.5 μl of ultrapure water, and 2 μl of PNGase F, achieving a total reaction volume of 45 μl. This blend was then incubated at 50 °C for 1 h to facilitate the removal of N-linked oligosaccharides. Subsequently, 15 μl of the loading buffer was incorporated and the mixture was denatured by heating at 100 °C for 5 min.

Protein lysates (30 - 50 mg) were loaded onto 4 - 12 % Omni-PAGE™ Hepes Plus gels (Epizyme Biomedical Technology, China). After separation, protein was transferred to a PVDF membrane (Bio-Rad). Membranes were blocked in 5 % skim milk in TBST (50 mM Tris–HCl, 150 mM NaCl, 0.05 % Tween 20) for 2 hours at room temperature, followed by an overnight incubation 4 °C with primary antibodies (See below). After three washes with TBST, membranes were incubated with HRP-conjugated secondary antibodies (1:10000 dilution) for 2 hours at room temperature. Following another three washes, blots visualization was achieved using an enhanced chemiluminescence detection kit (Bio-Rad, USA) and images were captured using a ChemiDoc XRS-System (Bio-Rad, USA). Signal intensities were normalized to GAPDH or Actin. The intensity of the selected band was analyzed using ImageJ. The following antibodies were used: anti-GFPT1 (ab125069), anti-GFPT2 (ab190966), anti-EGFR (ab52894), anti-pEGFR (phosphoY1068, ab40815), anti-ERK (ab184699), anti-pERK (phospho T202 for ERK1 and phospho T185 for ERK2, ab201015), anti-AKT (ab179463), anti-pAKT (phospho T308, ab38449), anti-pIRE1α (ab124945), anti-Xbp1s (ab220783), anti-PD-L1(ab213524), anti-CD276 (ab134161), anti-PVR (ab103630), anti-CD73 (ab133582), anti-Na^+^/K^+^ ATPase (ab76020, 1:10000) (others were used at 1:1,000 and all from Abcam); anti-IRE1α (3294) (was used at 1:1,000 and from Cell Signaling Technology); anti-Actin (AP0731), anti-GAPDH (AP0063) (both were used at 1:1,000 and from Bioworld); Goat pAb to Rabbit IgG HRP (ab6721), Goat pAb to Mouse IgG HRP (ab205719) (both were used at 1:10,000 and from Abcam).

### RNA isolation and RT-qPCR

For the isolation and purification of RNA from diverse cell types and tumor specimens, the RNA Isolation Kit (RNAiso Plus, Takara, Japan) was employed, followed by reverse transcription into cDNA utilizing the PrimeScript RT Regent Kit (Vazyme, Nanjing, China) in alignment with the guidelines provided by the manufacturer. Real-Time Quantitative Polymerase Chain Reaction (RT-qPCR) assays were conducted on the CFX96 real-time detection system (Bio-Rad, USA) with the SYBR Green PCR Mix (Bio-Rad, USA). The primers for detection are listed below: *GFPT2-*F ATGTGCGGAATCTTTGCCTAC, *GFPT2-*R ATCGAGAGCCTTGACTTTCCC; *CXCL10*-F GAAATTATTCCTGCAAGCCAATTT, *CXCL10*-R TCACCCTTCTTTTTCATTGTAGCA; *VEGF*-F CGACGGCTTGGGGAGATTGC, *VEGF*-R GGGCGGTGTCTGTCTGTCTG; *ACTIN-F* GCGTGACATTAAGGAGAAG, *ACTIN-R* GAAGGAAGGCTGGAAGAG. PCR cycling parameters were set to an initial denaturation at 95 °C for 10 min, subsequently followed by 40 cycles of 95 °C for 15 s, annealing at 60 °C for 30 s, and an extension phase at 72 °C for 30 s. To confirm the specificity of the PCR products, a melting curve analysis was consistently performed post amplification. The *ACTIN* served as the internal reference. Quantitative evaluations of target gene mRNA expression were derived utilizing the 2^−ΔΔCt^ methodology, where cycle threshold (Ct) values were normalized to *ACTIN*.

### Chromatin Immunoprecipitation

Chromatin immunoprecipitation (ChIP) assay was conducted using the Pierce™ Magna ChIP assay kit (26157, ThermoFisher) as per the manufacturer’s instructions. Briefly, 1 × 10^7^ cells were subjected to cross-linking using 1 % formaldehyde (w/v) for a duration spanning10 - 15 min, followed by the addition of 0.125 M glycine to terminate the cross-linking. Subsequent to the utilization of a nuclear and cytoplasmic protein extraction kit (P0027, Beyotime) for nuclear isolation, chromatin DNA was sonicated, producing fragments averaging between 200 - 500 bp in length. The resultant soluble chromatin was subjected to overnight immunoprecipitation at 4 °C using either anti-Xbp1s or control anti-IgG antibodies, employing gentle agitation. Protein A/G beads were incorporated to capture the chromatin-antibody complex, allowing for a 4 h incubation at 4 °C with mild agitation. DNA-protein complexes were subsequently liberated from the beads via elution buffer, subjected to a 30 min incubation at 65 °C under vigorous shaking conditions. Capturing the supernatant with a magnetic stand, the eluted mixture was combined with 5 M NaCl (6 μl) and 20 mg/ml Proteinase K (2 μl), and incubated at 65 °C for 1.5 h. DNA purification was achieved utilizing DNA binding buffer, DNA Clean-Up column, and DNA washing buffer, and the purified DNA was then quantified via qPCR. The primers for detection are listed below: *GFPT2 promoter 1-F* CCTCCTTCCTGGCTTCGCTA, *GFPT2 promoter 1-R* AGGATACAAGGAGGGGGCTG; *GFPT2 promoter 2-F* CAGGCCATCAACCTCCCAGA, *GFPT2 promoter 2-R* ACAGGTCCACGGTGAGGAAC.

### Normalized growth rate inhibition and Clonogenic assay

H1975 and PC9/OR cells, with stable transfections of either GFPT2 knockdown or an empty vector, were seeded in 96-well plates at a density of 10,000 cells per well. Subsequent to a 24-hour incubation, cell numbers between the two groups were compared utilizing a CCK-8 assay. For drug treatments assays, H1975 cells were seeded in 96-well plates at a density of 10,000 cells per well and allowed to adhere for 24 h preceding drug exposure. Cells were then treated with a range of AZA concentrations (0.001, 0.01, 0.1, 1, 2 and 5 μM) or Arbutin (0.01, 0.1, 1, 10, 100, 1000 and 5000 μM) for 48 h. In parallel, PC9, PC9/OR1 and PC9/OR2 cells were plated similarly and treated with various concentrations of Osimertinib (0.01, 0.1, 1, 5 μM) for 48 h. For both sets of treatments, cell viability post-drug exposure was quantified using the CCK-8 assay. Absorbance readings were normalized to DMSO-treated controls on the same plates, providing normalized growth rate inhibition values.

To delve deeper into the potential influence of GFPT2 expression on tumorigenic capacity, clonogenic assays were conducted. Either wild-type cells or those with GFPT2 knockdown were plated at a density of 500 cells/well in 6-well plates. The culture medium was refreshed every alternate day. After a 7-day incubation period, cells were fixed utilizing 4 % paraformaldehyde (w/v) for 10 min and subsequently stained with 20 % (v/v) crystal violet for 15 min. After washing off excess dye, cell colonies were visualized, captured and quantified.

### Immunofluorescence

Cells were seeded at a density of 10,000 cells per well in a 4-chamber confocal dish and, post-treatment, were fixed with 4 % paraformaldehyde for 10 min at room temperature following different treatment. Xenograft tumor tissues underwent fixation in 4 % paraformaldehyde overnight, sequentially dehydrated in 20 % and 30 % sucrose solution, and embedded in Tissue-Tek OCT compound (4583, Sakura). Sections of 10 μm thickness were prepared using a Leica microtome. Overall N-glycosylation was assessed by incubation with anti-Phaseolus Vulgaris Leucoagglutinin (PHA)-Rhodamine (1:200, RL-1112, Vector) for 1 h at room temperature. Specific immune checkpoint ligands and pEGFR glycosylation were probed using primary antibodies against PD-L1 (1:100, ab213524, Abcam), CD276 (1:200, ab134161, Abcam), PVR (1:200, ab103630, Abcam), CD73 (1:200, ab133582, Abcam) and pEGFR (1:500, ab40815, Abcam) overnight at 4 °C, followed by a 1 h incubation with anti-PHA (1:200, RL-1112, Vector). Samples were subsequently treated with Cy2-conjugated anti-rabbit secondary antibody (1:200,111-225-144, Jackson ImmunoResearch) for 2 h at room temperature in the dark. The PHA^+^ target protein intensity and ratio were analyzed using Image J. For PD-L1/PD-1 binding assessment, samples were treated with recombinant human PD-1 Fc protein (1086-PD-050, R&D System) at 4 °C overnight, followed by incubation with an anti-human Alexa Fluor 488 conjugate (1:200, A-10631, Thermo Fisher). For comparing tumor cell growth in different groups, tumor sections in a cryostat were incubated with either anti-Ki67 (1:200, ab15580, Abcam) or anti-cleaved CASP3 (1:200, ab32042, Abcam), along with anti-PanCK (1:200, ab7753, Abcam) primary antibody at 4 °C overnight, followed by a 2 h exposure to Alexa Fluor 555-conjugated anti-rabbit secondary antibody (1:200, A0453, Beyotime) and Cy5-conjugated anti-mouse secondary antibody (1:200, 15-175-166, Jackson ImmunoResearch), shielded from light. PanCK^+^Ki67^+^ and PanCK^+^ CASP3^+^ subpopulation ratios were determined by Image J. All samples received a 10 min DAPI counterstain (C1006, Beyotime) in the dark and then were mounted with Fluoromount-G® (Southern Biotech, 0100-20). Imaging was conducted on an Olympus FV3000 confocal microscope using either 60 X or 100 X objectives.

### ELISA

A 30 mg portion of tumor tissue was homogenized in 300 μl of deionized water, followed by a 1:50 dilution in assay diluent. TIF and cell culture supernatant were harvested post various treatments. The levels of VEGF, CCL5, CXCL9, CXCL10, CXCL11, IFNγ, IL-2 and GZMB in the samples were quantified using the respective ELISA kits: VEGF (EH015, ExCell), CCL5 (CSB-E17375h, CUSABIO), CXCL9 (CSB-E09024h, CUSABIO), CXCL10 (CSB-E08181h, CUSABIO), CXCL11 (CSB-E09023h, CUSABIO), IFNγ (EH005, ExCell), IL-2 (E-EL-H0099, Elabscience), and GZMB (E-EL-H1617c, Elabscience). Procedures followed the manufacturer’s guidelines. The absorbance was recorded using Biotek Synergy H1 microplate reader.

### MicroScale Thermophoresis assay

293T cells were engineered to ectopically express either GFP-GFPT1 or GFP-GFPT2. Cell homogenates were obtained and adjusted to predetermined concentrations. These preparations were then subjected to incubation with six identified compounds across a range of concentrations for 15 min. The mixtures were introduced into NanoTemper glass capillaries, and microthermophoresis was conducted using the Monolith NT.115 instrument (NanoTemper) in strict adherence to the manufacturer’s guidelines. Dissociation constants (Kd values) were computed employing the mass action equation, utilizing the NanoTemper software, and were grounded in duplicate read measurements.

### Public bioinformatics dataset analysis

The TCGA data set called Lung Adenocarcinoma (TCGA, PanCancer Atlas) was downloaded from the TCGA data portal (https://tcga-data.nci.nih.gov/tcga/). The correlations between the expression of HBP metabolites and survival in EGFR-mutated NSCLC were provided by Kaplan-Meier Plotter (http://kmplot.com/analysis/). The human proteome map (http://www.humanproteomemap.org/query.php) was used to describe the expressions of HBP metabolic enzymes. The nucleotide sequences in the GFPT2 promoter which Xbp1s binding to were predicted with JASPER 2022 (https://jaspar.genereg.net/). Gene ontology (GO) enrichment analysis was carried out with THE GENE ONTOLOGY RESOURCE (https://geneontology.org/). The poor prognosis of PD-1 inhibitors was obtained from timer 2.0 (http://timer.cistrome.org).

### Data availability

The authors declare that the data supporting the findings of this study are available within the paper and its Supplementary Information. Transcriptomic data for scRNA-seq have been deposited at the Gene Expression Omnibus (GEO) under accession numbers GSE259244. Source data are provided with this paper. All R code used to generate figures in the manuscript can be made available upon reasonable request.

## Acknowledgments

We acknowledge the financial support of the Natural Science Foundation of Jiangsu Province (BK20220149 and BK20192005, China), Young Elite Scientists Sponsorship Program by China Association for Science and Technology (2023QNRC001), the National Natural Science Foundation of China (Nos.82173882, 82173890, 82073929), Haihe Laboratory of Cell Ecosystem Innovation Fund (HH22KYZX0006).

## Author contributions

Jiali Liu conceptualized and supervised the project. Luyao Ao and Wenjing Jia performed the practical work. Jiawen Cui analyzed and uploaded the sc-RNA sequence data. Qixing Gong and Jia Wei designed and analyzed IHC staining of clinical specimens. Shencun Fang, Jun Wang and Chenghao Fu provided clinical specimens. Haobin Li performed Schrödinger molecular docking. Ying Yu, Ruiqi Wang, Feiyi Wang, Xin Shang, Yantong Li participated in assistance work. Jiali Liu, Luyao Ao, Wenjing Jia wrote the manuscript with the input from Fang Zhou and Guangji Wang.

## Additional information

Supplementary materials included supplementary Fig.1 to Fig. 16 and supplementary Table.1 to Table.13.

---

**Extended Data Fig.1.**
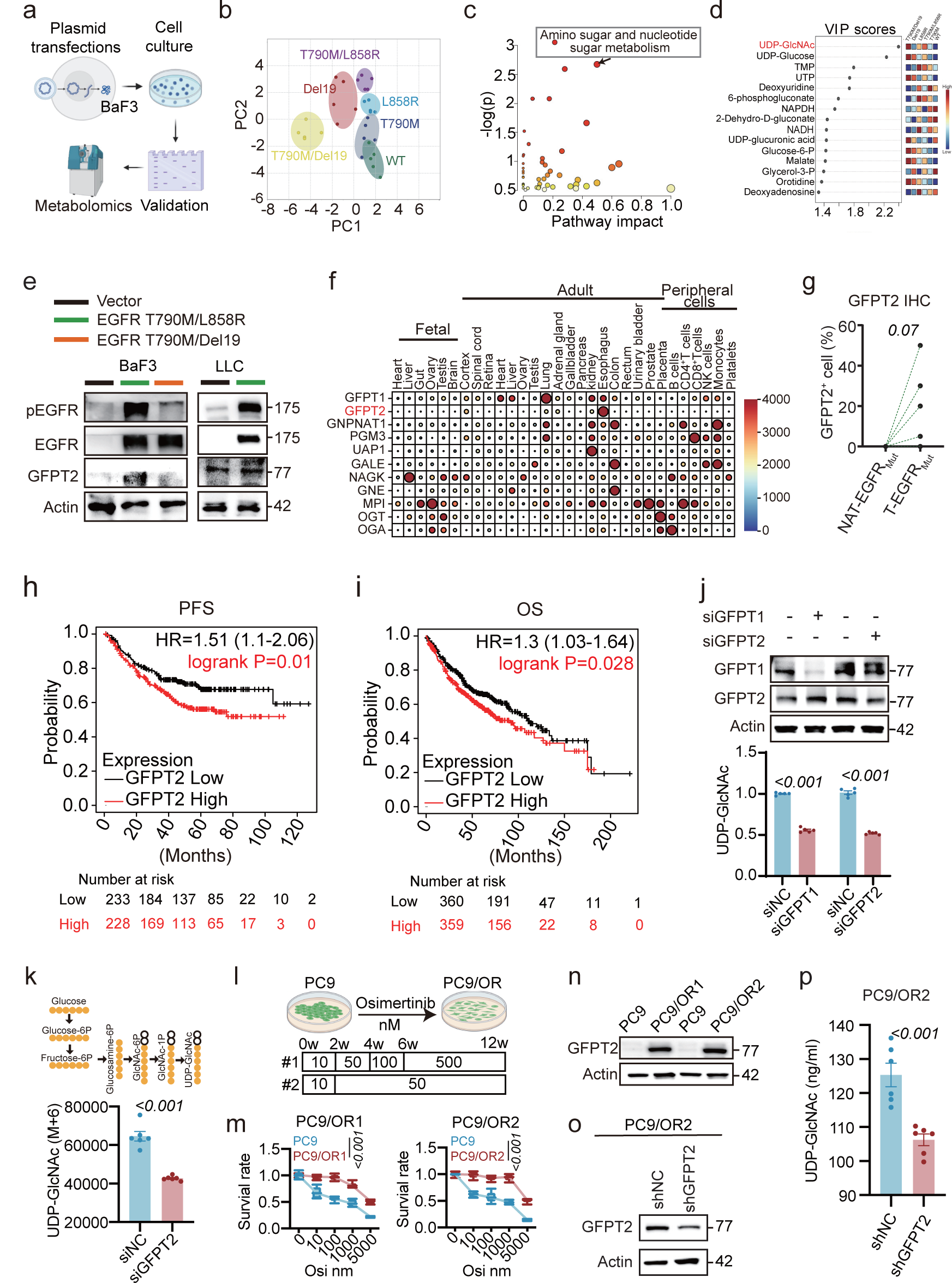
Elevated GFPT2 expression in EGFR_Mut_ NSCLC (Related to Fig.1) **a-d**, Metabolomics in BaF3 cells with various EGFR mutations. (**a**) Experimental setup schematic, (**b**) PCA of metabolite profiles, (**c**) Metabolic pathway impact plot detailing significance and impact, and (**d**) VIP scores for leading metabolites. **e**, Immunoblots of pEGFR, EGFR, and GFPT2 in BaF3, LLC cells transfected with EGFR_Mut_ plasmids, with Actin as a loading control. **f**, Heatmap of HBP metabolic enzymes across primary organs and immune cells in both adult and fetal stages, sourced from the human proteome map (http://www.humanproteomemap.org/query.php). **g**, Quantifications of GFPT2 IHC staining. **h-i**, Progression free survival (PFS) and overall survival (OS) of highly or lowly expressed GFPT2-patients suffering LUAD according to Kaplan-Meier Plotter (http://kmplot.com/analysis/). **j**, UDP-GlcNAc concentrations in H1975 cells following GFPT1 or GFPT2 knockdown, measured by HPLC-QTRAP/MS (n=5). **k**, UDP-GlcNAc (M+6) levels in siNC and siGFPT2 H1975 cells cultured with U-^13^C_6_ D-glucose (n=6). **l**, Generation of PC9/Osimertinib-resistant (PC9/OR) cells via gradual Osimertinib exposure. **m**, Viability of PC9/OR cells upon various Osimertinib concentrations (n=6). **n-o**, Immunoblots of GFPT2 expression in PC9 versus PC9/OR cells, with Actin as the loading control. **p**, HPLC-QTRAP/MS analysis of UDP-GlcNAc levels in PC9/OR2 cells (n=6). Data are mean ± s.e.m. except in **g**, Two-tailed paired Student’s t-test. **j**, **k**, **p**, Two-tailed unpaired Student’s t-test. **m**, Two-way ANOVA.

**Extended Data Fig.2.**
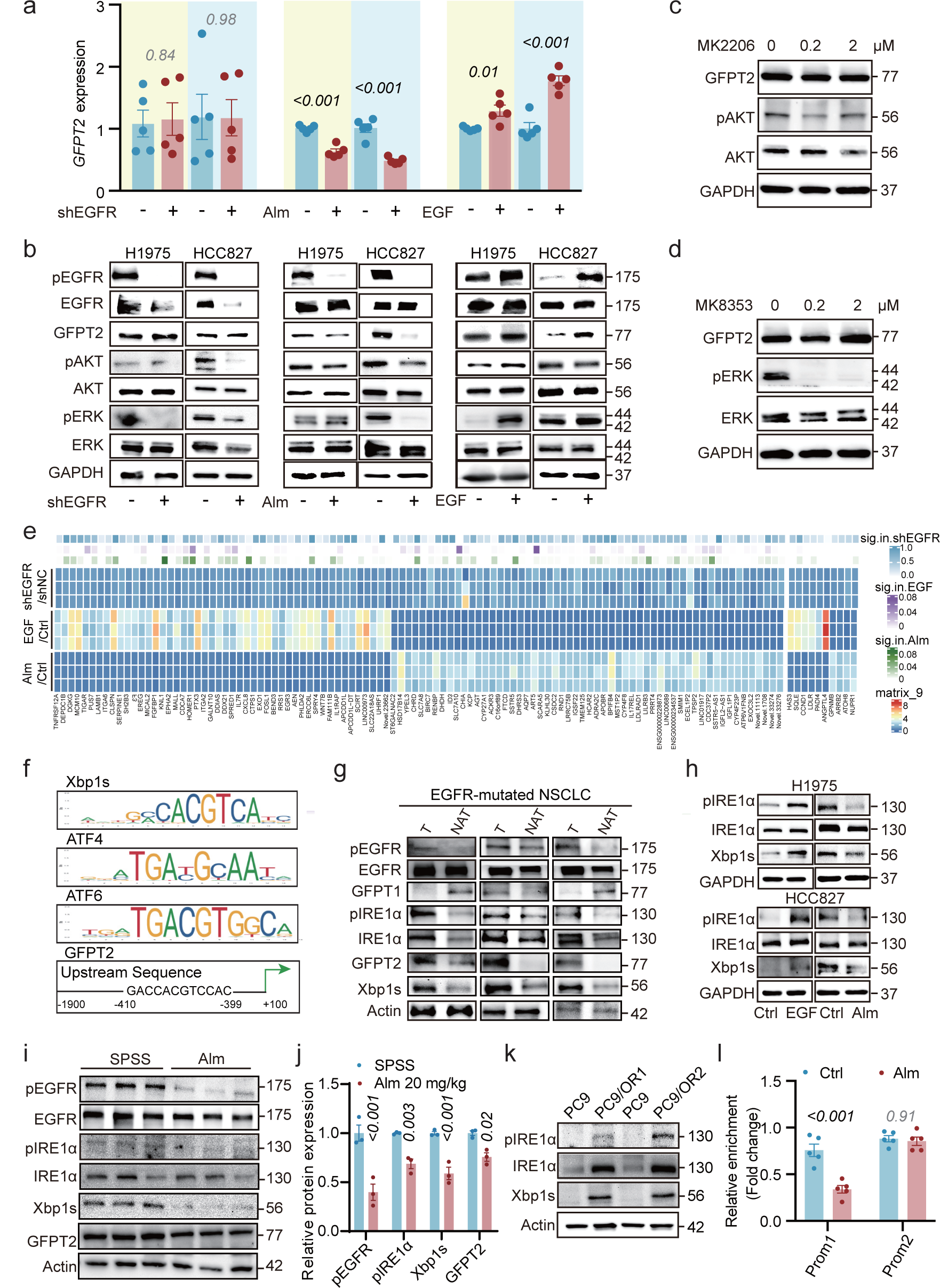
Regulation of GFPT2 via the EGFR/IRE1α/Xbp1s pathway in EGFR_Mut_ NSCLC (Related to Fig.1) **a**, GFPT2 mRNA levels in H1975 and HCC827 cells post-treatment with Alm, EGF, and shEGFR interventions (n=5). **b**, Immunoblots of pEGFR, EGFR, phosphorylated AKT (pAKT), AKT, phosphorylated ERK (pERK), ERK, and GFPT2 in H1975 and HCC827 cells under the same treatments as in **a**, with GAPDH as a loading control. **c**, **d**, Immunoblots of GFPT2, pAKT, and AKT in H1975 cells treated with AKT inhibitor MK2206 or ERK inhibitor MK8353. **e**, RNA-seq based identification of differentially expressed genes in H1975 cells treated with EGF or Alm, compared to EGFR knockdown cells (n=3). **f**, Prediction of transcription factor binding sites involved in ERS signaling pathways on GFPT2 promoter region (2000 bp upstream) by Jasper website, highlighting sequences similar to canonical Xbp1s binding sites. **g**, Immunoblots comparing ER stress-related proteins in T versus NAT from EGFR-mutated NSCLC patients, with Actin as a loading control (n=3). **h**, Western blots of IRE1α, pIRE1α, Xbp1s in H1975 and HCC827 cells treated with EGF and Alm, with GAPDH as loading control. **i-j**, Analysis of pEGFR, EGFR, IRE1α, p-IRE1α, Xbp1s, and GFPT2 in H1975 xenograft tumors in Balb/c nude mice post-Alm treatment, n=3, with Actin as loading control. **k**, Protein level comparison of IRE1α, pIRE1α, Xbp1s, and GFPT2 in PC9 and PC9/OR cells by Western blot, with Actin as loading control. **l**, ChIP-PCR quantification of Xbp1s binding to GFPT2 promoters in Alm-treated H1975 cells (n=5). Data are mean ± s.e.m. **a**, One-way ANOVA. **j**, Two-way ANOVA. **l**, Two-tailed unpaired Student’s t-test.

**Extended Data Fig.3.**
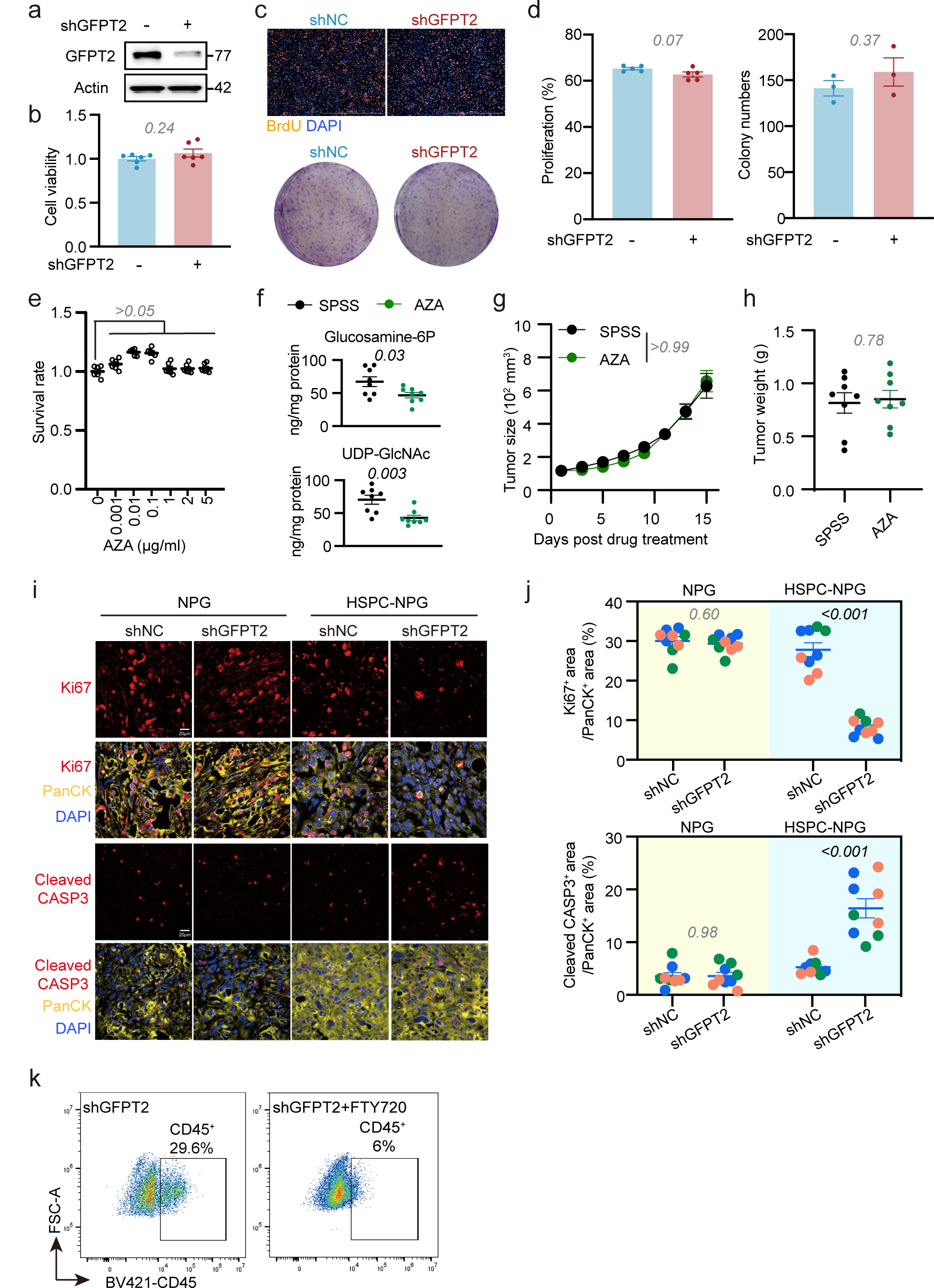
Effects of GFPT2 inhibition on tumor growth (Related to Fig.2) **a**, Verification of GFPT2 knockdown stability via blotting, with Actin as loading control. **b**, Cell viability post GFPT2 knockdown (n=6). **c-d**, BrdU staining (n=5) and colony formation assay (n=3) depicting proliferation in shNC and shGFPT2 H1975 cells. **e**, Survival rates of H1975 cells exposed to gradient concentrations of Azaserine (AZA) (n=6). **f**, HBP metabolite levels in tumor homogenates, assessed by HPLC-QTRAP/MS after two-week AZA administration (n=8). **g**, **h**, Tumor size and weight of H1975 xenograft in Balb/c nude mice following AZA treatment (n=8). **i-j**, Immunofluorescent detection of Ki67 and cleaved CASP3 in H1975 xenograft sections (n=3, three field each). **k**, CD45^+^cell amount in EGFR-mutated LLC xenograft post FTY720 treatment. Data are mean ± s.e.m. **b**, **d**, **f**, **h**, **j** Two-tailed unpaired Student’s t-test. **e**, One-way ANOVA. **g**, Two-way ANOVA for tumor size.

**Extended Data Fig.4.**
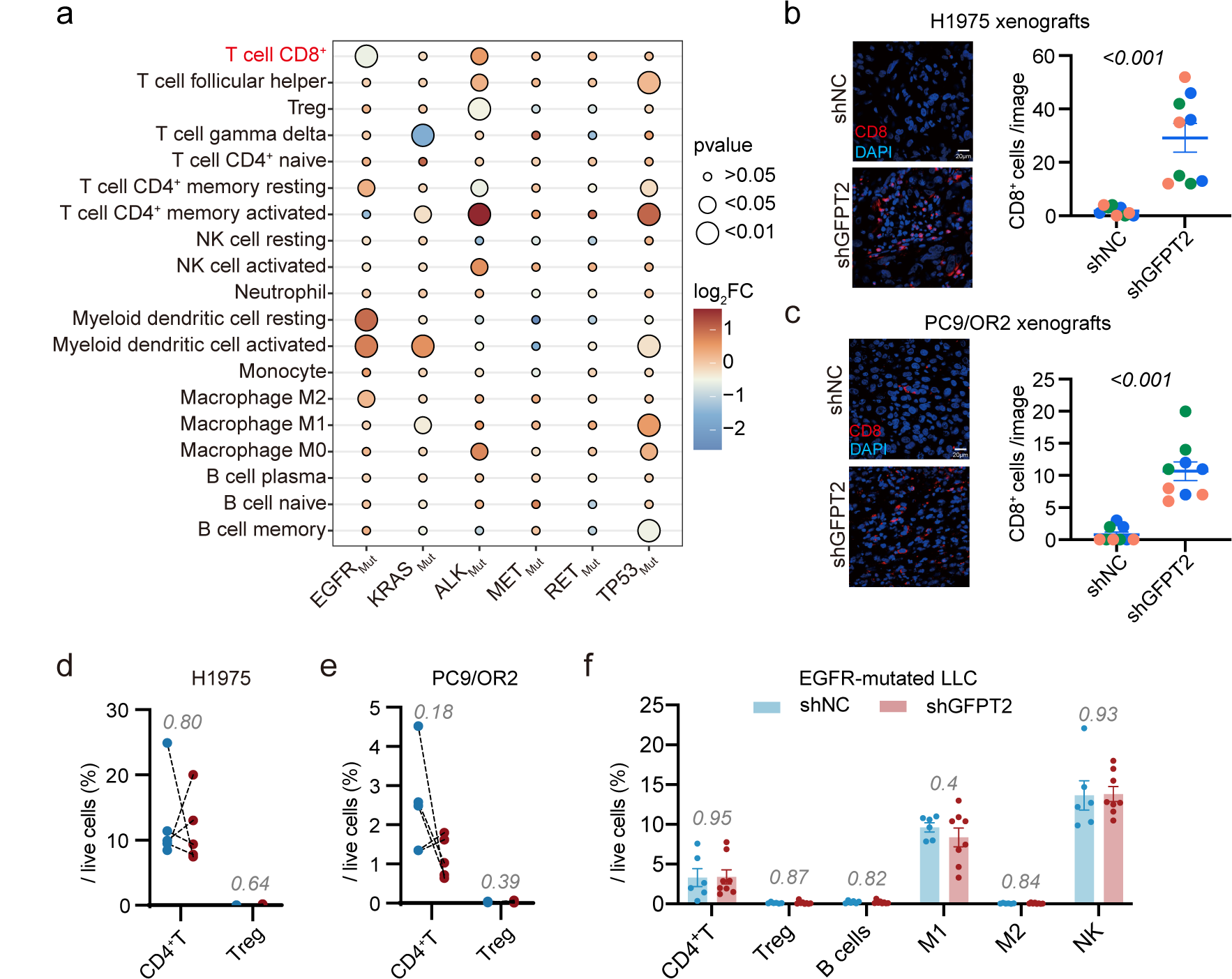
Effects of GFPT2 inhibition on tumor immune microenvironment (Related to Fig.2) **a**, Tumor immune microenvironment in common oncogene mutation in LUAD according to TIMER2.0 (http://timer.cistrome.org/) with CIBERSORT Abs. mode. **b**, **c**, Immunofluorescent detection of CD8α in H1975 and PC9/OR2 xenograft sections (n=3, three field each). **d-e**, CD4^+^T cells and Tregs in HSPC-NPG mice xenografts from H1975 or PC9/OR2 cells post GFPT2 silenced were examined via FCM (n=5). **f**, Immune cells in C57BL/6N mice xenograft from EGFR-mutated LLC cells after GFPT2 knockdown (n=6 for shNC and n=8 for shGFPT2). Data are mean ± s.e.m. except in **d**, **e**, Two-tailed paired Student’s t-test. **b**, **c**, **f**, Two-tailed unpaired Student’s t-test.

**Extended Data Fig.5.**
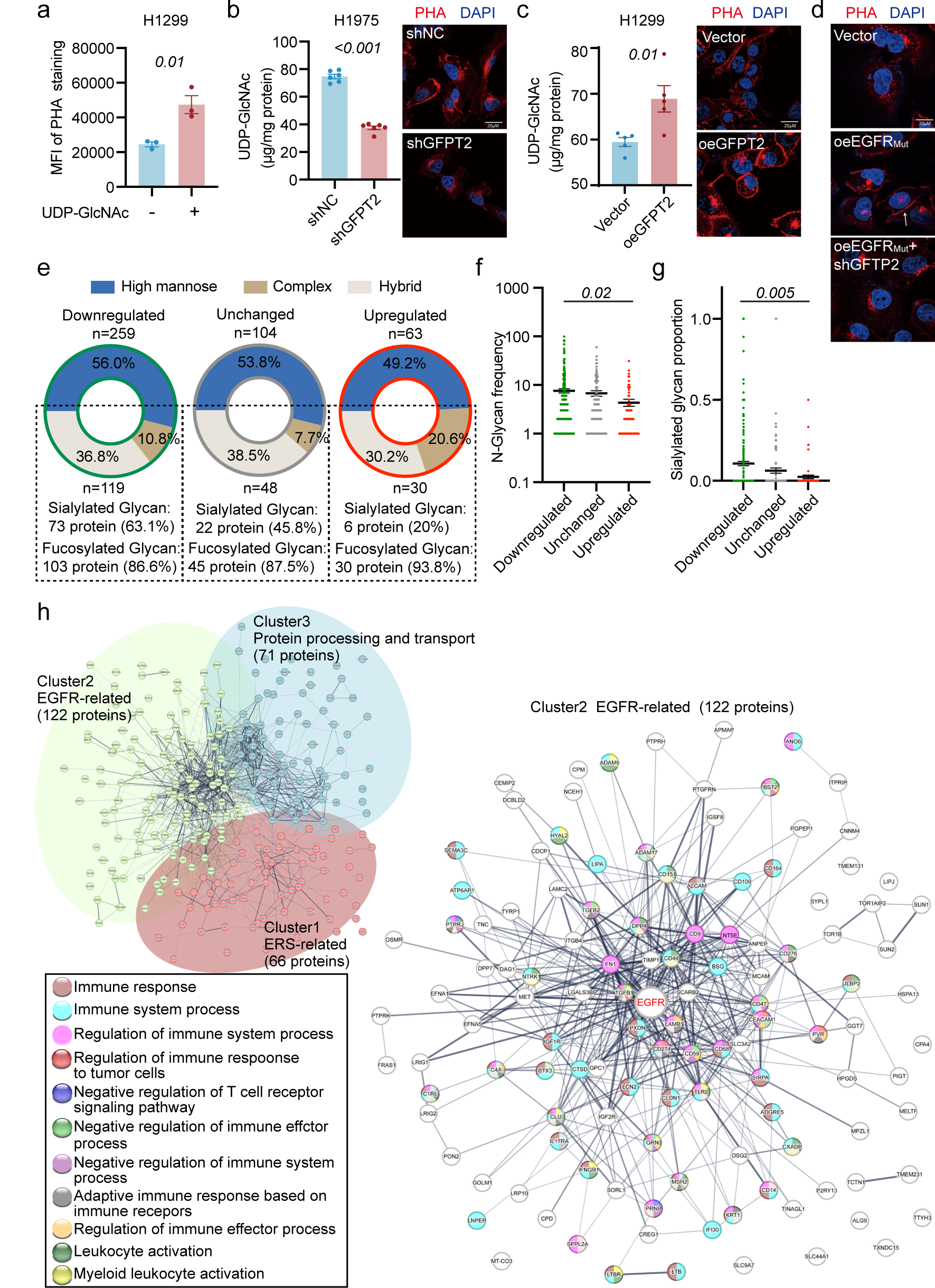
N-glycoproteomes post genetic disruption of tumoral GFPT2. **a**, FCM analysis depicting mean fluorescent intensity (MFI) of PHA staining in H1299 cells treated with UDP-GlcNAc (10 mM) (n=3). **b**, **c**, LC-QTRP/MS analysis illustrating UDP-GlcNAc levels and PHA staining of protein glycan coat in H1975 (n=6) and H1299 (n=5) cells. **d**, Representative images of shNC and shGFPT2 H1299 cells following the induction of oncogenic EGFR mutation (T790M/L858R) subjected to immunofluorescent staining for PHA. **e**, GFPT2-mediated alterations in 426 protein glycoforms, as revealed by N-glycoproteomic analysis. **f**, **g**, Changes in glycan occurrence and proportion of sialylated glycan on identified proteins. **h**, Protein exhibiting reduced glycosylation segregated by a K-means clustering algorithm into three discrete cohorts. Data are mean ± s.e.m. **a**, **b**, **c**, Two-tailed unpaired Student’s t-test. **f**, **g**, One-way ANOVA.

**Extended Data Fig.6.**
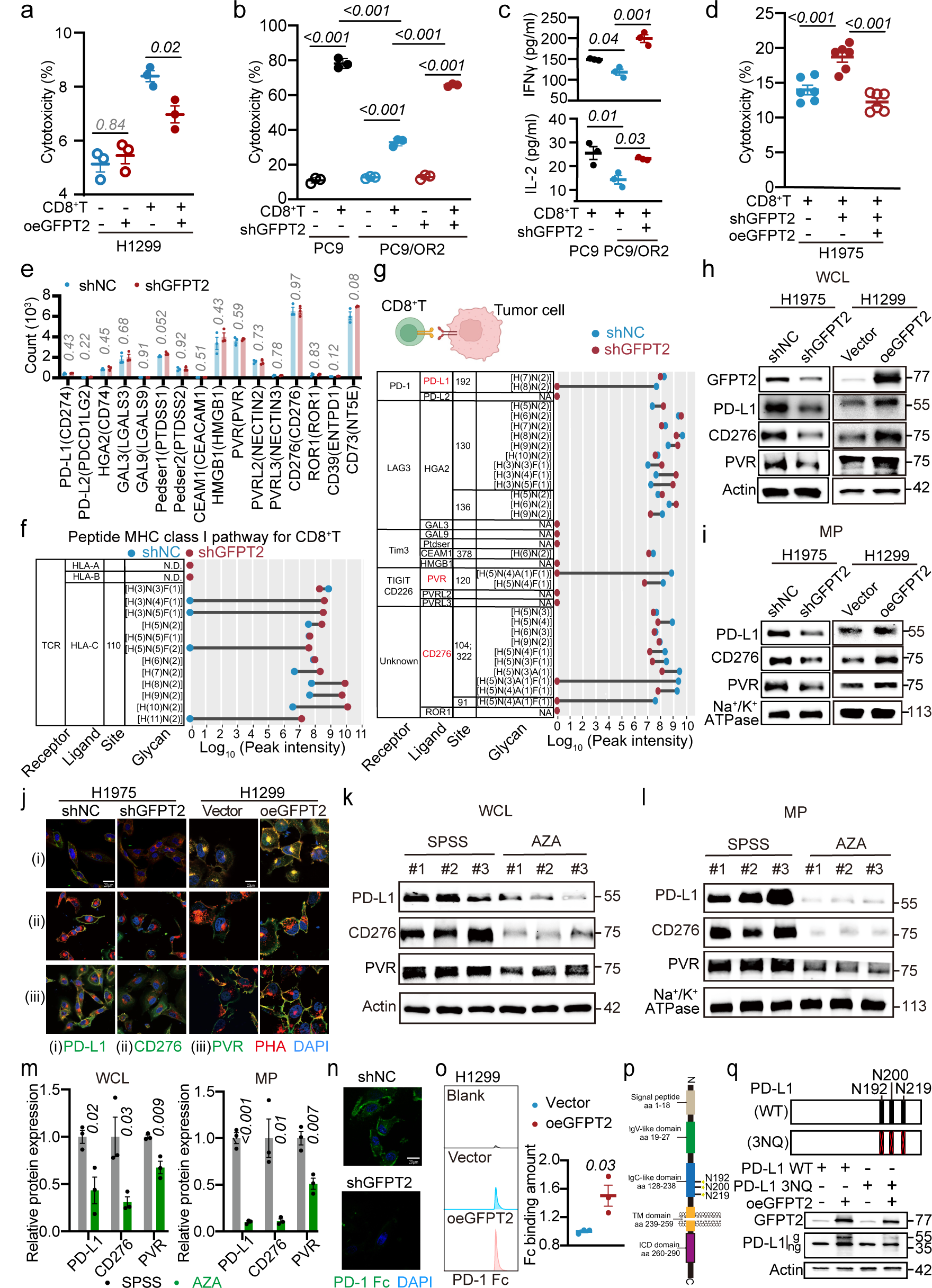
GFPT2 increases the N-glycosylation of immunosuppressive ligands including PD-L1, CD276 and PVR (Related to Fig.3) **a**, **b**, FCM analyses of H1299 and PC9/OR2 cell survival in co-culture with activated human CD8^+^T cell sorted from PBMC (n=3).**c**, IFNγ and IL-2 secretion in cell supernatant in indicated co-culture settings (n=3). **d**, FCM analyses of H1975 cell survival in co-culture with activated human CD8^+^T cell sorted from PBMC (n=3). **e**, Bulk-seq analysis of gene expression levels of various immunogenicity ligands in H1975 cells following GFPT2 knockdown (n=3). **f**, **g**, Analysis of N-linked glycans and peak intensities indicating glycosylation modifications on indicated immune checkpoint ligands and MHC I protein in shNC and shGFPT2 H1975 cells. **h-i**, Immunoblotting analysis of PD-L1, CD276, and PVR in whole cell lysate (WCL) and membrane protein (MP). Actin is used as a loading control for WCL, and Na^+^/K^+^ ATPase as a loading control for MP. **j**, Immunofluorescent images of H1975 cells with GFPT2 knockdown and H1299 cells with overexpressed GFPT2, with specific labeling for indicated immunosuppressive ligands and its associated glycosylation patterns. **k-m**, Immunoblotting examined the expression of total and membrane-associated proteins of indicated immunosuppressive ligands in H1975 xenograft tumors treated with AZA (n=3). Actin is used as a loading control for total protein, and Na^+^/K^+^ ATPase as a loading control for membrane protein. **n**, Immunofluorescence (IF) analysis assessing PD-1 Fc binding in GFPT2-silenced H1975 cells (representative images for triple independent experiments). **o**, FCM quantification of PD-1 Fc binding to PD-L1 on H1299 cell membranes upon GFPT2 overexpression, calculated by multiplying the percentage of positive cells by mean fluorescence intensity (n=3). **p**, Schematic representation of PD-L1 structure and glycosylation sites. **q**, Immunoblotting analysis of the effects of GFPT2 on the glycosylation modifications of wild-type PD-L1 and mutant PD-L1. g, glycosylated form; ng, non-glycosylated form. Data are mean ± s.e.m. **a**, **b**, **c**, **d**, One-way ANOVA. **e**, **m**, **o**, Two-tailed unpaired Student’s t-test.

**Extended Data Fig.7.**
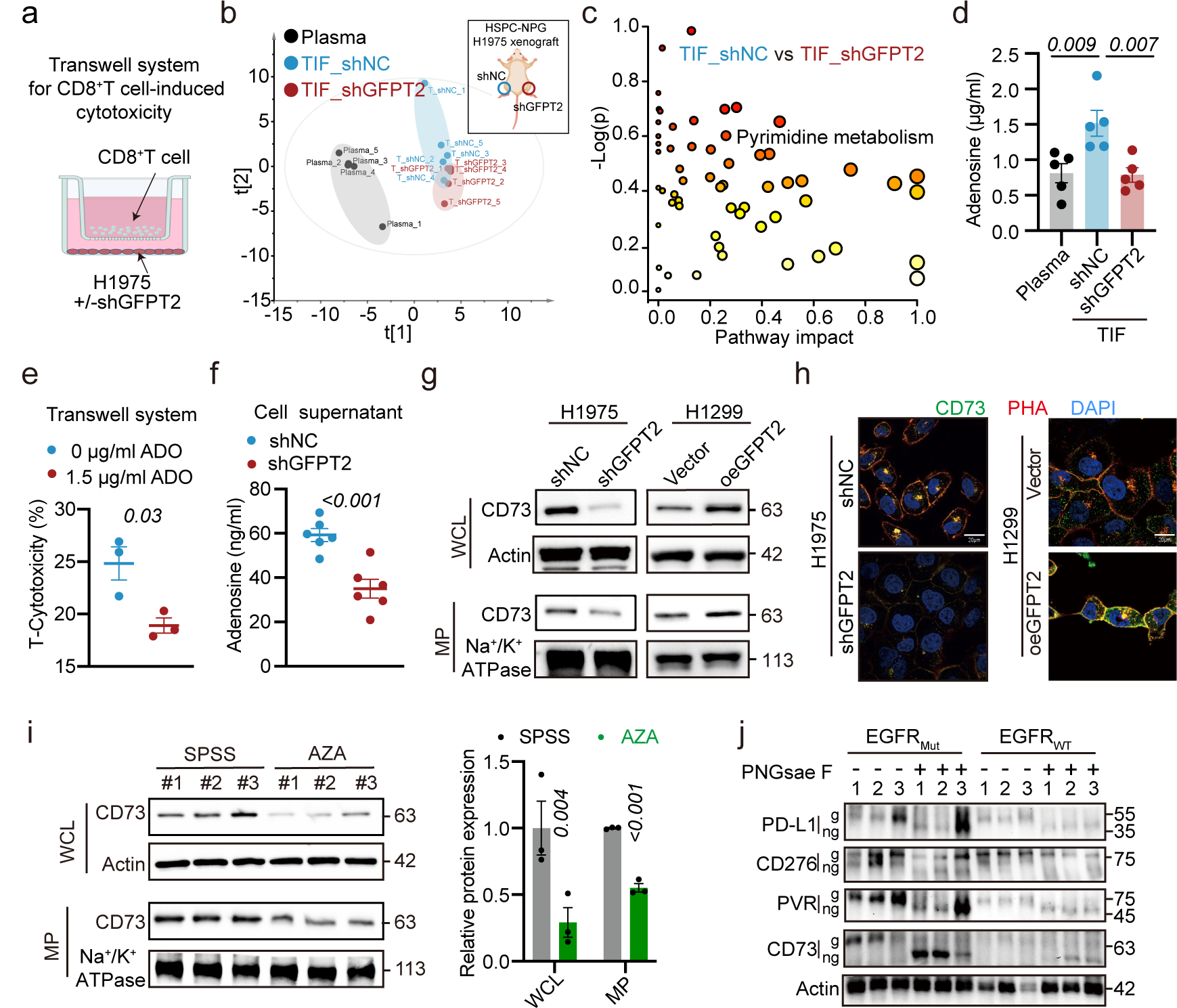
GFPT2 increases the N-glycosylation of CD73 (Related to Fig.3) **a**, Schematic of the Transwell co-culture system. **b-c**, Metabolomic analysis comparing TIF in shNC and shGFPT2 xenografts, assayed by HPLC-QTOF/MS (n=5). **(b)** PCA comparing the metabolites. **(c)** Metabolic pathway impact plot measuring the significance (-log(p)) and impact (proportion of pathway altered). **d**, HPLC-QTRAP/MS quantitative analysis of adenosine concentrations in plasma and TIF (n=5). **e**, FCM analysis of H1975 cell survival during co-culture with activated human CD8^+^T sorted from PBMC with or without adenosine (ADO) treatment (n=3). **f**, HPLC-QTRAP/MS quantification of changes in adenosine levels in the supernatant of GFPT2-knockdown H1975 cells (n=5). **g**, Immunoblotting analysis of total and membrane protein levels of CD73. Actin is used as a loading control for total protein, and Na^+^/K^+^ ATPase as a loading control for membrane protein. **h**, Immunofluorescent images of H1975 cells with GFPT2 knockdown and H1299 cells with overexpressed GFPT2, with specific labeling for CD73 and associated glycosylation patterns. **i**, Immunoblotting analysis examining the expression of total and membrane-associated proteins of indicated immunosuppressive ligands in H1975 xenograft tumors treated with AZA (n=3). Actin is used as a loading control for total protein, and Na^+^/K^+^ ATPase as a loading control for membrane protein. **j**, Immunoblotting for protein glycosylation of PD-L1, CD276, PVR, and CD73 in tumor tissues from NSCLC patients with EGFR wild-type and EGFR mutation (n=3). g, glycosylated form; ng, non-glycosylated form. Data are mean ± s.e.m. **e**, **f**, **i**, Two-tailed unpaired Student’s t-test. **d**, One-way ANOVA.

**Extended Data Fig.8.**
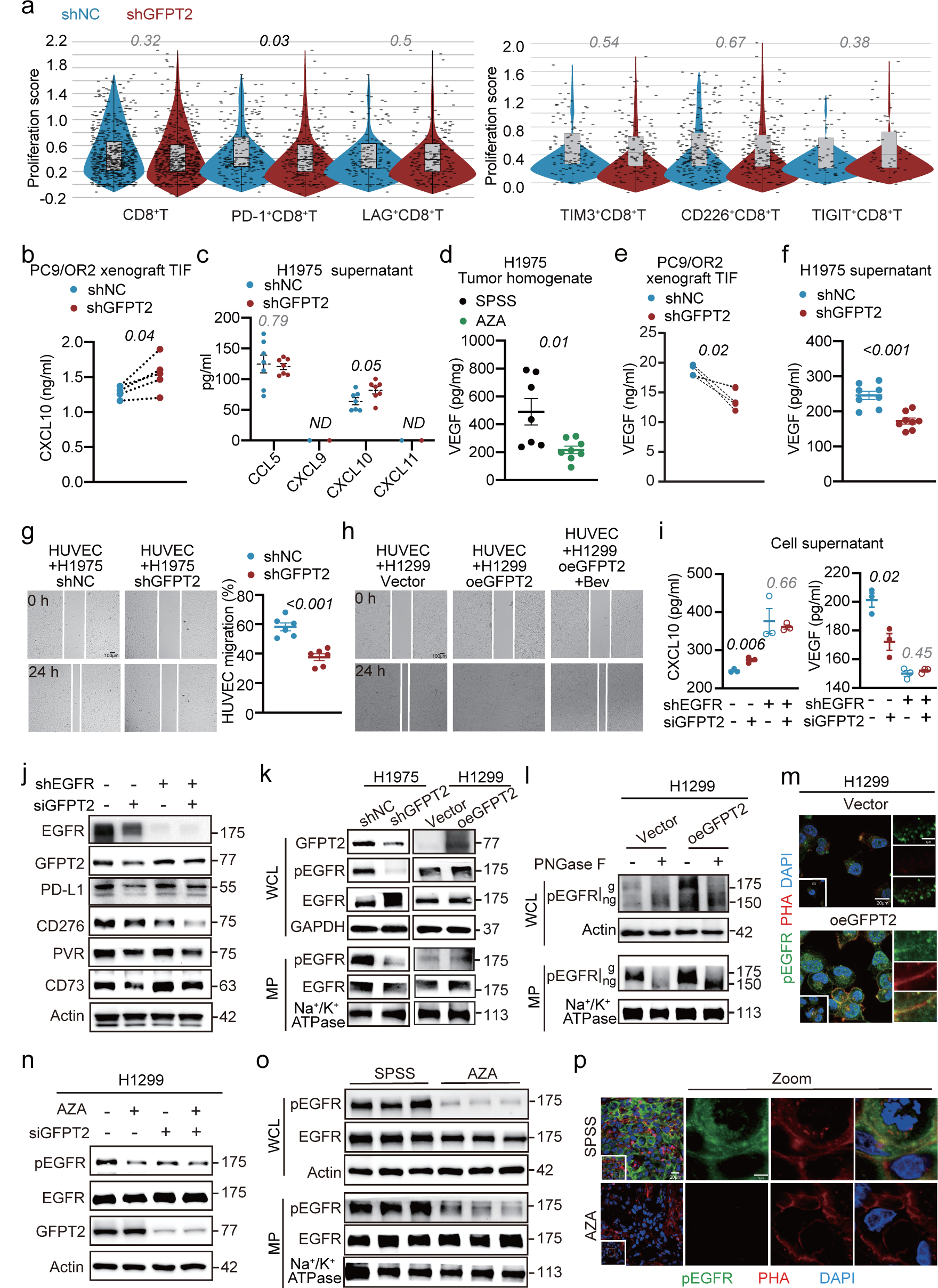
Impact of tumor cell GFPT2 on the secretion of CXCL10 and VEGF, and its effect on EGFR glycosylation (Related to Fig.4) **a**, scRNA-seq analysis of sorted CD8^+^ T cells from EGFR-mutated LLC xenograft. Violin plots display proliferation signaling in intratumoral CD8^+^T cells. Box plots indicate median and interquartile range. **b**, ELISA measurement of CXCL10 in the TIF of shNC and shGFPT2 PC9/OR2 xenografts in mice (n=5). **c**, ELISA quantification of various chemokines in the culture supernatant of H1975 cells with GFPT2 knockdown (n=6). **d-e**, ELISA assessments of VEGF levels in tumor homogenates and TIF following genetic intervention of tumor cell GFPT2 (n=5) or treatment with the chemical inhibitor AZA (n=7). **f**, ELISA measurement of VEGF in the culture supernatant of H1975 cells with GFPT2 knockdown (n=8). **g-h**, Transwell co-culture system used to examine the effects of knocking down (n=6) or overexpressing GFPT2 (n=6) on endothelial cell migration. **i**, ELISA determination of CXCL10 and VEGF levels secreted in the supernatant of H1975 cells after specific genetic interventions (n=3). **j**, Immunoblotting examination of the expression of three immunosuppressive ligands and CD73 in H1975 cells following the same genetic interventions. **k-l**, Immunoblotting analysis of the expression, activation, and glycosylation of EGFR in whole cell lysates and cell membrane fractions of H1975 cells with GFPT2 knockdown and H1299 cells with GFPT2 overexpression. **m**, IF investigation of membrane translocation and glycosylation levels of pEGFR in H1299 cells after GFPT2 overexpression. **n**, Immunoblotting examination of whether GFPT2 knockdown affects the AZA intervention on pEGFR. **o**, Immunoblotting analysis examining the expression of total and membrane-associated proteins of indicated immunosuppressive ligands in H1975 xenograft tumors treated with AZA (n=3). Actin is used as a loading control for total protein, and Na^+^/K^+^ ATPase as a loading control for membrane protein. **p**, Immunofluorescent images of AZA-treated H1975 xenografts, with specific labeling for pEGFR and its associated glycosylation patterns. Data are mean ± s.e.m. except in **b**, **e**, Two-tailed paired Student’s t-test. **a**, **c**, **d**, **f**, **g**, Two-tailed unpaired Student’s t-test. **i**, One-way ANOVA.

**Extended Data Fig.9.**
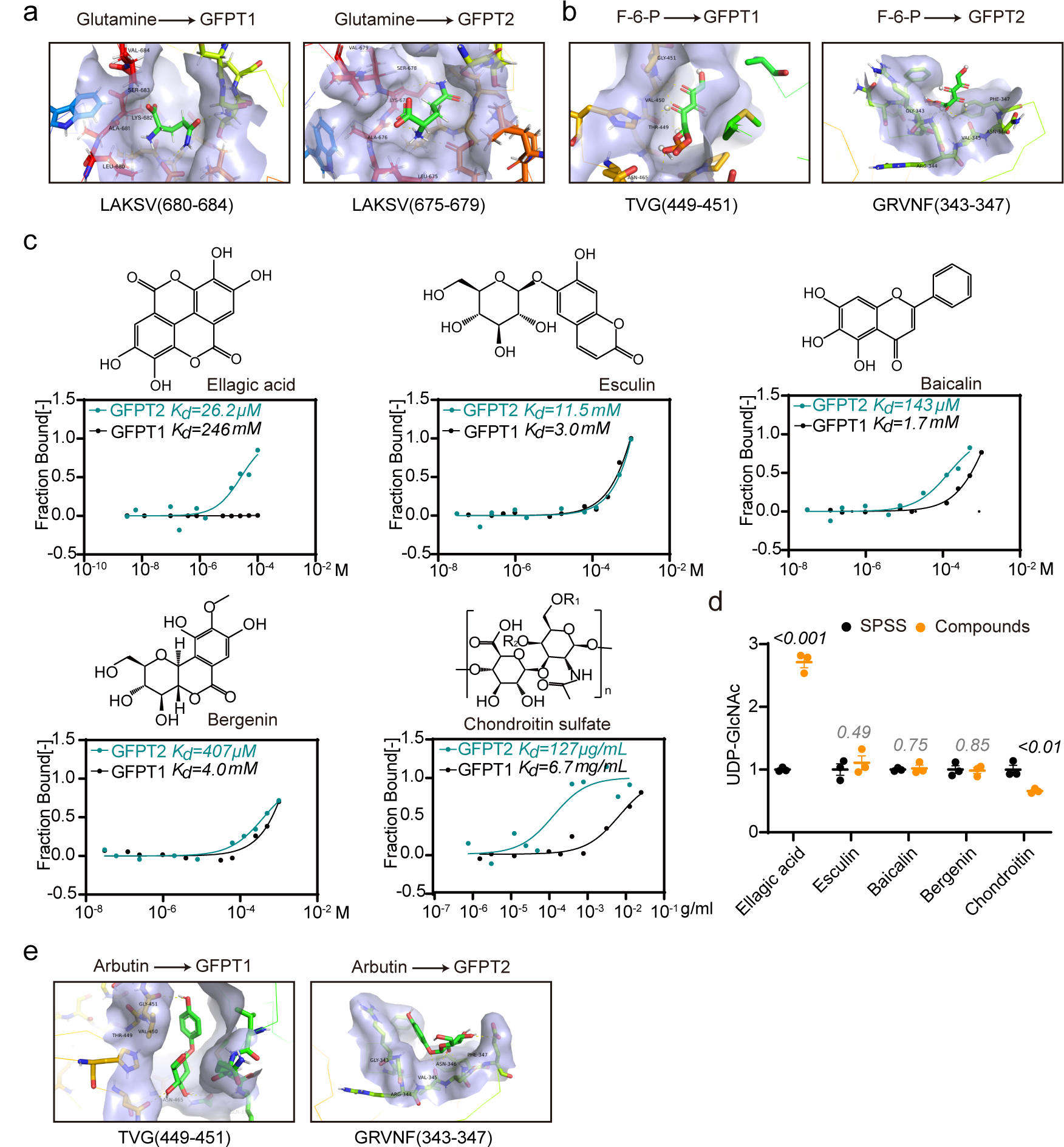
Screening for GFPT2 isoform-specific inhibitor (Related to Fig.5) **a-b**, Binding patterns of glutamine and fructose-6-phosphate with GFPT1 and GFPT2. **c**, Structural formulas of other five candidate hits with their affinities to GFPT1 and GFPT2 assessed by Microscale Thermophoresis (MST) and calculation of dissociation constants (Kd values). **d**, Effects of other five candidate hits on intracellular UDP-GlcNAc production (n=3). **e**, Binding patterns of Arbutin with GFPT1 and GFPT2. Data are mean ± s.e.m. **d**, Two-tailed unpaired Student’s t-test.

**Extended Data Fig.10.**
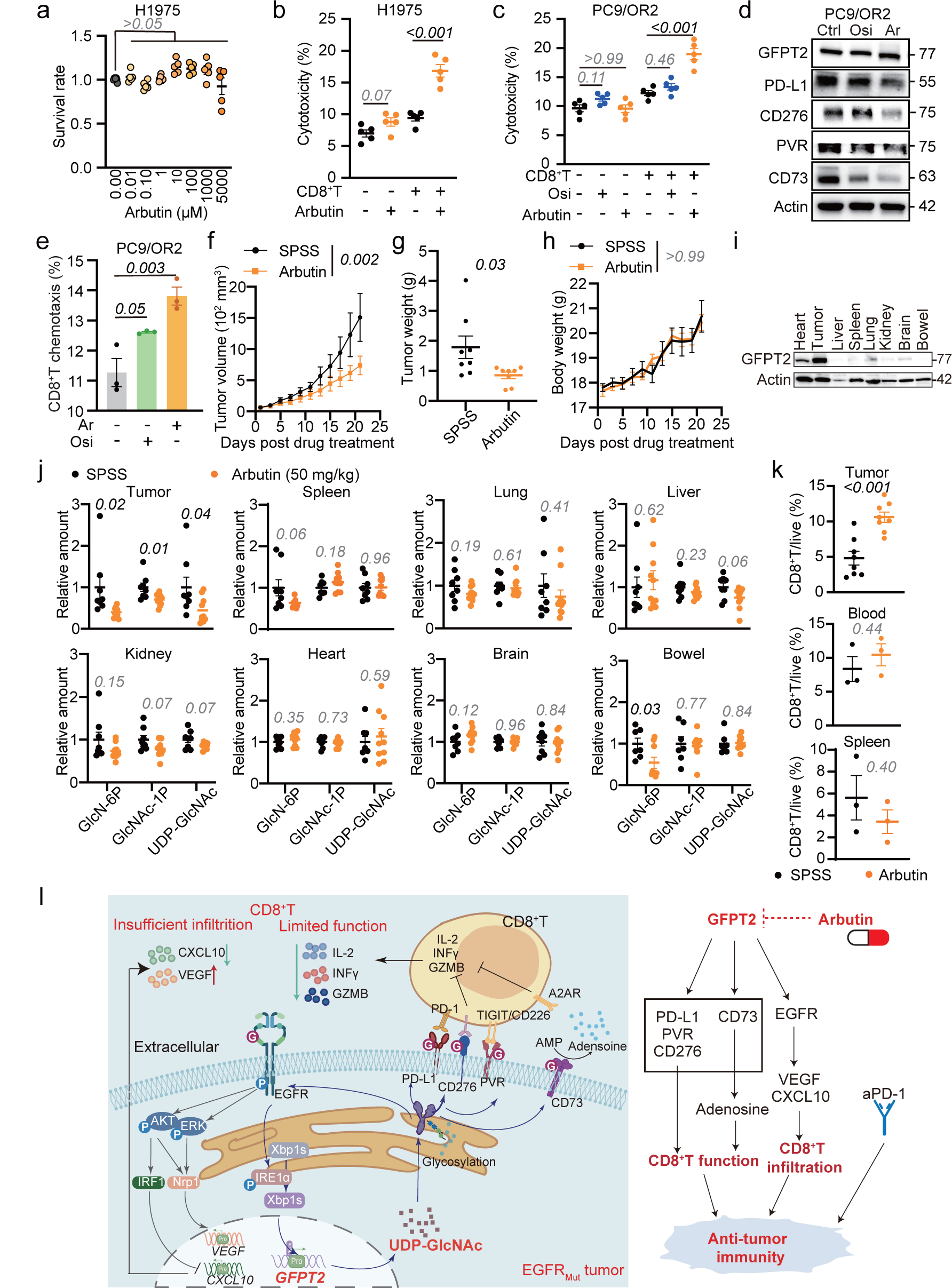
Arbutin enhances the efficacy of immunotherapy (Related to Fig.5) **a**, CCK-8 assay evaluating the effect of different concentrations of Arbutin on the viability of H1975 cells (n=5). **b-c**, FCM analyses of H1975 and PC9/OR2 cell survival in co-culture with activated human CD8^+^T cells sorted from PBMC (n=5). **d**, Western blot analysis of specified protein levels in PC9/OR2 cells following treatment with Osimertinib (Osi) and Arbutin (Ar). **e**, PC9/OR2 cells and CD8^+^T cells were co-cultured using a Transwell system to assess the effects of Osi and Ar on CD8^+^T cell chemotaxis (n=3). **f-h**, Changes in tumor size, tumor weight, and body weight of EGFR-mutated LLC xenograft in C57BL/6N following Arbutin treatment (n=8). **i**, Immunoblotting for GFPT2 in tumor tissues and major organs with Actin serving as the loading control. **j**, HPLC-QTRAP/MS analysis of HBP metabolites in tumor tissues and major organs after Arbutin treatment (n=8). **k**, FCM analysis of changes in CD8^+^T cell infiltration within the tumor (n=8), as well as peripheral blood and spleen, following Arbutin administration (n=3). **l**, Schematic representation of how GFPT2 targeting enhances immunotherapy in EGFR-mutated NSCLC via metabolite-mediated modulation of protein glycosylation. Data are mean ± s.e.m. **g**, **j**, **k**, Two-tailed unpaired Student’s t-test. **a**, **b**, **c**, **e**, One-way ANOVA. **f**, **h**, Two-way ANOVA.

## Notes

### Competing Interest Statement

The authors have declared no competing interest.

