## Supplementary Figures for "Targeting GFPT2 to reinvigorate immunotherapy in EGFR-mutated NSCLC"

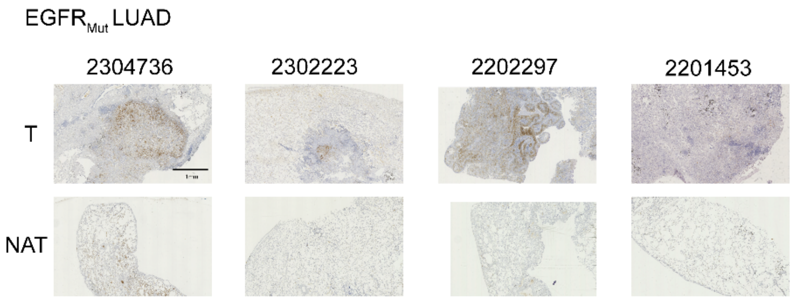


**Supplementary Fig 1. IHC images for GFPT2 staining of T and NAT from 4 $EGFR_{Mut}$ LUAD patients.**

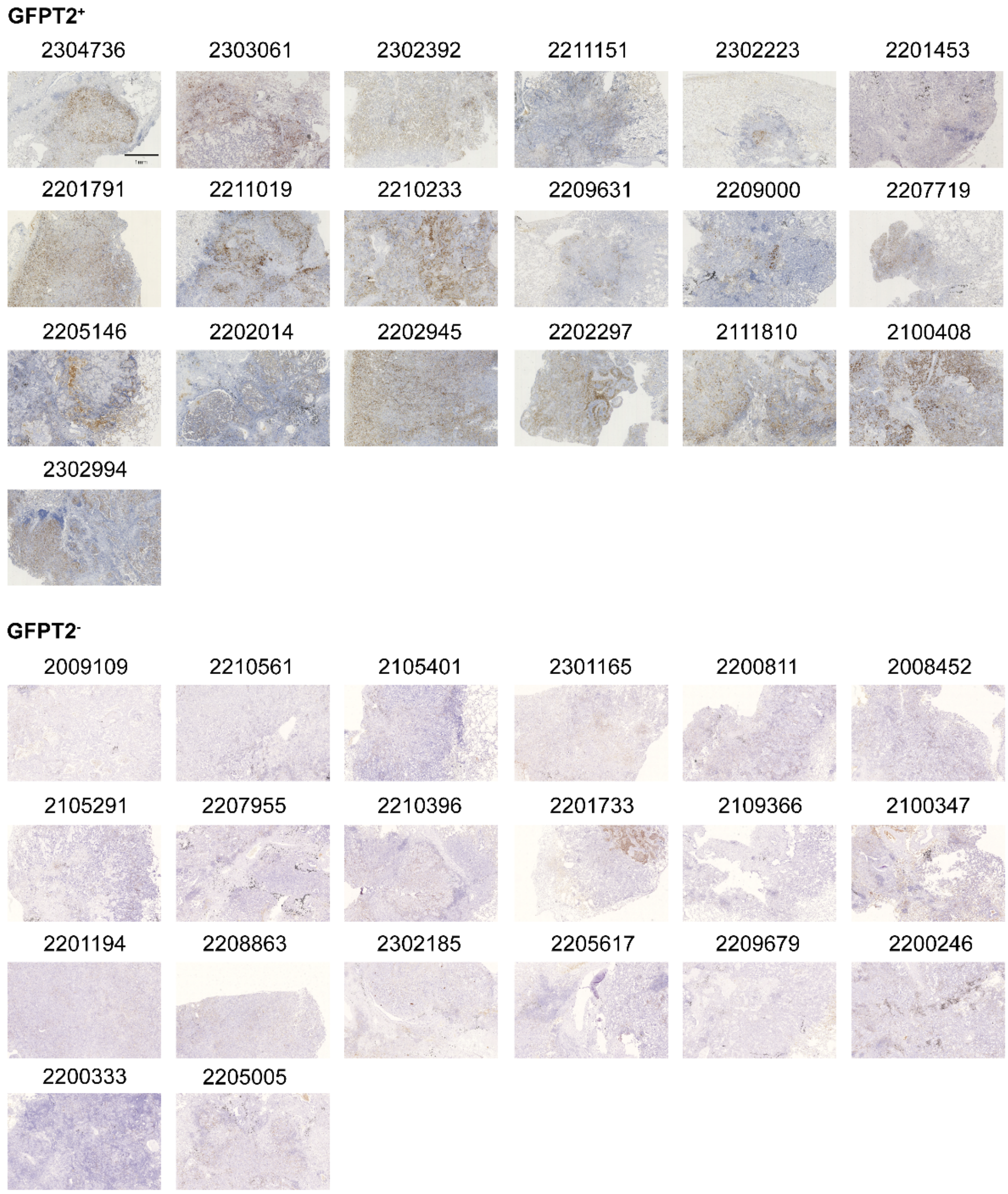


**Supplementary Fig 2. IHC images for GFPT2 staining of 19 paired LUAD T and NAT.**

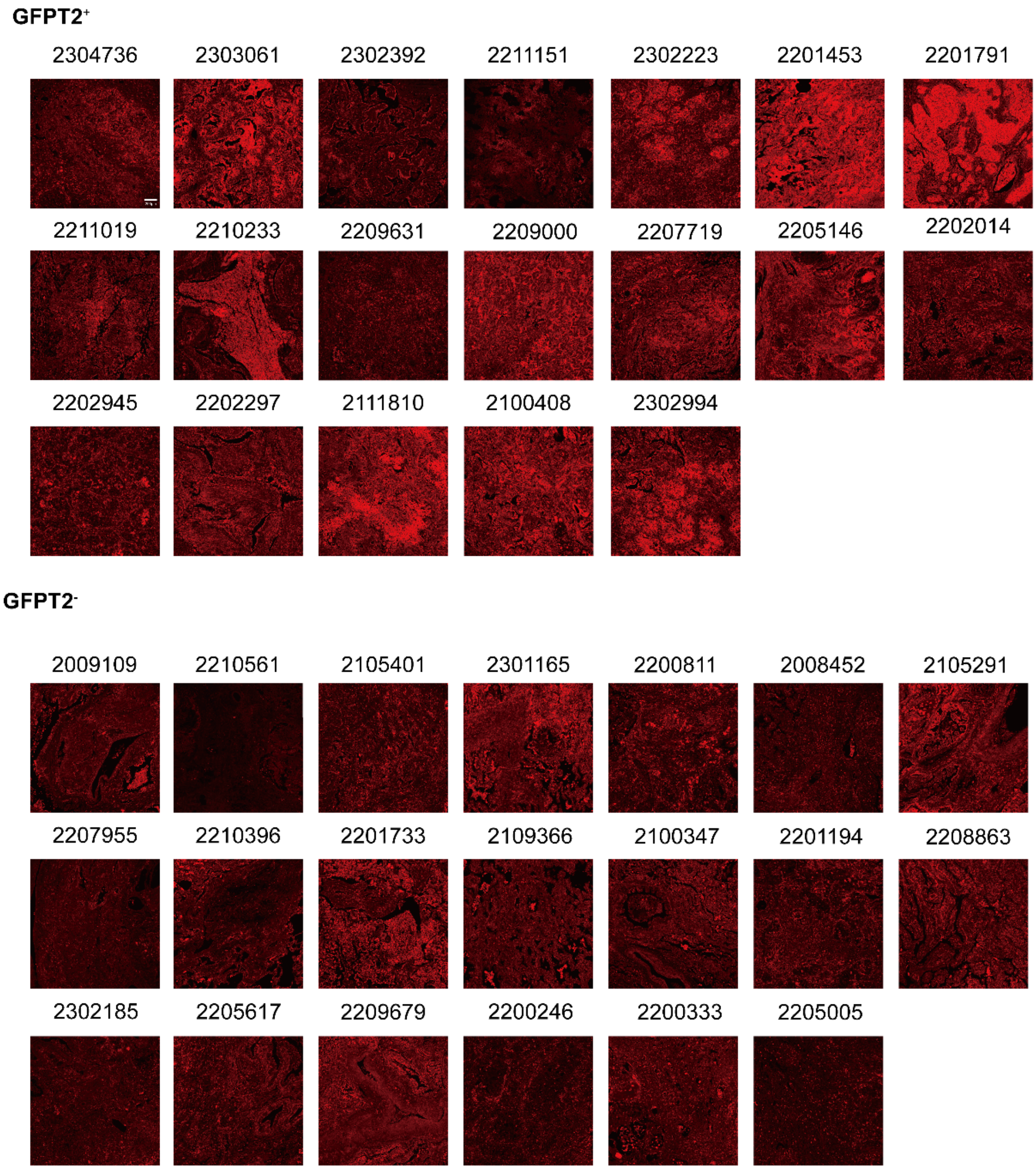


**Supplementary Fig 3. IF images for PHA staining of 19 GFPT2 positive expressed LUAD T and 20 GFPT2 negative expressed LUAD T.**

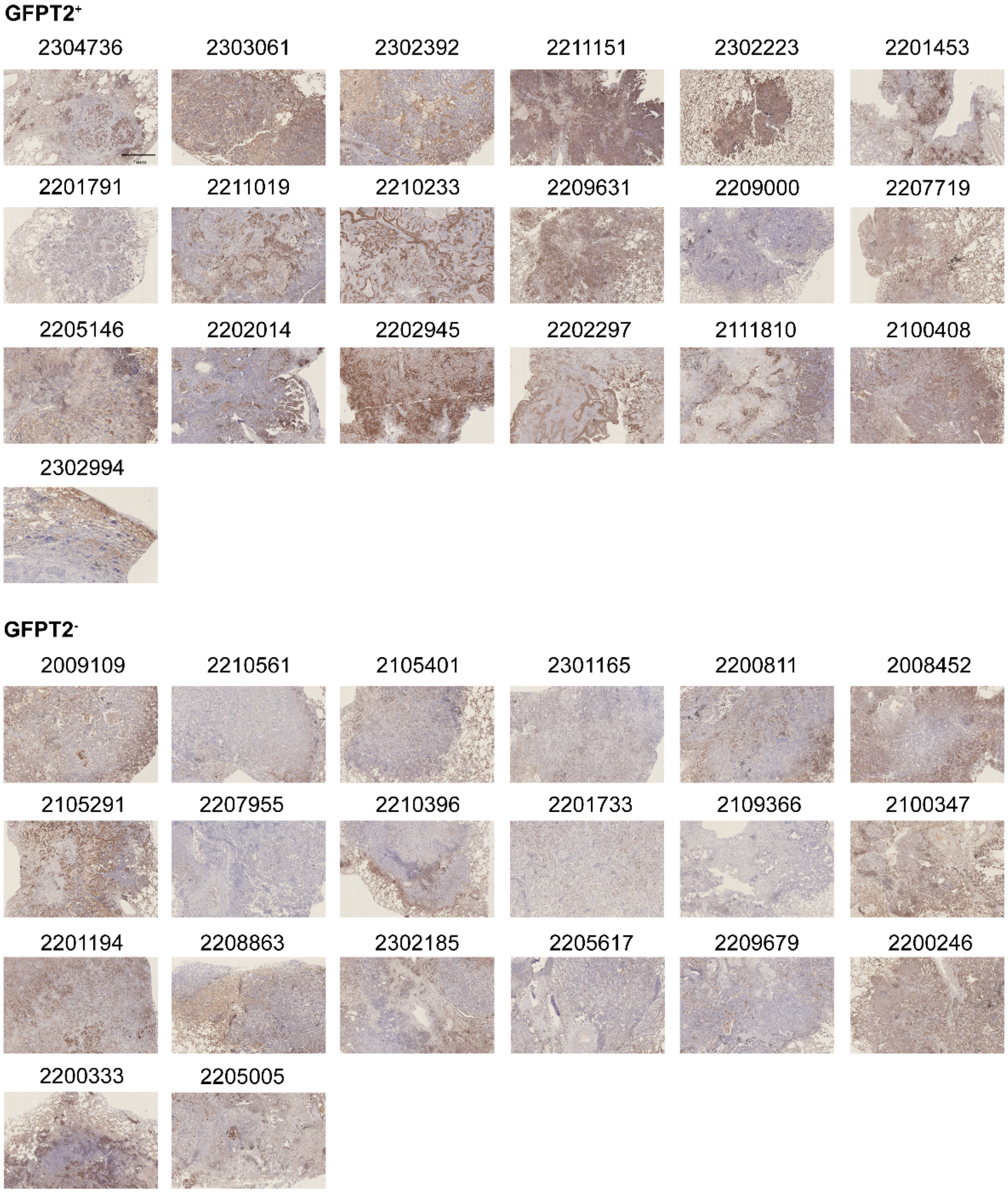


**Supplementary Fig 4. IHC images for pEGFR staining of 19 GFPT2 positive expressed LUAD T and 20 GFPT2 negative expressed LUAD T.**

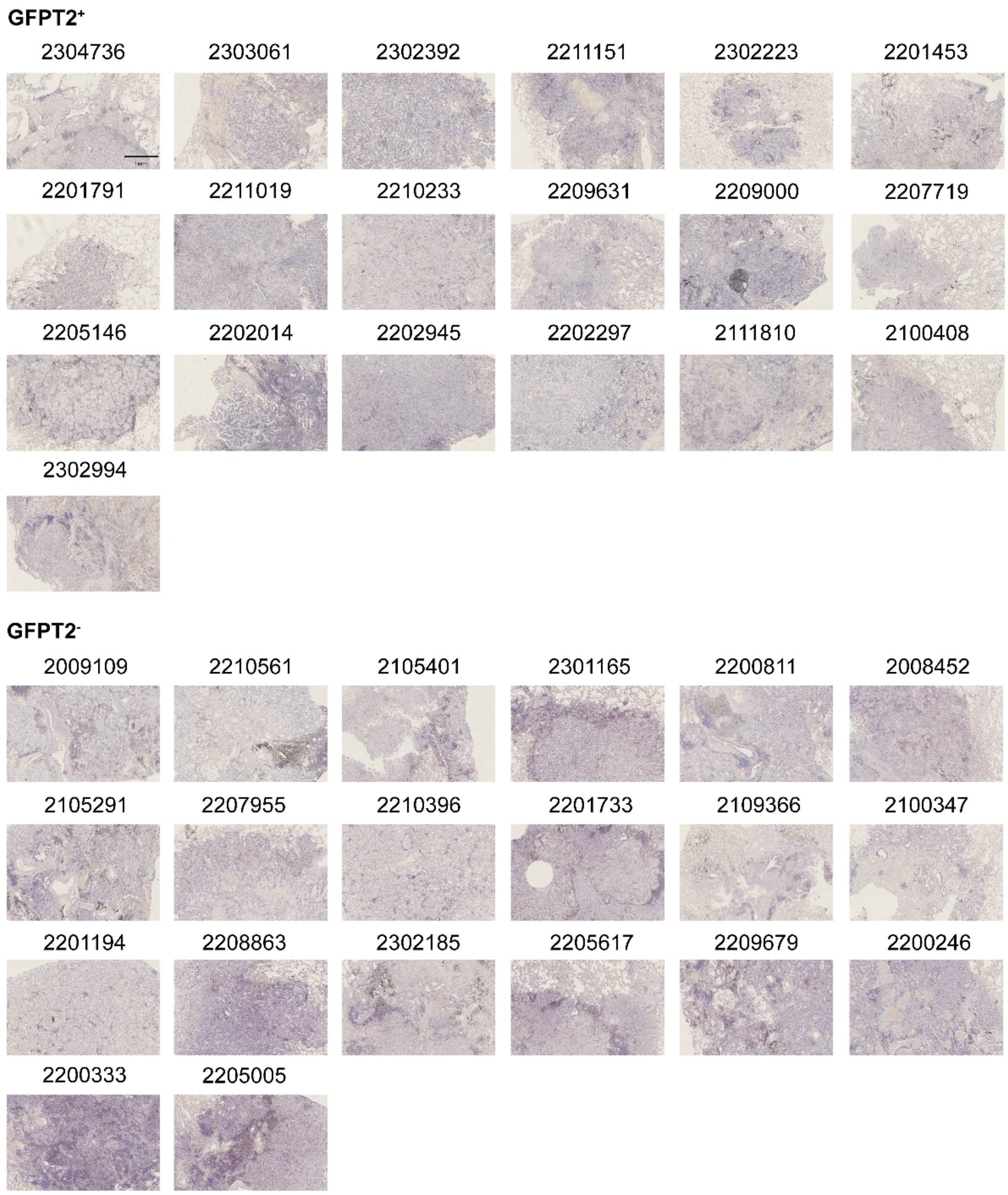


**Supplementary Fig 5. IHC images for CD8T$^{+}$T staining of 19 GFPT2 positive expressed LUAD T and 20 GFPT2 negative expressed LUAD T.**

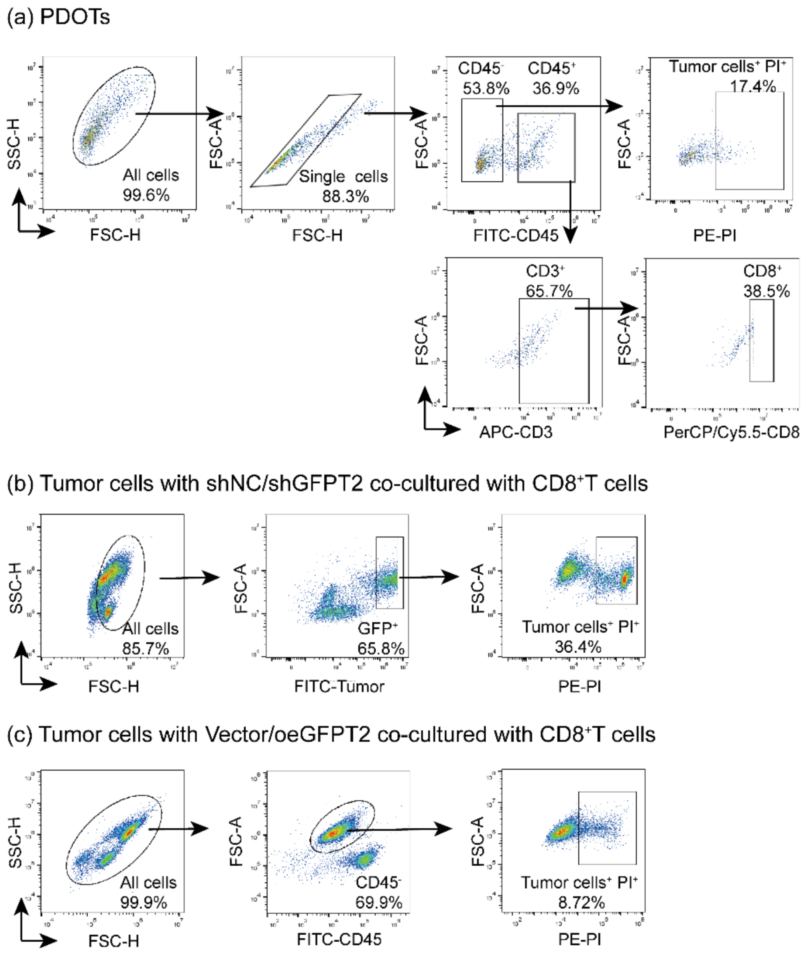


**Supplementary Fig 6. Flow cytometry gating strategy for ex vivo co-culture and PDOT.**

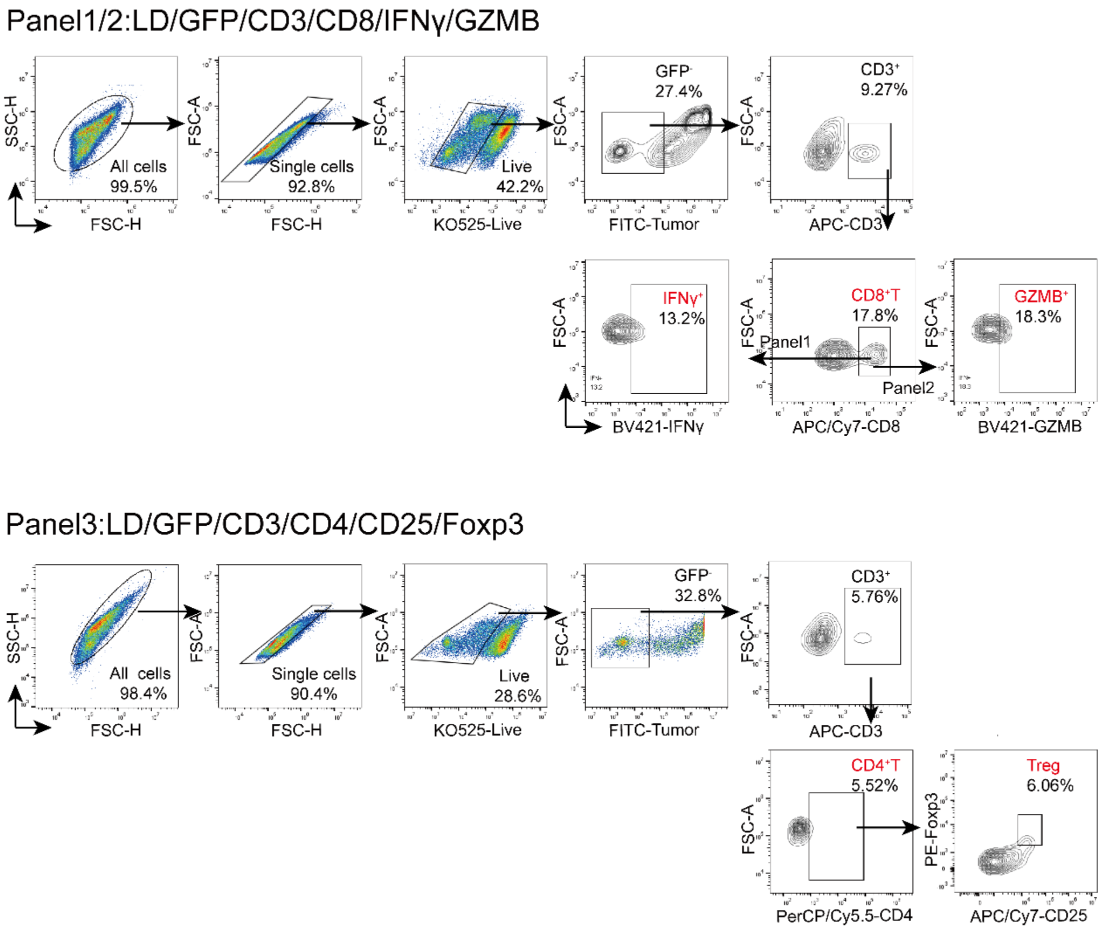


**Supplementary Fig 7. Flow cytometry gating strategy for tumor immune microenvironment (TIME) in humanized mouse models.**

Panel1:LD/CD45/CD3/CD8/IFNγ/GZMB

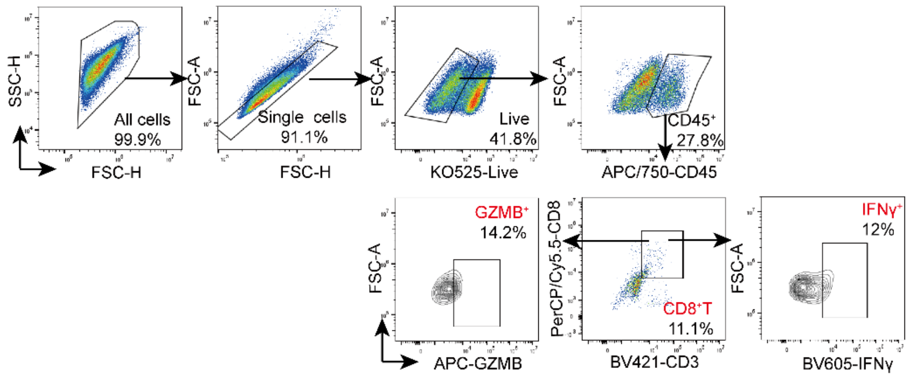


Panel2:LD/CD45/CD3/CD4/CD25/Foxp3

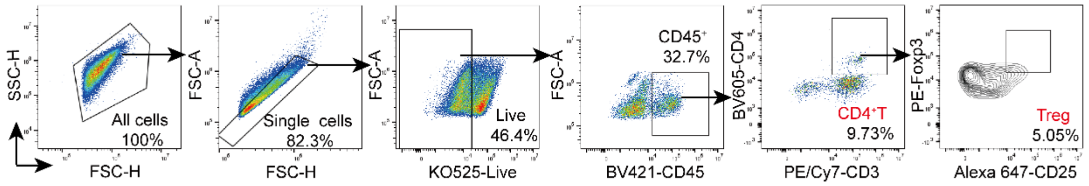


Panel3:LD/CD45/F4/80/CD11b/CD86/CD206

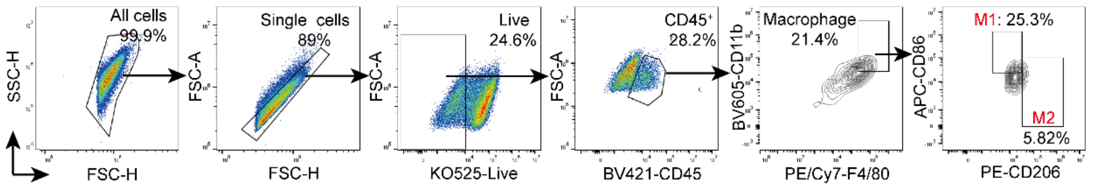


Panel4:LD/CD45/CD3/CD19/NK1.1

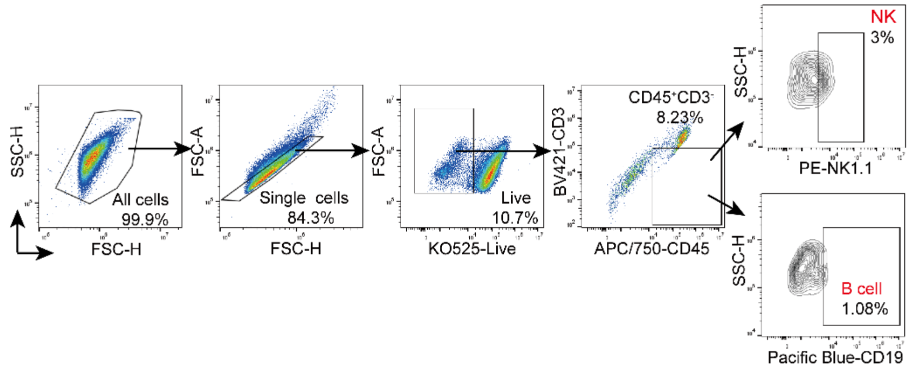


**Supplementary Fig 8. Flow cytometry gating strategy for the tumor immune microenvironment (TIME) in fully immunocompetent mice.**

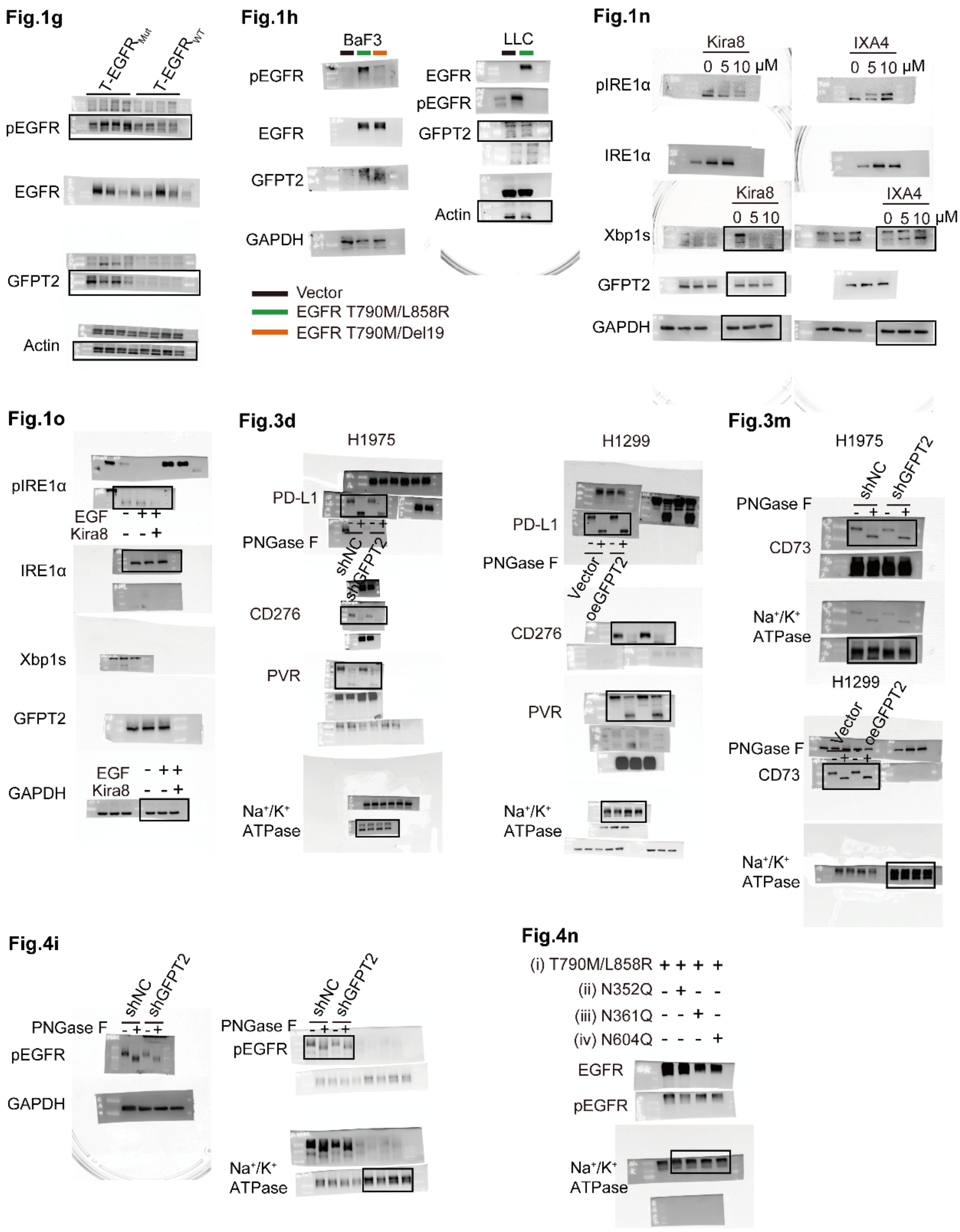


**Supplementary Fig 9. Original immunoblotting data related to main figures.**

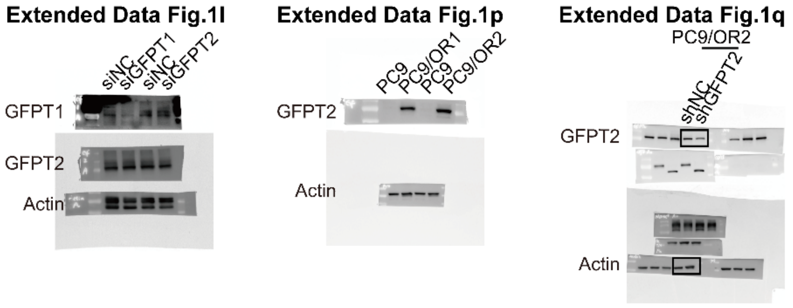


**Supplementary Fig 10. Original immunoblotting data related to Extended Data Fig1.**

Extended Data Fig.2a

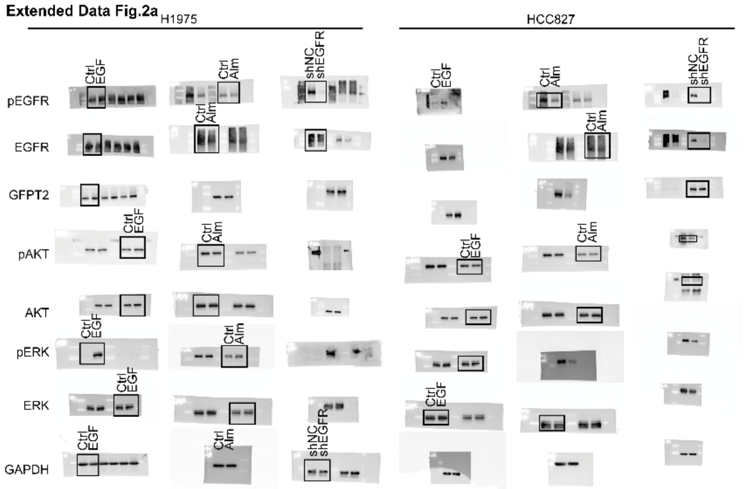


Extended Data Fig.2c

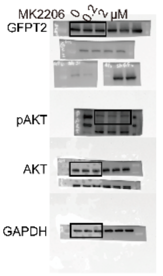


Extended Data Fig.2g

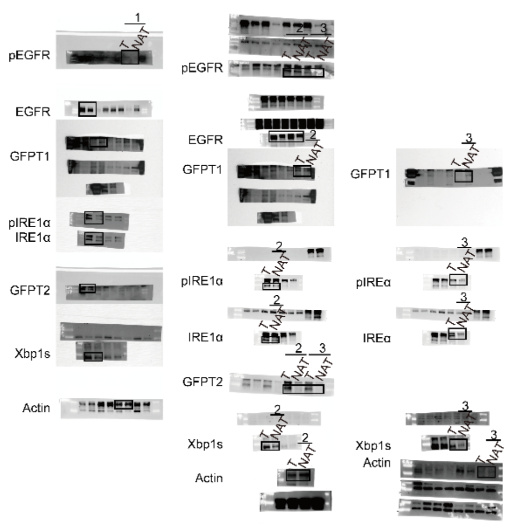


Extended Data Fig.2d

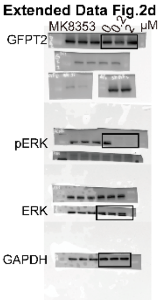


Extended Data Fig.2h

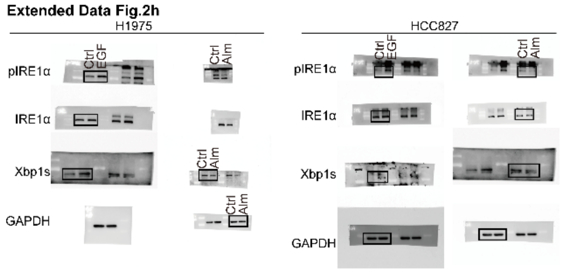


Extended Data Fig.2i

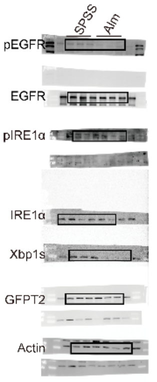


Extended Data Fig.2k

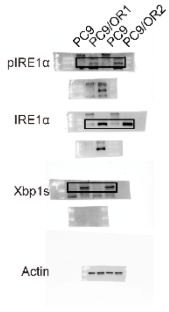

**Supplementary Fig 11. Original immunoblotting data related to Extended Data Fig2.**

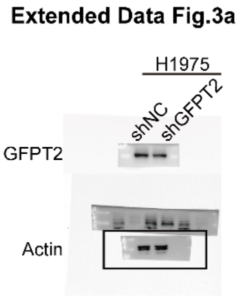


**Supplementary Fig 12. Original immunoblotting data related to Extended Data Fig3.**

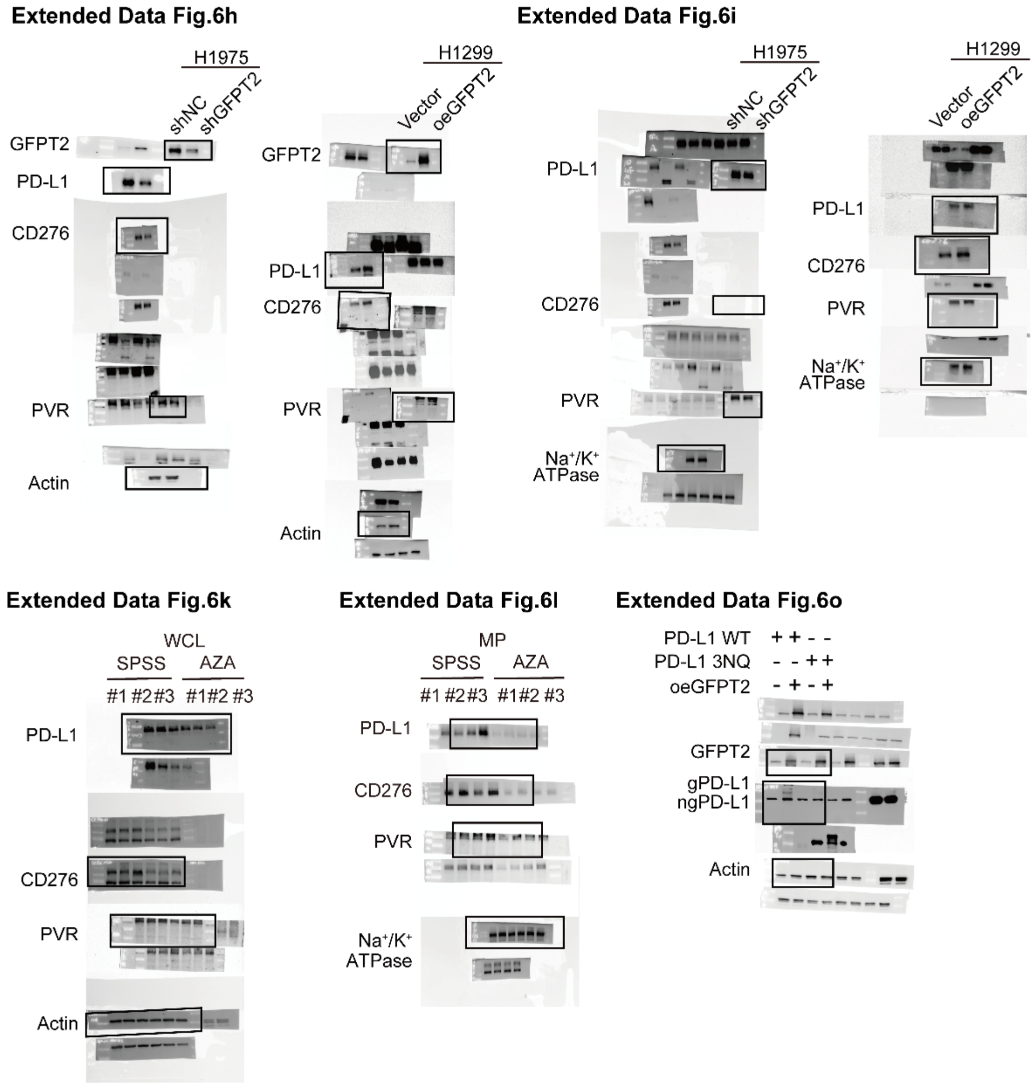


**Supplementary Fig 13. Original immunoblotting data related to Extended Data Fig6.**

**Extended Data Fig.7g**

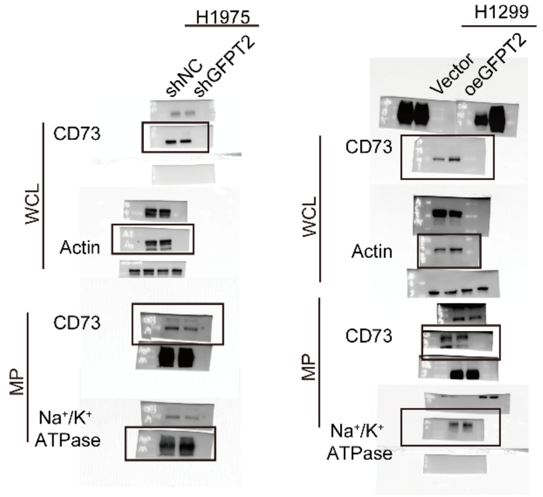


**Extended Data Fig.7i**

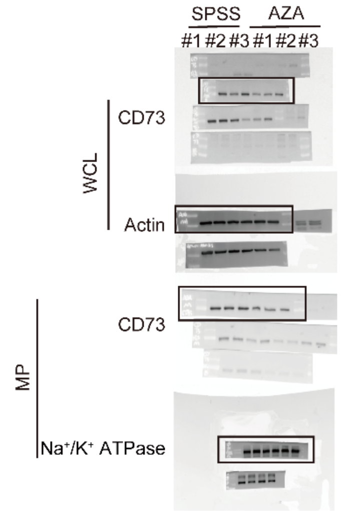


**Extended Data Fig.7j**

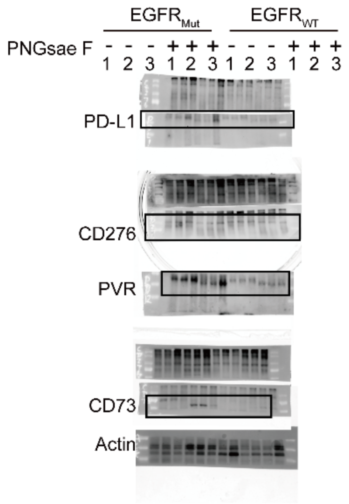


**Supplementary Fig 14. Original immunoblotting data related to Extended Data Fig7.**

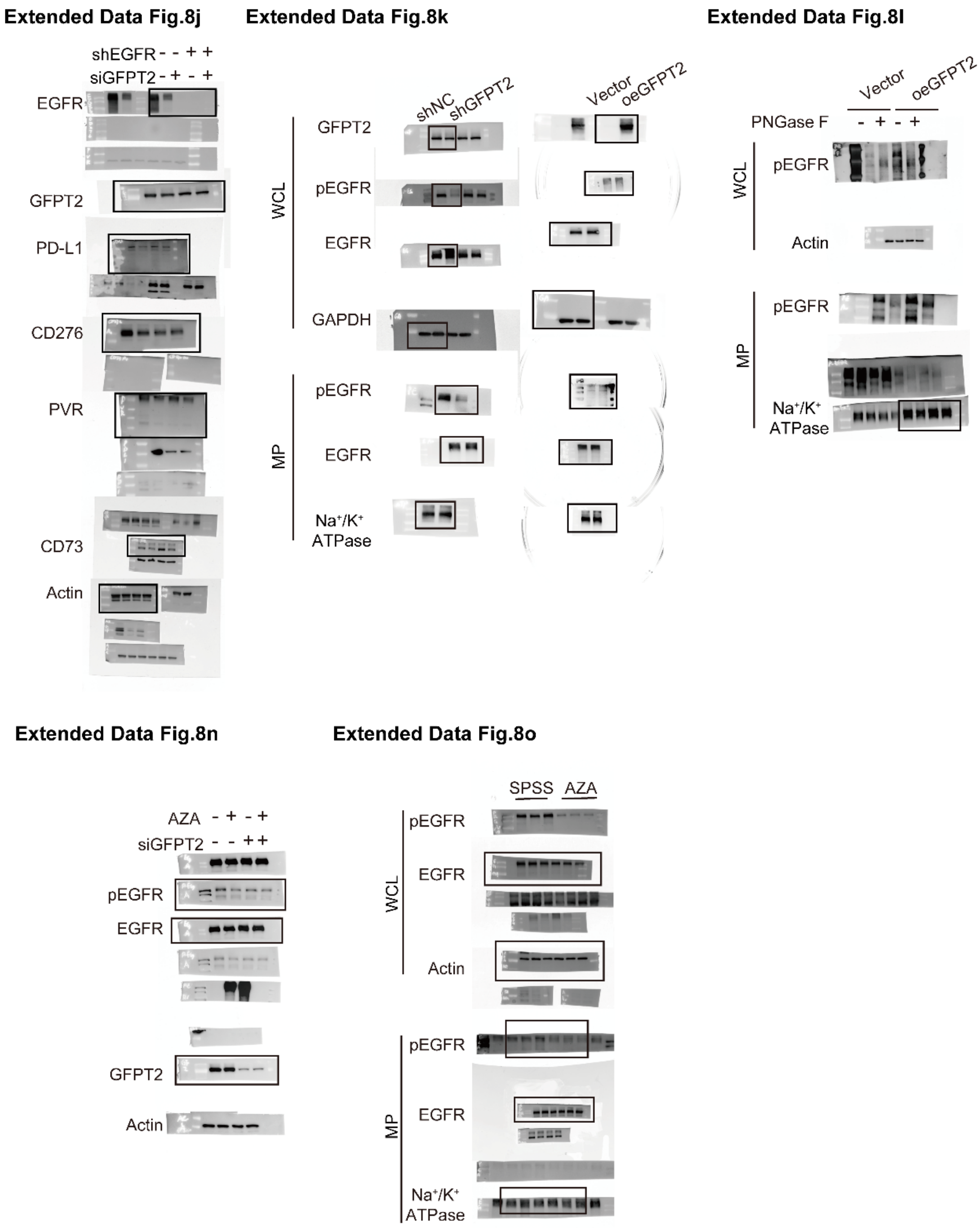


**Supplementary Fig 15. Original immunoblotting data related to Extended Data Fig8.**

**Extended Data Fig.10d**

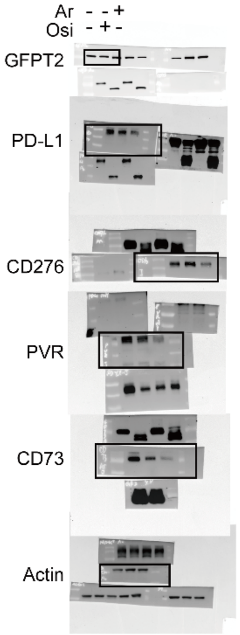


**Extended Data Fig.10i**

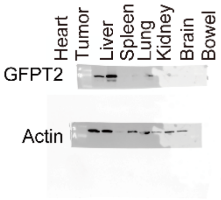


**Supplementary Fig 16. Original immunoblotting data related to Extended Data fig10.**
